# Aminoacyl-tRNA specificity of a ligase catalyzing non-ribosomal peptide extension

**DOI:** 10.1101/2025.09.13.676032

**Authors:** Dinh T. Nguyen, Josseline S. Ramos-Figueroa, Alexander A. Vinogradov, Yuki Goto, Mayuresh G. Gadgil, Rebecca A. Splain, Hiroaki Suga, Wilfred A. van der Donk, Douglas A. Mitchell

**Author notes:** **Corresponding Authors** Wilfred A. van der Donk − Department of Chemistry and Howard Hughes Medical Institute, University of Illinois at Urbana-Champaign, Urbana, IL, United States;,., Douglas A. Mitchell − Departments of Biochemistry and Chemistry, Vanderbilt University School of Medicine, Nashville, TN, United States.

## Abstract

Peptide aminoacyl-transfer ribonucleic acid ligases (PEARLs) are amide bond-forming enzymes that extend the main chain of peptides using aminoacyl-tRNA (aa-tRNA) as a substrate. In this study, we investigated the substrate specificity of the PEARL BhaB_C_^Ala^ from *Bacillus halodurans*, which utilizes Ala-tRNA^Ala^. By leveraging flexizyme, a ribozyme capable of charging diverse acids onto a desired tRNA, we generated an array of aa-tRNAs in which we varied both the amino acid and the tRNA to dissect the substrate scope of BhaB_C_^Ala^. We demonstrate that BhaB_C_^Ala^ catalyzes peptide extension with non-cognate proteinogenic and non-canonical amino acids, hydroxy acids, and mercaptocarboxylic acids when attached to tRNA^Ala^. For most of these, the efficiency was considerably reduced compared to Ala, indicating the enzyme recognizes the amino acid. By varying the different parts of the tRNA, enzyme specificity was shown to also depend on the acceptor stem and the anticodon arm of the tRNA. These findings establish the molecular determinants of PEARL specificity and provide a foundation for engineering these enzymes for broader applications in peptide synthesis.

---

Amide bond formation is a critical process for the preparation of peptide and protein-based therapeutics.^1^ Accessing these structures usually relies on solid-phase peptide synthesis or ribosomal translation.^2-4^ The recent increased interest in using biocatalysis presents opportunities to incorporate enzymes that form amide bonds into synthetic processes.^5-12^ Enzyme-based synthesis potentially offers scalability with high efficiency, selectivity, and waste minimization.^13-16^ One class of amide-bond-forming enzymes are the peptide aminoacyl-transfer ribonucleic acid ligases (PEARLs).^17-21^ PEARLs utilize aminoacyl-transfer ribonucleic acid (aa-tRNA) to form a new peptide bond at the C-terminus of a peptide (Figure 1A).^17,22^ Whereas the specificity for the peptide substrate has been investigated,^23^ the factors that determine the specificity for the aminoacyl-tRNA substrate are currently unresolved. Here we investigate this specificity for a representative PEARL from *Bacillus halodurans*, BhaB_C_^Ala^ (NCBI accession identifier: BAB05753.1),^18^ which naturally utilizes Ala-tRNA^Ala^. Our data show that the combination of amino acid and cognate tRNA ensures fidelity and that the anticodon arm and the acceptor stem predominantly determine the tRNA specificity of BhaB_C_^Ala^. The enzyme also recognizes the charged amino acid (i.e., attached to the tRNA), but exhibits some tolerance that may be explored for engineering purposes.

**Figure 1.**
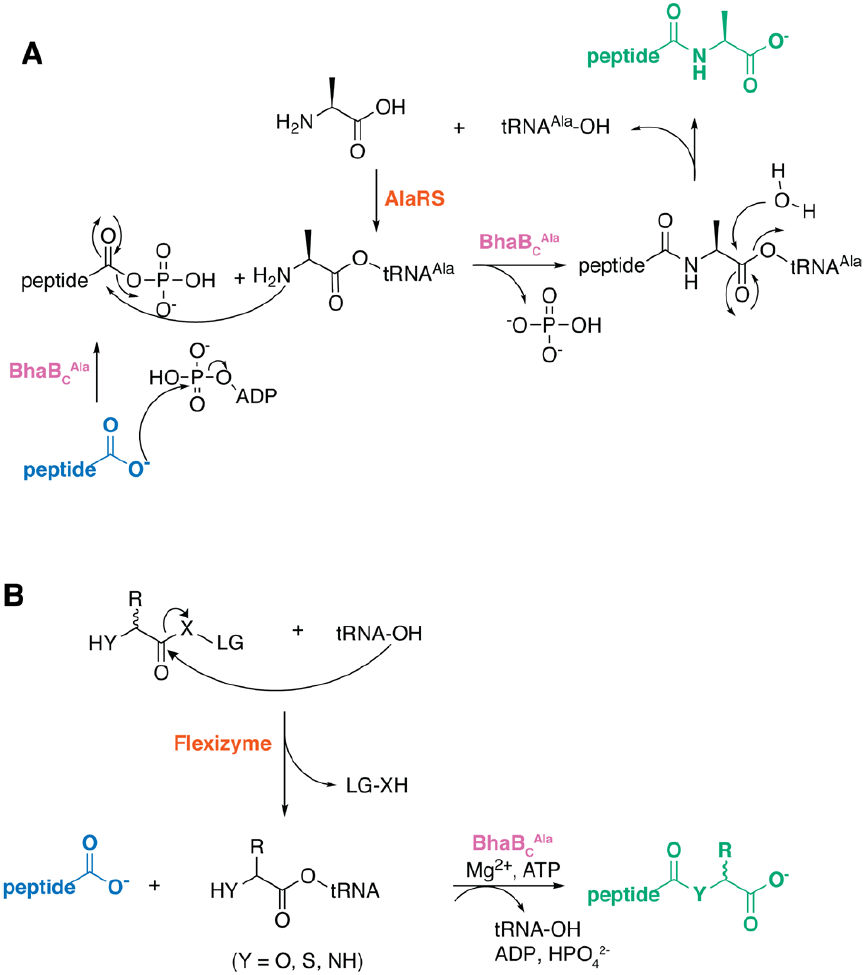
A) The proposed mechanisms of amide-bond formation catalyzed by PEARLs,^22^ in this case BhaB_C_^Ala^.^18^ Here, the co-substrate Ala-tRNA^Ala^ is formed by AlaRS. C) Examination of the substrate specificity of PEARLs using diverse aa-tRNA chimera generated by flexizyme, the focus of this work. X-LG here represents 4-chlorobenzyl thioester, 3,5-dinitrobenzylester, or cyanomethyl ester, which are leaving groups recognized by flexizyme (details in Supporting Information and Table S4). Blue highlights the substrate peptide, green highlights the product, purple highlights the PEARL enzyme, and orange highlights the means used to generate the co-substrate aminoacyl-tRNAs.

In previous studies on PEARLs, where product formation was assessed through both *in vitro* assays and *in vivo* co-expression in *Escherichia coli*, the aa-tRNA was generated using aminoacyl-tRNA synthetases (aaRS). Because aaRSs are highly specific toward their cognate aa-tRNAs, often containing editing domains that hydrolyze mischarged tRNAs,^24^ further exploration of the aa-tRNA specificity using aaRSs is limited to alanine and natural tRNA isoforms. Therefore, we first investigated whether both isoforms of tRNA^Ala^ in *E. coli* (GGC and UGC anticodons) were competent substrates for BhaB_C_^Ala^-catalyzed addition of Ala to its substrate peptide BhaA (NCBI: WP_010898193.1) *in vitro*. Analysis by matrix-assisted laser desorption/ionization time-of-flight mass spectrometry indicated that both isoacceptors were substrates (Figure S1). The limitation of aaRSs to investigate substrate scope can be overcome using flexizymes, 45-or 46-nucleotide ribozymes that catalyze the formation of aminoacyl-tRNAs from diverse activated amino acids and tRNA sequences (Figure 1B).^25-32^ Using flexizymes, we investigated whether BhaB_C_^Ala^ can extend the substrate peptide with various acids attached to tRNA^Ala^. We also investigated the tolerance of BhaB_C_^Ala^ toward alanine attached to various non-cognate tRNA sequences.

We first conducted BhaB_C_^Ala^ reactions using Ala-tRNA^Ala^(UGC) generated by flexizyme instead of AlaRS. The flexizyme reaction products were partially purified by ethanol precipitation, and the concentrations of aa-tRNA were estimated using a previously reported NaIO_4_-RNA extension assay (Figure S2).^33-35^ The reaction efficiency of BhaB_C_^Ala^ at different aa-tRNA concentrations was analyzed using liquid chromatography-electrospray ionization mass spectrometry (LC-ESI-MS). For all experiments, we used a truncated version of BhaA lacking the first 19 N-terminal residues (Δ19BhaA), which underwent quantitative modification by BhaB_C_^Ala^ when using Ala-tRNA^Ala^(UGC) generated by AlaRS (Figure S3). The BhaB_C_^Ala^ reaction efficiency increased with higher concentrations of Ala-tRNA^Ala^ prepared by flexizyme and reached complete modification, illustrating that the BhaB_C_^Ala^ enzymatic assay was compatible with flexizyme-prepared aa-tRNA (Figure S4). To compare various charged tRNA substrates, all subsequent assays were conducted at a range of acylated-tRNA concentrations (Supporting Information, Materials and Methods).

We next examined whether the cognate tRNA^Ala^(UGC) charged with different acids would be accepted by BhaB_C_^Ala^ (Figures 2, S5-S16). The selected acids, including proteinogenic and non-canonical amino acids, a hydroxy acid, and a mercaptocarboxylic acid, were first chemically activated and used as flexizyme substrates as previously reported.^36-42^ BhaB_C_^Ala^ catalyzed amide formation with all evaluated aminoacyl-tRNA^Ala^ analogs (Figures S5-S10). Except for Gly-tRNA^Ala^, which was accepted with efficiency akin to Ala-tRNA^Ala^, most proteinogenic analogs were accepted with significantly lower activity. We next evaluated non-canonical amino acids attached to tRNA^Ala^ (Figures S11-S14). While BhaB_C_^Ala^ catalyzed efficient addition of β-Ala,^36^ much lower efficiency was observed with D-Ala.^37^ We did not observe addition of 2-aminoisobutyric acid (Aib)^38^ or N-Me-Ala.^39^ BhaB_C_^Ala^ also catalyzed C-O and C-S bond formation when lactate and thioglycolic acid, respectively, were attached to tRNA^Ala^(UGC) (Figures S15-S16),^40,41^ although product formation was reduced compared to that observed with L-Ala and Gly. The resulting thioester product successfully underwent native chemical ligation^43^ with cysteamine (Figure S17). Collectively, these results show that BhaB_C_^Ala^ appears to have evolved to recognize L-Ala, but that other small amino acids and hydroxy/mercaptocarboxylic acids attached to tRNA^Ala^(UGC) are also accepted, showing considerable tolerance with respect to the nucleophile. The observation that various acyl groups attached to tRNA^Ala^(UGC) were substrates for BhaB_C_^Ala^ suggests the enzyme also recognizes features of the tRNA. Since Gly-tRNA^Ala^ was an efficient substrate, we first tested whether Gly-tRNA^Gly^(GCC) generated by *E. coli* GlyRS was accepted by BhaB_C_^Ala^. This reaction resulted in only moderate conversion (Figure S18) while the reaction with Gly-tRNA^Ala^(UGC) at similar concentration of substrates gave quantitative modification (Figure S6), indicating that tRNA identity contributes to substrate recognition. The results described thus far suggest that BhaB_C_^Ala^ evolved to recognize Ala-tRNA^Ala^ *in vivo*^18^ based on the combination of the amino acid and the tRNA component. To further examine this hypothesis, we used flexizyme to charge Ala onto various non-cognate tRNAs having different sequences (Figures 3, S19-S20, Table S1). The tRNAs for Trp, Glu and Cys were chosen because they are substrates of previously reported PEARLs or PEARL-like enzymes.^17,18,44,45^ All Ala-tRNA chimera lead to lower BhaB_C_^Ala^ reaction efficiency compared to Ala-tRNA^Ala^ (Figures S21-S24), with *P. syringae* Ala-tRNA^Cys^(GCA) and *E. coli* Ala-tRNA^Trp^(CCA) being particularly poor substrates (Figures 3, S22-S23).

**Figure 2.**
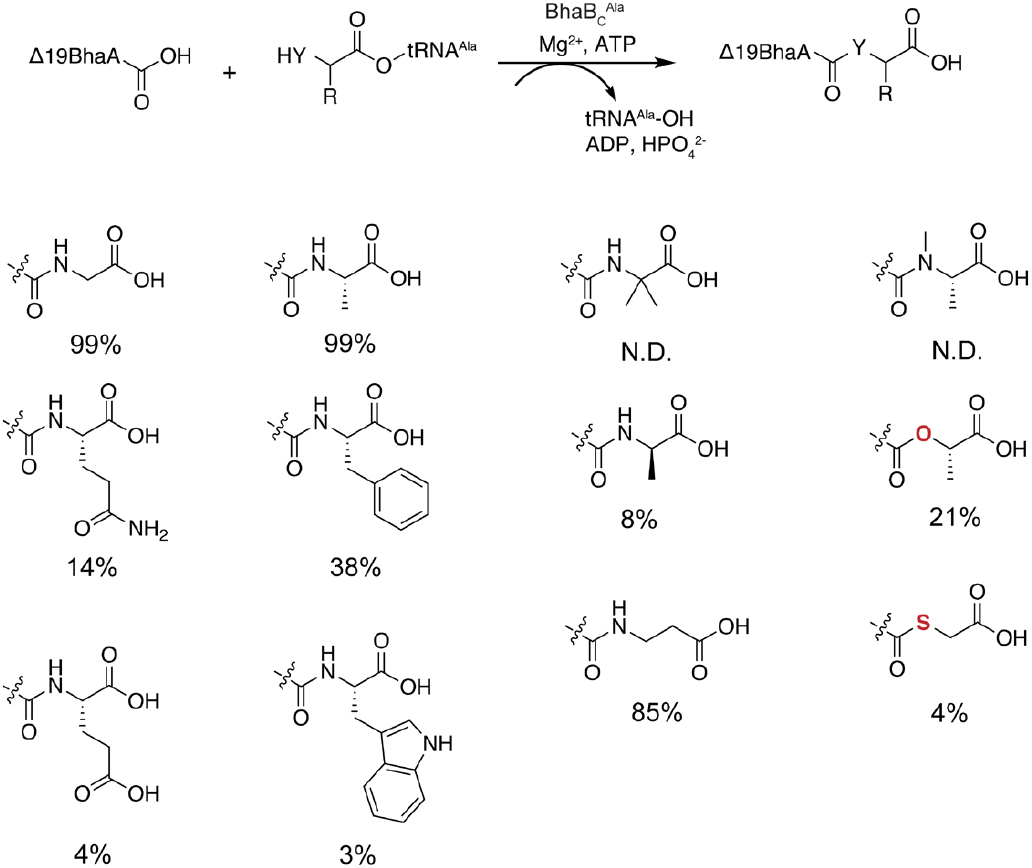
Amino acid specificity of BhaB_C_^Ala^. Various acids were charged onto *E. coli* tRNA^Ala^ using flexizyme. These aa-tRNA^Ala^ chimera were then tested for BhaB_C_^Ala^-catalyzed ligation with Δ19BhaA (sequence in Figure S3). Reaction efficiencies were evaluated at multiple aa-tRNA concentrations using LC-ESI-MS; values shown represent the highest efficiencies observed. All EIC traces are provided in Figures S4 and S6-S17. N.D. = not detected.

**Figure 3.**
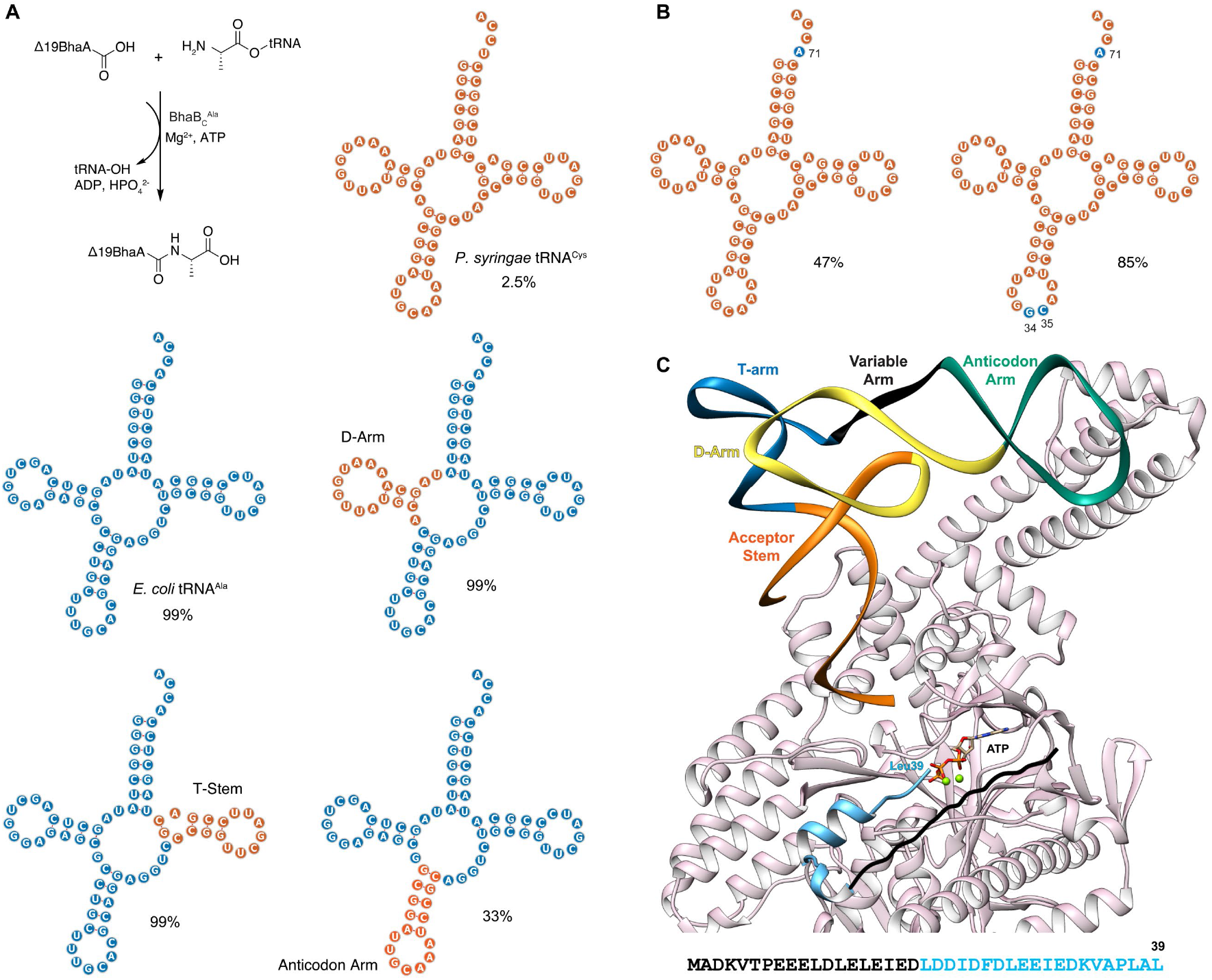
The tRNA specificity of BhaB_C_^Ala^. A) Various chimeric tRNAs with interchanged the D-arm, T-stem, or anticodon Arm of *E. coli* tRNA^Ala^ (UGC) and *P. syringae* tRNA^Cys^ (GCA) were aminoacylated with Ala using flexizyme. These Ala-tRNAs were used in the BhaB_C_^Ala^-catalyzed ligation with Δ19BhaA (sequence in Figure S3). B) Mutations in *P. syringae* tRNA^Cys^ (GCA) improved BhaB_C_^Ala^ activity. In panel A and B, regions derived from *E. coli* tRNA^Ala^ are in blue and regions from *P. syringae* tRNA^Cys^ in orange. The cloverleaf structures of each tRNA were predicted using tRNAScanSE^48^ and Forna.^49^ Reactions were performed using a range of Ala-tRNA concentrations and analyzed using LC-ESI-MS; results are summarized in Table S1; values shown represent the highest estimated efficiencies observed. EIC traces are provided in Figure S4 and Figures S22-S30. C) AlphaFold3 model of the complex of BhaB_C_^Ala^, BhaA, *E. coli* tRNA^Ala^, Mg^2+^, and ATP. The tRNA is colored by structural region: acceptor stem (orange), D-arm (yellow), anticodon arm (green), variable arm (black), and T-stem (blue). The sequence of ^Δ^19BhaA is shown in light blue, with the first 19 residues present in full-length BhaA in black. Predicted local distance difference test (pLDDT) scores and predicted aligned error (pAE) plots for the model are provided in Figure S31 and Supplementary Data 1. The protein image was made using Chimera.^50^

To determine which regions of the tRNA were critical for BhaB_C_^Ala^ recognition, we used flexizyme to charge Ala onto chimeric tRNAs having individual regions of *E. coli* tRNA^Ala^(UGC) replaced with the corresponding sequences from the poor substrate *P. syringae* tRNA^Cys^(GCA) (Figures 3A, S20-S21, S25-S27, Table S1). We observed quantitative product formation with chimera containing the D-arm and T-stem from tRNA^Cys^ (Figures 3A, S25-S26), albeit a reduction in efficiency was noted with the D-arm chimera at lower aa-tRNA concentrations. In contrast, grafting the anticodon arm from tRNA^Cys^(GCA) into tRNA^Ala^(UGC) significantly reduced activity, suggesting that the anticodon arm plays an important role in recognition (Figures 3A, S27). This conclusion was supported by introducing the anticodon arm from the poor substrate *E. coli* Ala-tRNA^Trp^(CCA) (Figure S23) into tRNA^Ala^(UGC), which also markedly decreased BhaB_C_^Ala^ activity (Figure S28). These results provide an explanation for the poor acceptance of Ala-tRNA^Trp^(CCA) and Ala-tRNA^Cys^(GCA), but do not explain why *T. bispora* Ala-tRNA^Glu^(CUC) was observed to be a more competent substrate (Figure S24).

We next evaluated the acceptor stem for BhaB_C_^Ala^ activity, a region of importance for related aa-tRNA utilizing enzymes.^46^ A variant of the poor substrate *P. syringae* Ala-tRNA^Cys^ carrying a U71A substitution to match the discriminator base in the acceptor stem of *E. coli* tRNA^Ala^, showed significantly improved activity compared to the wild type Ala-tRNA^Cys^(GCA) (Figures 3B, S29, Table S1). Use of this substrate by the enzyme was further improved by mutating the second and third position of the anticodon (GCA) to match that of *E. coli* tRNA^Ala^ (GGC) (Figures 3B, S30, Table S1), underscoring the importance of the anticodon. These findings also provide an explanation for Ala-tRNA^Glu^(CUC) being a moderately competent substrate. Unlike the poor substrates tRNA^Cys^ and tRNA^Trp^, this tRNA^Glu^ naturally has the same discriminator base and third base in the anticodon as tRNA^Ala^ (Figure S20). The collective results also agree well with an AlphaFold3^47^ model of BhaB_C_^Ala^, BhaA, and *E. coli* tRNA^Ala^(UGC), in which the enzyme primarily interacts with the acceptor stem and the anticodon arm of the tRNA (Figures 3C, S31).^23^ The AlphaFold3 model also shows that the first 19 residues of the substrate peptide do not interact with the protein, consistent with the findings above that these residues are not essential for BhaB_C_^Ala^ activity (Figures S3, S31-S32).

In summary, we used flexizyme to generate various non-cognate aa-tRNA pairs and examined the substrate specificity of BhaB_C_^Ala^, an amide-forming enzyme that naturally uses Ala-tRNA^Ala^. Our data show that the observed specificity of BhaB_C_^Ala^ toward its cognate aa-tRNA *in vivo*^18^ is determined by both the amino acid and the tRNA component. Since all our tRNA molecules were obtained by *in vitro* transcription, we currently cannot rule out that post-transcriptional modifications may contribute further to this specificity. Despite this specificity, *in vitro* BhaB_C_^Ala^ can extend peptide chains with a range of amino acids and can catalyze bond formation beyond amides, such as esters and thioesters. Both experimental and structural models indicate that BhaB_C_^Ala^ recognizes the tRNA through both the acceptor stem and the anticodon region, with contributions from the discriminator base and the anticodon sequence. These findings lay the groundwork for future potential directed evolution of BhaB_C_^Ala^ variants capable of efficiently extending peptide chains with diverse amino acids beyond the native substrate.

## ASSOCIATED CONTENT

### Supporting Information

The Supporting Information is available free of charge at https://pubs.acs.org/doi/

Experimental procedures, Figures S1-S33 showing assay data and AlphaFold models, and Tables S1-S5 with nucleotide sequences, primers, and MS data.

## AUTHOR INFORMATION

### Funding

This work was supported in part by grants from the National Institutes of Health (GM058822 to W.A.vdD. and GM123998 to D.A.M.), by KAKENHI (JP23H04546, JP24K01634, JP24H01754, and JP24K21760 to Y.G.; JP22H02218 to A.V.; and JP20H05618 to H.S.) from the Japan Society for the Promotion of Science, and by the Adopting Sustainable Partnerships for Innovative Research Ecosystem (JPMJAP2418) from the Japan Science and Technology Agency to Y.G. W.A.vdD is an Investigator of the Howard Hughes Medical Institute.

### Notes

The authors declare no competing financial interests.

## ACKNOWLEDGEMENTS

We thank Dr. Lingyang Zhu for assistance with quantitative NMR experiments and Dr. Hyunji Lee for providing reagents. The imager for PAGE gels in this study is maintained by the Core Facilities at the Carl R. Woese Institute for Genomic Biology.

## Notes

### Competing Interest Statement

The authors have declared no competing interest.

