## Supporting Information for "Aminoacyl-tRNA specificity of a ligase catalyzing non-ribosomal peptide extension"

[1] Carl R. Woese Institute for Genomic Biology, University of Illinois at Urbana-Champaign, 1206 West Gregory Drive, Urbana, Illinois, 61801, USA.
[2] Department of Chemistry and Howard Hughes Medical Institute, 600 South Mathews Avenue, University of Illinois at Urbana−Champaign, Urbana, Illinois 61801, USA.
[3] Department of Chemistry, Graduate School of Science, The University of Tokyo, 7-3-1, Hongo, Bunkyo, Tokyo 113-0033, Japan
[4] Department of Pharmacy and Pharmaceutical Sciences, Faculty of Science, National University of Singapore, Singapore 117544
[5] Department of Chemistry, Graduate School of Science, Kyoto University, Kitashirakawa-Oiwakecho, Sakyo, Kyoto 606-8502, Japan
[6] Department of Biochemistry, Vanderbilt University School of Medicine – Basic Sciences, Nashville, TN 37232, USA
[7] Department of Chemistry, Vanderbilt University, Nashville, TN 37232, USA
*

**Figure S32.** An active site view of the Alphafold3 model of BhaA, $BhaB_C^{Ala}$, ATP, $Mg^{2+}$, and *E. coli* $tRNA^{Ala}$(UGC) depicting that the first 19 residues of BhaA do not interact with $BhaB_C^{Ala}$. 42

### MATERIALS AND METHODS

#### Purification of $BhaB_C^{Ala}$

A previously constructed pET28-$BhaB_C^{Ala}$ plasmid (sequence in Table S2) was used to transform *E. coli* BL21(DE3) and the cells were selected on Luria Broth [10 g/L tryptone, 5 g/L yeast extract, 10 g/L NaCl] agar plates containing 50 µg/mL kanamycin (Kan), incubated at 37 °C for 12-14 h. Single colonies were picked and grown overnight in Terrific Broth [12 (g/L) tryptone, 24 (g/L) yeast extract, 0.072 M $K_2HPO_4$ and 0.017 M $KH_2PO_4$ and 0.4% glycerol (*v/v*)] containing 50 µg/mL kanamycin. The overnight culture was then diluted at a 1:100 (*v/v*) ratio in Terrific Broth containing the same amount of antibiotics in 1 L Fernbach flasks and grown at 37 °C with 220 rpm shaking until $OD_{600}$ reached 1.5. Then, 2 mM $MgCl_2$ and 0.5 mM IPTG were added, and the culture was allowed to continue growth at 18 °C with shaking at 220 rpm for 18 h. The cells were then harvested by centrifugation at 7,000 × g for 10 min, with the supernatant being discarded, and the pellets were frozen at -80 °C prior to purification.

The cells were then resuspended in lysis buffer (50 mM 4-(2-hydroxyethyl)piperazine-1-ethanesulfonic acid [HEPES], 500 mM NaCl, 5% glycerol, pH 7.5) containing 4 µg/mL DNAse I, 4 mg/mL lysozyme, 2 µM E64, 2 µM benzamidine, 2 µM leupeptin, using 5 mL of buffer per grams of wet cells. The cells were then lysed using a high-pressure homogenizer (Avenstin, Inc) at 10 kpsi with three homogenizer cycles. The homogenate was subjected to centrifugation at 36,000 × g for 1 h, and every 70 mL of the supernatant was applied to 1 mL of Ni-NTA resin (HisPur, Thermo). The mixture was rocked for 30 min at 4 °C, after which the Ni-NTA resin (HisPur, Thermo) was loaded onto a column and washed with 15 column volumes (CV) of lysis buffer [containing 1 mM tris(2-carboxyethyl)phosphine (TCEP), but without protease inhibitors and DNAse I] followed by 15 CV of wash buffer (50 mM HEPES, 1 M NaCl, 5% glycerol, 30 mM imidazole pH 7.5, 1 mM TCEP). The protein was then eluted using 6 CV of elution buffer (50 mM HEPES, 300 mM NaCl, 250 mM imidazole, pH 7.5, 1 mM TCEP). The protein was then buffer-exchanged to storage buffer 1 (50 mM HEPES, 100 mM NaCl, 2.5% glycerol, 1 mM TCEP pH 7.0) using a Slide-A-Lyzer G3 dialysis cassette (20 kDa molecular weight cutoff, MWCO) with at least a 100x volume exchange.

After dialysis, the protein mixture was clarified by centrifugation at 16,000 × g for 10 min. The supernatant was then injected into a 5 mL HiTrap SP HP (Cytiva) cation exchange column connected to an ÄKTA Start chromatography system. Solvent A consisted of 50 mM HEPES, 100 mM NaCl, 2.5% glycerol, and 1 mM TCEP, pH 7.0, while solvent B consisted of 50 mM HEPES, 1 M NaCl, 2.5% glycerol, and 1 mM TCEP, pH 7.0. The system was run at a flow rate of 5 mL/min, and the column was pre-equilibrated with solvent A before injection. After sample injection, the column was washed with 6 CV of solvent A. Following washing, a gradient elution was performed from 100% solvent A and 0% solvent B to 0% solvent A and 100% solvent B over 20 CV, with 4 mL fractions collected. Each fraction was then subjected to Sodium Dodecyl Sulfate Polyacrylamide gel electrophoresis (SDS-PAGE) using a 4–20% precast polyacrylamide gel (Biorad) and ran at 250 V for 30 min. Fractions corresponding to peaks containing $BhaB_C^{Ala}$ were then subjected to a $BhaB_C^{Ala}$ assay with and without AlaRS (see $BhaB_C^{Ala}$ assays with AlaRS) to identify those with minimal *E. coli* AlaRS carryover. Fractions with minimal carryover were pooled and concentrated to less than 2 mL using a 15 mL Amicon 30 kDa MWCO Ultra centrifugal filter (EMD Millipore).

Then 1 mL of the concentrated $BhaB_C^{Ala}$ was injected into a HiLoad 16/600 Superdex 200 pg size exclusion column pre-equilibrated with storage buffer 2 (50 mM HEPES, 300 mM NaCl, 2.5% glycerol, 1 mM TCEP, pH 7.0) on an ÄKTA Go system. The column was then run with an isocratic flow of 1 column volume (120 mL) using storage buffer 2 at a flow rate of 1.6 mL/min. Fractions were collected in 2 mL aliquots. Fractions corresponding to peaks containing $BhaB_C^{Ala}$ were subjected to a $BhaB_C^{Ala}$ assay $BhaB_C^{Ala}$ assay with and without AlaRS (see $BhaB_C^{Ala}$ assays with AlaRS) to identify those with minimal *E. coli* AlaRS carryover. Fractions with no AlaRS carryover were pooled (Figure S3), concentrated using a 4 mL Amicon 30 kDa MWCO Ultra centrifugal filter (EMD Millipore), flash frozen, and stored at -80 °C.

**In vitro transcription and purification of RNAs (flexizyme and tRNAs)**

The double-stranded DNA (dsDNA) templates for *in vitro* transcription were generated from overlapping primers[1] (Integrated DNA technologies) with sequences provided in Table S3. Specifically, 5' overhangs were assembled using the following extension reaction: 1 × NEB buffer 2, 4 µM of forward and reverse primers, 100 µM of each dNTPs, and 1 U of DNA polymerase I large (Klenow) fragment per 1 µg of DNA. These 50 µL reactions were incubated at 25°C for 15 min and then quenched with 10 mM EDTA at 75°C for 25 min. The DNA templates were cleaned by adding 5 µL (1/10 reaction volume) of 3 M NaOAc (pH 5.2) and 125 µL of cold ethanol, followed by incubation at 4 °C. The DNA pellet was collected by centrifugation at 16,000 × g for 15 min, and the supernatant was discarded. The pellet was washed twice with 500 µL of 70% ethanol and then air-dried for 15 min. Once dried, the pellet was dissolved in 12.5 µL of $H_2O$ and prepared for *in vitro* transcription.

The *in vitro* transcription reaction was performed using RNase-free reagents with the following components: 100 mM tris (pH 7.5), 7.5 mM each rNTP, 36 mM $MgCl_2$, 50 mM DTT, 0.1 mg/mL bovine serum albumin, 0.8 U/µL RNase inhibitor (RiboLock), 0.5 mU/µL *E. coli* inorganic phosphatase (Thermo Scientific), and 0.9 mg/mL T7 RNA polymerase. The DNA pellet (12.5 µL per 125 µL in vitro transcription reaction) was added to the transcription mix. The reaction was incubated at 37 °C overnight. To remove the DNA, 6.25 U of DNase I (Thermo Scientific) per 125 µL reaction and 1× DNase I reaction buffer with $MgCl_2$ (Thermo Scientific) were added.

The tRNAs were then purified using an RNA Monarch Kit (500 µg) (NEB) per instructions of the manufacturer. One spin column was used for each 125 µL of *in vitro* transcription reaction. The tRNAs were eluted from the spin column using 75 µL of RNAse-free $H_2O$. Then, 2 mM NaOAc pH 5.2 was added to the eluted tRNAs. The concentration of the tRNAs was assayed using the Qubit RNA Broad Range Assay Kit (ThermoFisher Scientific) on a Qubit 2.0 fluorometer (ThermoFisher Scientific).

Flexizymes (from 1 mL *in vitro* transcription reactions) were purified by adding 1/10 volume of 5 M $NH_4OAc$, 1/20 volume of 0.5 M EDTA, and an equal volume of isopropanol. After centrifugation at 16,000 × g for 10 min, the supernatant was removed, and the RNA pellet was washed with 800 µL of 70% ethanol. The RNA was then air-dried (covered with a Kimwipe) before being dissolved in 100 µL of $H_2O$ and added to an Amicon 10 kDa MWCO Ultra centrifugal filter (EMD Millipore). Following a 1,000× volume exchange with RNase-free $H_2O$, the RNA was purified from proteins using acidic phenol extraction. Specifically, an equal volume of acid-phenol: chloroform: isoamyl alcohol (25:24:1, pH 4.5) (Invitrogen) was added, and the mixture was shaken vigorously and centrifuged at 16,000 × g for 5 min. The aqueous layer was collected, and the organic layer was extracted with an equal volume of 0.3 M NaOAc (pH 5.2). The aqueous layers were combined and extracted again using the same protocol. Chloroform: isoamyl alcohol (24:1) (Sigma) was added to the collected aqueous layers, and the mixture was shaken vigorously and subjected to centrifugation at 16,000 × g for 5 min. The aqueous layer was collected and precipitated by adding 1/10 volume of 3 M NaOAc (pH 5.2) and 2.2 volumes of ethanol. The RNA pellet was collected by centrifugation at 16,000 × g for 15 min and washed twice with 75% ethanol. After removing the supernatant, the pellet was air-dried for 15 min (covered with a Kimwipe) before being redissolved in 2 mM NaOAc (pH 5.2). The concentration of the tRNAs was assayed using the Qubit RNA Broad Range Assay Kit (ThermoFisher Scientific) on a Qubit 2.0 fluorometer (ThermoFisher Scientific).

The quality of all RNAs was assessed using denaturing Urea-PAGE. Specifically, the tRNAs after eluting from the column were diluted 50 times using $H_2O$. 2 µL of diluted RNAs [concentration 70-270 (ng/µL)] was then mixed with 18 µL of 2x RNA loading dye (NEB) and heated at 95 °C for 3 min. Then, 3 µL was loaded onto 10% Mini-PROTEAN TBE-Urea Gel (Biorad) preheated at 250V for 30 min. The gel was run at 180 V for 36 min and stained with SYBR Gold (Thermo) before being visualized using the UV channel on ImageQuant800 (Cytiva) (Figure S33).

**Flexizyme assays to prepare aa-tRNAs**

The reactions were prepared by mixing 50 mM HEPES-KOH pH 7.5 (or 150 mM HEPES-NaOH pH 7.5), 41-61 µM eFx or 24 µM dFx,[2,3] 27-31 µM tRNA, 100 mM $MgCl_2$ for dFx (600 mM in case of Aib-DBE and Thiogly-DBE) or 400-600 mM $MgCl_2$ for eFx, and 5 mM activated amino acids. The amino acids (see Table S5) were activated by chemical synthesis with 4-chlorobenzyl thioester (CBT, used with eFx), 3,5-dinitrobenzyl ester (DBE, used with dFx), or cyanomethyl ester (CME, used with eFx) according to published protocols, purified, and stored in DMSO.[2,4-8] For a 110 µL reaction, the detailed protocol for charging of tRNA was as follows: First, HEPES, flexizyme, tRNA, and $H_2O$ were mixed well together up to 66 µL in a 1.7 mL Eppendorf tube. The mixture was heated to 95 °C in an aluminum heating block for 2 min and then cooled down at room temperature on air for 7 min. Next, 22 µL of 500 mM – 3 M $MgCl_2$ stock was added, and the mixture was incubated at room temperature for 5 min. The reaction was placed on ice for 3 min, after which 22 µL of 25 mM activated amino acid (stored in DMSO) was added. The reactions were incubated on ice with time specified in Table S5.

When significant precipitation was observed due to amino acid insolubility, DMSO was added to improve solubility. For a 110 µL Lac-CBT reaction, 22 µL of DMSO along with 5 µL of 500 mM HEPES (pH 7.5) were added to the reaction mixture after adding Lac-CBT. In the case of thioglycolate-CBT (Thiogly-CBT), the reaction was run in 150 mM Bicine pH 9 containing 5 mM DTT and 40% (*v/v*) DMSO instead of 20% (*v/v*) DMSO.

The RNA clean-up protocol was performed at room temperature as follows, with examples of volumes of solutions applied to each step described in the table below. First, the flexizyme reactions were quenched by adding 0.3 M NaOAc (pH 5.2). The resulting mixture was divided into two tubes, one for downstream $NaIO_4$-RNA extension assays and one for downstream $BhaB_C^{Ala}$ assays, with volumes specified in the accompanying table. EtOH was added to the quenched reactions, which were then centrifuged at 16,000 × g for 15 min. The supernatant was discarded, and the pellet was washed by adding 70% EtOH 0.1 M NaOAc (pH 5.2) (made by mixing aqueous 3M NaOAc pH 5.2 with ethanol and water) and vortexed. The mixture was centrifuged again at 16,000 × g for 5 min, and the wash was repeated. The pellet was then washed with 70% EtOH 1 mM NaOAc (pH 5.2) (diluted from 70% EtOH 0.1 M NaOAc (pH 5.2) with 70% EtOH) and centrifuged at 16,000 × g for 3 min. The supernatant was discarded, and the pellet was air-dried for 15 min (covered with a Kimwipe). The pellets were stored at -80 °C for a maximum of 5 days until use. A table describing the specific volumes is shown below.

| | | 0.3 M NaOAc pH 5.2 | EtOH | 70% EtOH 0.1 M NaOAc (pH 5.2) | 70% EtOH 1 mM NaOAc (pH 5.2) |
|---|---|---|---|---|---|
| Flexizyme reaction (total) | 110 µL | Add 440 µL. Total volume = 550 µL | | | |
| For $BhaB_C^{Ala}$ assays | | 500 µL | 1000 µL | 600 µL | 400 µL |
| For $NaIO_4$ - RNA extension assays | | 50 µL | 100 µL | 60 µL | 40 µL |

**$NaIO_4$-RNA extension assays to measure tRNA aminoacylation.**[9-11]

The schematic of this assay is described in Figure S2. The RNA pellets cleaned up from 10 µL of Fx reactions were dissolved in 15 µL of 1 mM NaOAc pH 5.2. The solution was then split into three Eppendorf tubes, each containing 5 µL of RNA, to run three reactions corresponding to Schemes 1, 2, and 3 in Figure S2.

In the tube for Scheme 1: 42.5 µL of RNA buffer (50 mM NaOAc, pH 5.2, 150 mM NaCl, 10 mM $MgCl_2$, and 0.1 mM EDTA) was added to the RNA solution. Then, 2.5 µL of 100 mM $NaIO_4$ was added, and the reaction was incubated for 1 h at 10 °C. The reaction was quenched by adding 10 µL of 100 mM DTT and left on ice.

In the tube for Scheme 2: 45 µL of the RNA buffer was added to the RNA solution.

In the tube for Scheme 3: 40 µL of 1× deacylation buffer (50 mM Bicine-NaOH pH 9.7, and 1 mM EDTA) was added to the RNA solution, and the mixture was incubated at 42 °C in an aluminum heating block for 36 min. The reaction was quenched by adding 6 µL of 3 M NaOAc. Then, 2.7 µL of 100 mM $NaIO_4$ was added, and the reaction was incubated for 1 h at 10 °C. The reaction was quenched by adding 10 µL of 100 mM DTT and left on ice for 20 min.

All three mixtures were then purified using the Monarch RNA cleanup kit (10 µg) following the protocol of the manufacturer. The processed RNAs were then eluted from the spin columns using 12 µL of RNAse-free $H_2O$. The RNAs were then quantified using $A_{260}$ using a spectrophotometer.

The purified RNAs were subjected to the RNA extension reaction as follows: 1× NEB Buffer 2, 500–750 nM RNA, 1 µM Cy3-labeled primer (see Table S4), 400 µM of each dNTP (Goldbio), and $H_2O$ to bring the total volume to 23.5-24.5 µL in a 0.2 mL PCR tube. The mixture was subjected to the following cycle on a thermocycler: 95°C for 2 min, 70°C for 2 min, 50°C for 2 min, and then held at 4 °C. Next, 0.5-1.5 µL of Klenow Fragment (3'→5' exo-) (M0212S) was added, and the reaction was incubated at 37 °C for 45 min. An equal volume of 2× loading dye (8 M urea, 0.06% Orange G) was added to the reaction mixture.

The resulting mixture was subjected to PAGE using a 15% Mini-Protean TBE-Urea gel, 10-well, which was pre-run at 250 V for 25-30 min in 1× TBE buffer. Each well was rinsed thoroughly with the running buffer, and immediately afterwards, a 10 µL aliquot of the reaction mixture was loaded into the wells. The gel was run at 200 V for 150-180 min. The gel was then imaged using the Cy3 fluorescence channel on the ImageQuant800 (Cytiva) fluorescence application. The bands in each lane, which represented assayed free and aminoacylated tRNAs, were estimated relative to each other using ImageJ based on peak height. This estimation and the concentration of free tRNAs applied to the flexizyme assay were used to estimate the concentration of aminoacyl tRNAs being applied to the $BhaB_C^{Ala}$ reaction.

**$BhaB_C^{Ala}$ assays with AlaRS.**

The concentration of a synthetic Δ19BhaA (1.3 mM; Genscript) solution in a buffered $D_2O$ solution (600 µL of 100 mM potassium phosphate buffer at pH 7.5 resuspended in $D_2O$) was quantified using quantitative $^1H$ NMR (qNMR) on a Bruker Avance NEO 600 MHz spectrometer equipped with a 5-mm BBO prodigy probe, following standard protocols.[12] The solution was measured inside a 5-mm Wilmad 535-pp NMR tube. A solution of 18 µM of Δ19BhaA was mixed well with 100 mM HEPES-NaOH (pH 7.5), 100 mM NaCl, 5 mM $MgCl_2$, 5 mM ATP (from 50 mM ATP stock with pH adjusted to 7.5 prior to usage), 1 mM TCEP (from 100 mM TCEP stock with pH adjusted to 7.0 prior to usage), 5 mM L-Ala, and 13 µM *E. coli* $tRNA^{Ala}$ (UGC). If AlaRS was used, 10 µM *E. coli* AlaRS, purified according to a published protocol,[13] was added to the reaction mixture, and mixed thoroughly. Then, 10 µM of $BhaB_C^{Ala}$ was added, and the reaction proceeded at room temperature overnight. Two volumes of cold acetonitrile (put at -20 °C prior to usage) were then added to the reaction mixture, and precipitates were removed by centrifugation at 16,000 × g for 10 min. The solvents from the collected supernatant were removed by rotary evaporation using a SpeedVac concentrator (Thermo). The resulting dried powder was then dissolved in an equal volume of $H_2O$ to the original reaction volume. Then, 1 µL was subjected to mass spectral analysis (see Mass spectral analysis of the $BhaB_C^{Ala}$ assays).

**Sequences.**

BhaA: MADKVTPEEELDLELEIEDLDDIDFDLEEIEDKVAPLAL

Δ19BhaA: LDDIDFDLEEIEDKVAPLAL

**$BhaB_C^{Ala}$ assays with Gly-$tRNA^{Gly}$ generated from GlyRS.**

In the case GlyRS was used to generate Gly-$tRNA^{Gly}$ (GCC), GlyRS was purified by Ni-affinity chromatography as reported in previous protocols.[13] 5 µM *E. coli* GlyRS was used under similar concentrations as mentioned above for AlaRS and the reaction was incubated for 4 h at 37 °C. The reaction mixture was purified by Monarch RNA spin column according to the manufacturer protocol and the aminoacylated tRNA was used in $BhaB_C^{Ala}$ reactions as indicated below. An aliquot of the aminoacylated $tRNA^{Gly}$ was utilized in $NaIO_4$-RNA extension reactions to estimate the degree of aminoacylation. The 25 µL reaction mixture with $BhaB_C^{Ala}$ contained: 100 mM HEPES (pH 7.5), 100 mM NaCl, 5 mM ATP (from 50 mM ATP stock with pH adjusted to 7.5 prior to usage), 1 mM TCEP (from 100 mM TCEP stock with pH adjusted to 7.0 prior to usage), 5 mM $MgCl_2$, 1.6 µM Δ19BhaA (concentration determined by qNMR as described above), 8.8 µM Gly-$tRNA^{Gly}$ (as determined by the estimated aminoacylation efficiency), and 4 µM $BhaB_C^{Ala}$. This mixture was incubated overnight at RT, quenched with an equal volume of cold acetonitrile (put at -20 °C prior to usage), subjected to centrifugation at 16,000 × g for 5 min, and an aliquot of 5 µL submitted for LC-MS analysis.

**$BhaB_C^{Ala}$ assays with aa-tRNA prepared by flexizyme.**

The reaction mixtures contained: 100 mM HEPES (pH 7.5), 100 mM NaCl, 5 mM ATP (from 50 mM ATP stock with pH adjusted to 7.5 prior to usage), 1 mM TCEP (from 100 mM TCEP stock with pH adjusted to 7.0 prior to usage), 5 mM $MgCl_2$, 1.8 µM Δ19BhaA (concentration determined by qNMR as described above), $BhaB_C^{Ala}$ (4 µM for non-cognate amino acid experiments and 6 µM for non-cognate tRNA experiments), and serial dilutions of aa-tRNA concentrations. The aminoacylation efficiency of the tRNA, estimated from $NaIO_4$-RNA extension assays (Figures S2, S5, S21), was used to estimate the amount of aa-tRNA applied in the $BhaB_C^{Ala}$ assays. The concentrations of aa-tRNA are shown in the extracted ion chromatograms (EIC) shown in Figures S4, S6-S16, and S22-S30. The detailed protocol for 10 µL reactions was as follows. First, a mixture of buffer-cofactor-peptide was assembled by mixing HEPES, NaCl, ATP, TCEP, $MgCl_2$, Δ19BhaA, and $H_2O$ to a volume of 7.7 µL. The aa-tRNA pellet (from ethanol precipitation of 50-100 µL flexizyme reactions as described above) was resuspended in 9.8 µL of 1 mM NaOAc (pH 5.2) and added to the mixture, bringing the volume to 17.5 µL. To perform serial dilutions of aa-tRNAs, four additional buffer-cofactor-peptide mixtures (7.7 µL each) were prepared as above, and 9.8 µL of 1 mM NaOAc (pH 5.2) was added instead of aa-tRNA. 8.75 µL of the aa-tRNA-containing mixture was then mixed in 8.75 µL with each of the four buffer-cofactor-peptide (no aa-tRNA) mixtures in sequence to achieve serial 2-fold dilutions. This procedure resulted in five reaction tubes, each with a total volume of 8.75 µL and identical amounts of all components except aa-tRNA. Then, 1.25 µL $BhaB_C^{Ala}$ (32 µM for non-cognate amino acid experiments and 48 µM for non-cognate tRNA experiments), purified via ion-exchange chromatography followed by size-exclusion chromatography as mentioned above, was added to each reaction mixture, and the reactions were incubated overnight at room temperature. In the case of thiogly-$tRNA^{Ala}$, an additional reaction containing 1 mM cysteamine (pH 7.5) was performed to assess native chemical ligation with the resulting thioester. A 6 µL aliquot of each reaction was then injected directly into the column or subjected to acetonitrile precipitation prior to mass spectral analysis. Specifically, 4 µL aliquot of the reaction was mixed with 8 µL of cold acetonitrile (put at -20 °C prior to usage), and precipitates were removed by centrifugation at 16,000 × g for 10 min. 6 µL of the clarified supernatants were then subjected to mass spectral analysis.

**Mass spectral analysis of the $BhaB_C^{Ala}$ assays**

Reactions to be analyzed were injected into a Kinetex C8 LC column (2.1 × 150 mm, 2.6 µm, 100 Å) on an Agilent 6545B Q-TOF interfaced with an Agilent 1290 Infinity II LC system. The mobile phase was: solvent A: $H_2O$ with 0.1% formic acid (FA), solvent B: $CH_3CN$ with 0.1% FA. The flow rate was 0.4 mL/min. The gradient was as follows: first 3 min 5% B, next 8 min 5% B - 80% B, last 2 min 80–95% B. Before the next samples were injected, the column was equilibrated with 3 min of 5% B. The samples were run to waste in the first 4 min before applying to the mass spectrometer. MS parameters were as follows: mass range 100 to 3000 *m/z*; 325 °C at 10 L/min; nebulizer, 35 psig; nozzle voltage, 0 V; sheath gas, 375 °C at 11 L/min; capillary, 3500 V; fragmentor, 120 V; skimmer, 65 V; MS scan rate (10 spectra/s). Positive ionization mode was utilized for all MS experiments. The EICs of species corresponding to the substrate and products were generated based on the *m/z* (z= 2) of each peptide. The reaction efficiency (x %) was estimated using the peak area of each EIC (A) as follows: $x = [A_{product} /(A_{product} + A_{substrate})]$ (%), with the assumption that the ionization efficiency of the substrate and the product is very similar, as in other similar studies.[14,15] This assumption is supported by the highly similar chemical nature of the substrates and products because their relative masses are very close and the reactions do not introduce additional positively charged moieties. However, due to the complex nature of ESI-MS causing potential subtle differences in ionization efficiency of different peptides,[16,17] the method should be considered semi-quantitative.

**Figure S1.** Mass spectrometry results of BhaA modified by $BhaB_C^{Ala}$ using different isoforms of *E. coli* $tRNA^{Ala}$. The reactions were analyzed by matrix-assisted laser desorption/ionization time-of-flight mass spectrometry (MALDI-TOF-MS). Reactions contained the indicated $tRNA^{Ala}$ (6 µM), BhaA (20 µM), $BhaB_C^{Ala}$ (3 µM), ATP (5 mM), alanine (8 mM), and *E. coli* AlaRS (3 µM) in a solution of HEPES (100 mM, pH 7.5), $Mg^{2+}$ (10 mM), and KCl (10 mM) in a 25-µL reaction volume. The asterisk (*) represents deamination of the indicated peptide species, a result specific to reflector positive mode in MALDI-TOF-MS.[18,19] Sequence of $His_6$-BhaA:

GSSHHHHHHSSGLVPRGSHMADKVTPEEELDLELEIEDLDDIDFDLEEIEDKVAPLAL

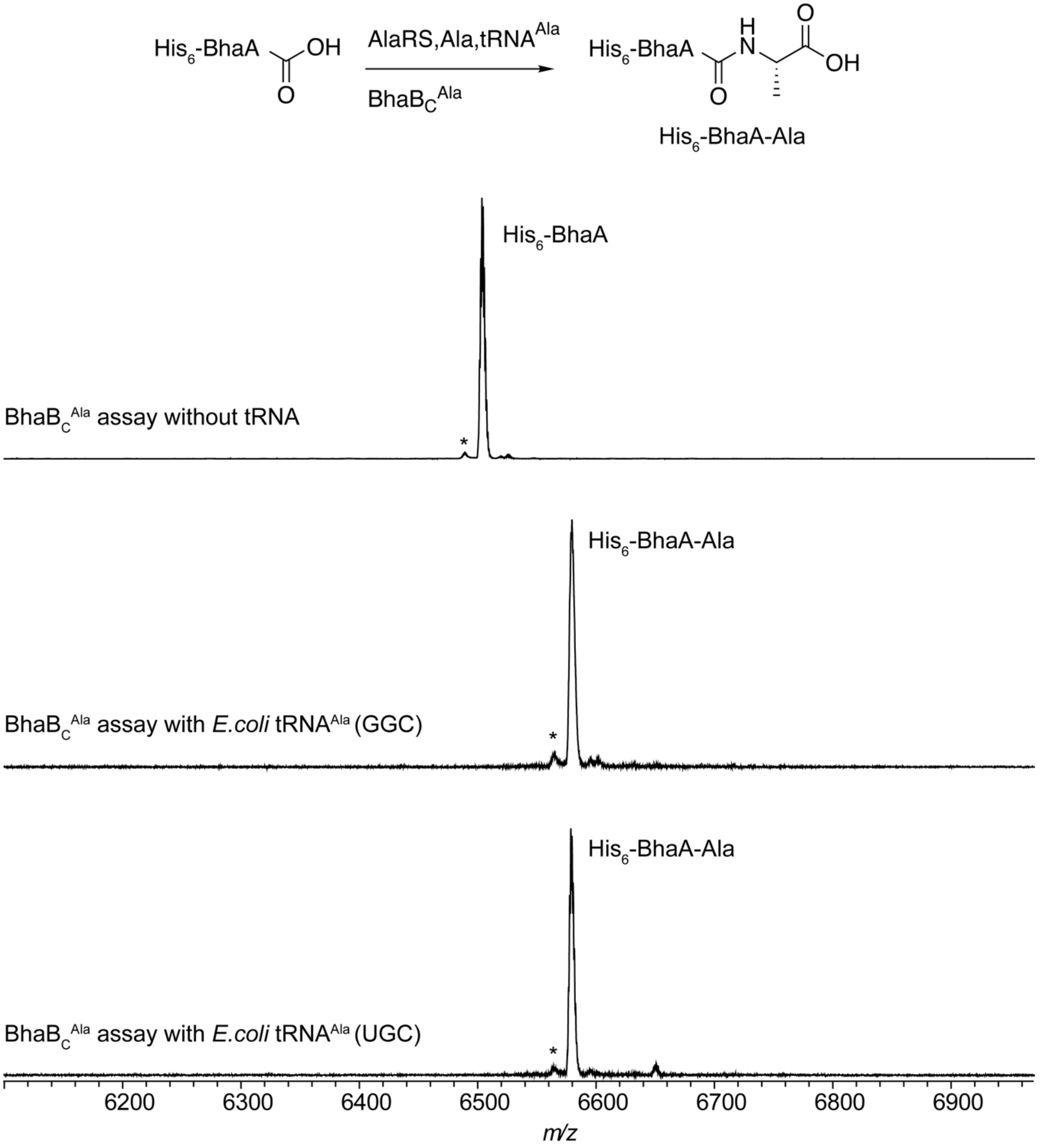

**Figure S2.** The $NaIO_4$-RNA extension assay. A) Schematic of the assay used to determine the efficiency of the flexizyme-catalyzed tRNA aminoacylation reactions.[9-11] tRNAs bearing free 3′-ends were selectively oxidized by $NaIO_4$, preventing templated extension by DNA Polymerase I Klenow fragment. [9-11] In contrast, aminoacylated tRNAs were protected from oxidation and permit extension, resulting in a distinguishable size difference upon PAGE analysis. Cy3 was used for visualization using a fluorescence imager. B) PAGE analysis with a fluorescence imager (Cy3 channel) of the method described using Ala-tRNA$^{Ala}$ (UGC) of *E. coli*. Detailed experimental conditions are in Materials and Methods. In this experiment (R1), the aminoacylation efficiency is estimated at 35%. The sequence of the primer is provided in Table S4. C) Calibration lanes generated by mixing species a and b at the indicated [a/(a+b)] percentages.

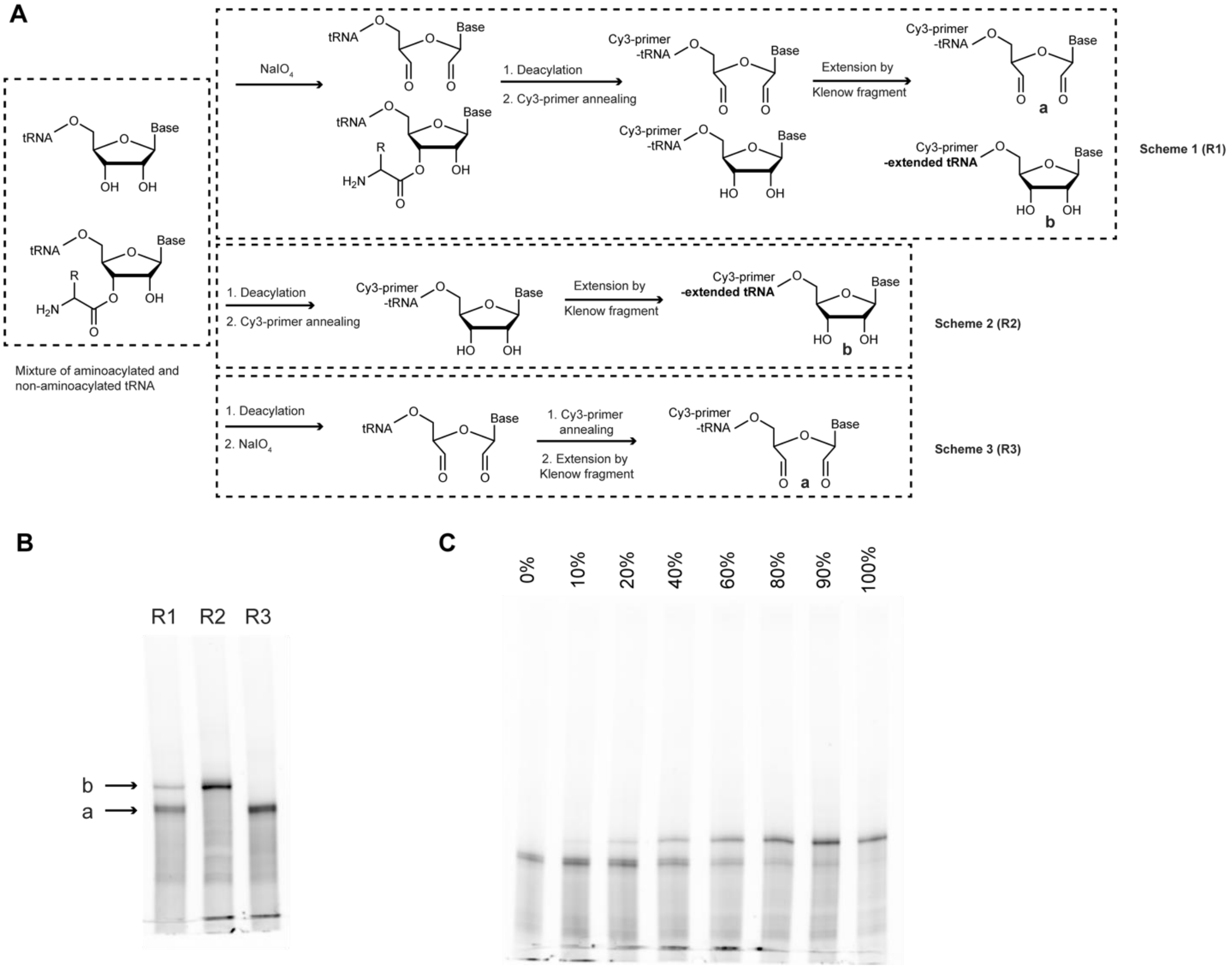

**Figure S3.** Examination of Δ19BhaA as a substrate of $BhaB_C^{Ala}$. Extracted ion chromatograms (EIC) of the liquid chromatography-mass spectrometry (LC-MS) trace of the $BhaB_C^{Ala}$ reaction using Δ19BhaA substrate peptide (sequence shown on top). The aminoacyl-tRNA complex of Ala and *E. coli* $tRNA^{Ala}$ (UGC) was generated by AlaRS. The black trace is the EIC of the substrate Δ19BhaA. The red trace is the EIC of the product Δ19BhaA-Ala.

Δ19BhaA: LDDIDFDLEEIEDKVAPLAL

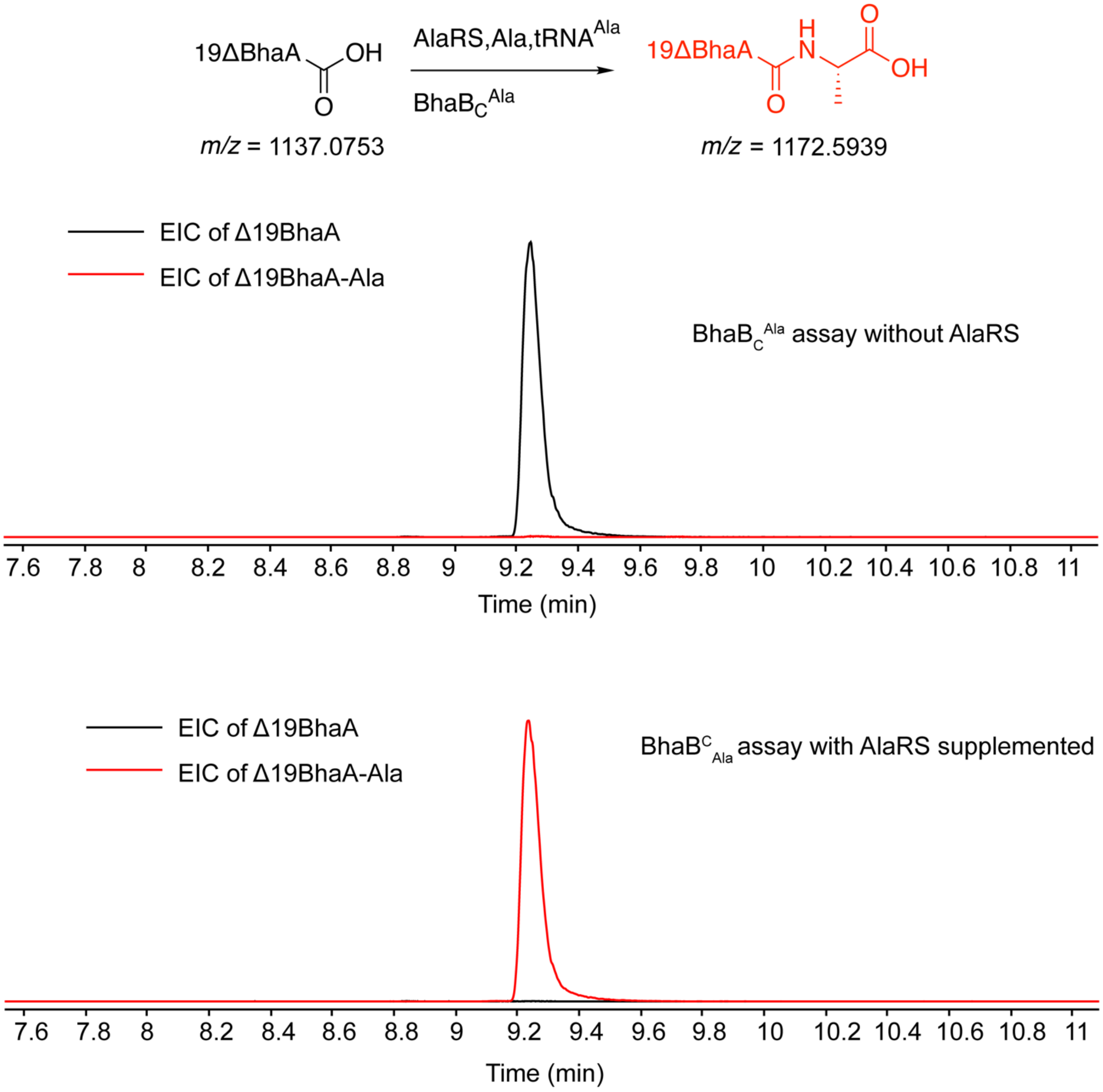

**Figure S4.** EICs of Δ19BhaA modified by BhaB$_C^{Ala}$ when reacting Ala *E. coli* tRNA$^{Ala}$ (UGC) charged with using flexizyme. Each panel represents BhaB$_C^{Ala}$ reaction when utilizing a different amount of flexizyme-generated Ala-tRNA$^{Ala}$. The concentration of the substrate peptide (Δ19BhaA) was 1.8 µM. The estimated concentration of Ala-tRNA$^{Ala}$ in each panel using the method in Fig. S1 was as follows: A) 37 µM, B) 19 µM, C) 9.3 µM, D) 4.7 µM, and E) 2.3 µM. The estimated reaction efficiency was A) 99%, B) 97%, C), D) 98%, and E) 72%. The black trace is the EIC of the substrate Δ19BhaA. The red trace is the EIC of the product Δ19BhaA-Ala. F) The mass spectrum of the substrate and the product. The calculated error can be found in Table S6.

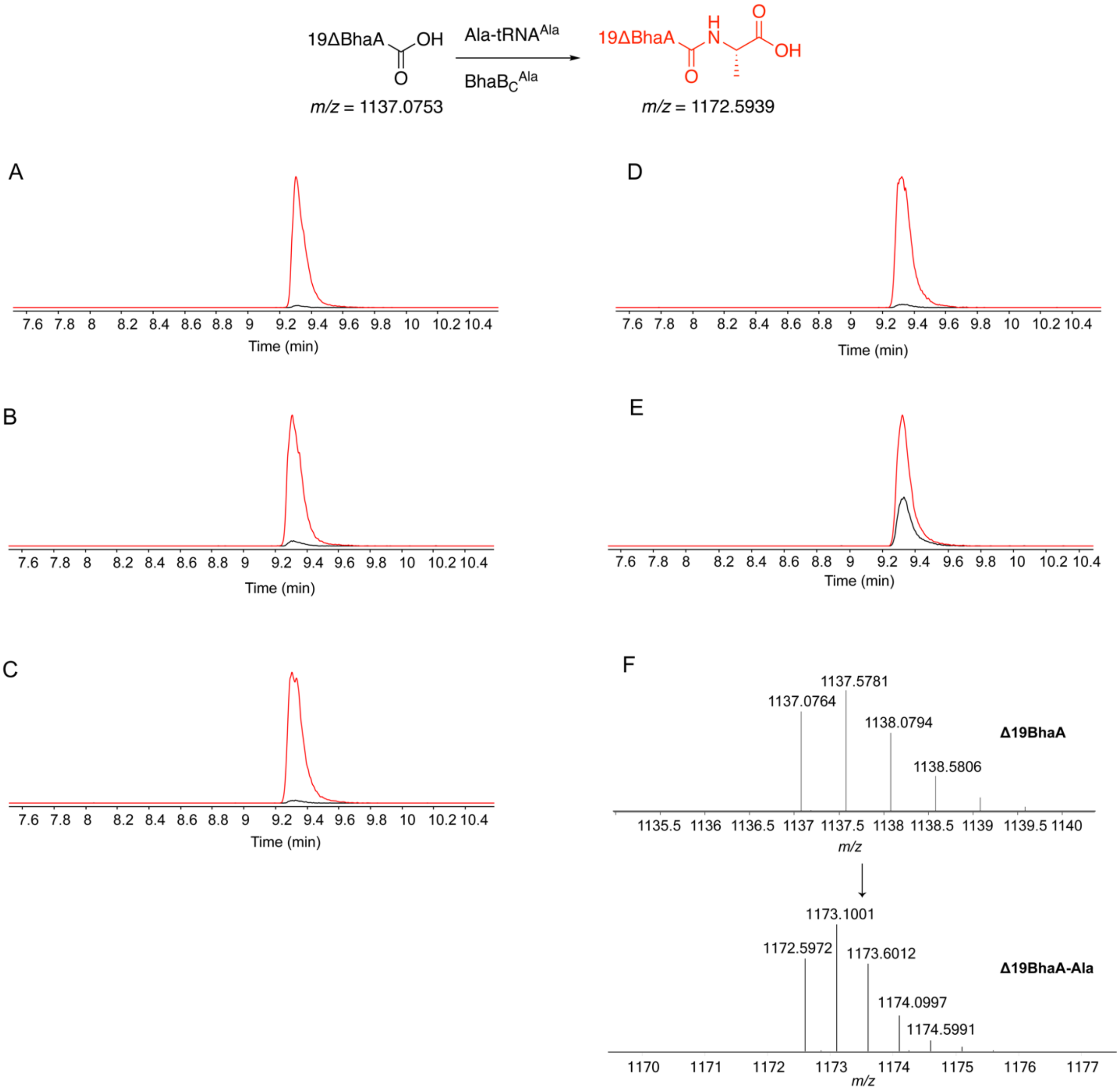

**Figure S5.** PAGE analysis of the $NaIO_4$-RNA extension assays (Figure S2) on flexizyme-catalyzed acylation of *E. coli* $tRNA^{Ala}$(UGC) with various acids. The upper band represents the product formed from aminoacylated-tRNA (species b, Figure S2) and the lower band represents the free tRNA (species a, Figure S2). Lane 1: N-methyl-L-Ala (estimated aminoacylation efficiency: 29%); Lane 2: D-Ala (30%), Lane 3: L-Glu (16%), Lane 4: L-Gln (11%), Lane 5: L-Trp (23%), Lane 6: 2-aminoisobutyric acid (Aib) (26%), Lane 7: β-Ala (17%), Lane 8: Gly (40%), Lane 9: Phe (28%). The result with L-Ala is shown in Figure S2B.

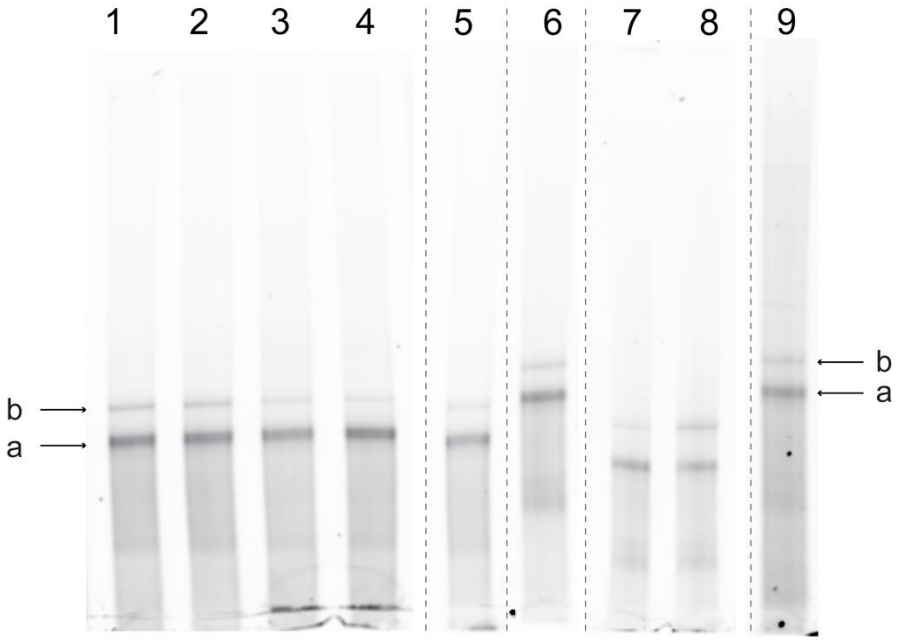

**Figure S6.** EICs of Δ19BhaA modified by $BhaB_C^{Ala}$ in the presence of *E. coli* $tRNA^{Ala}$ (UGC) charged with Gly. Each panel represents the $BhaB_C^{Ala}$ reaction when utilizing different amounts of Gly-$tRNA^{Ala}$. The concentration of the substrate peptide (Δ19BhaA) was 1.8 µM. The estimated concentration of Gly-$tRNA^{Ala}$ in each panel was as follows: A) 27 µM, B) 14 µM, C) 6.8 µM, D) 3.4 µM, E) 1.7 µM. The estimated reaction efficiency is as follows: A) 99%, B) 99%, C) 95%, D) 63%%, E) 32%. The black trace is the EIC of the substrate Δ19BhaA. The red trace is the EIC of the product Δ19BhaA-Gly. F) The ESI mass spectrum of the product. The calculated error comparing calculated and observed masses is shown in Table S6.

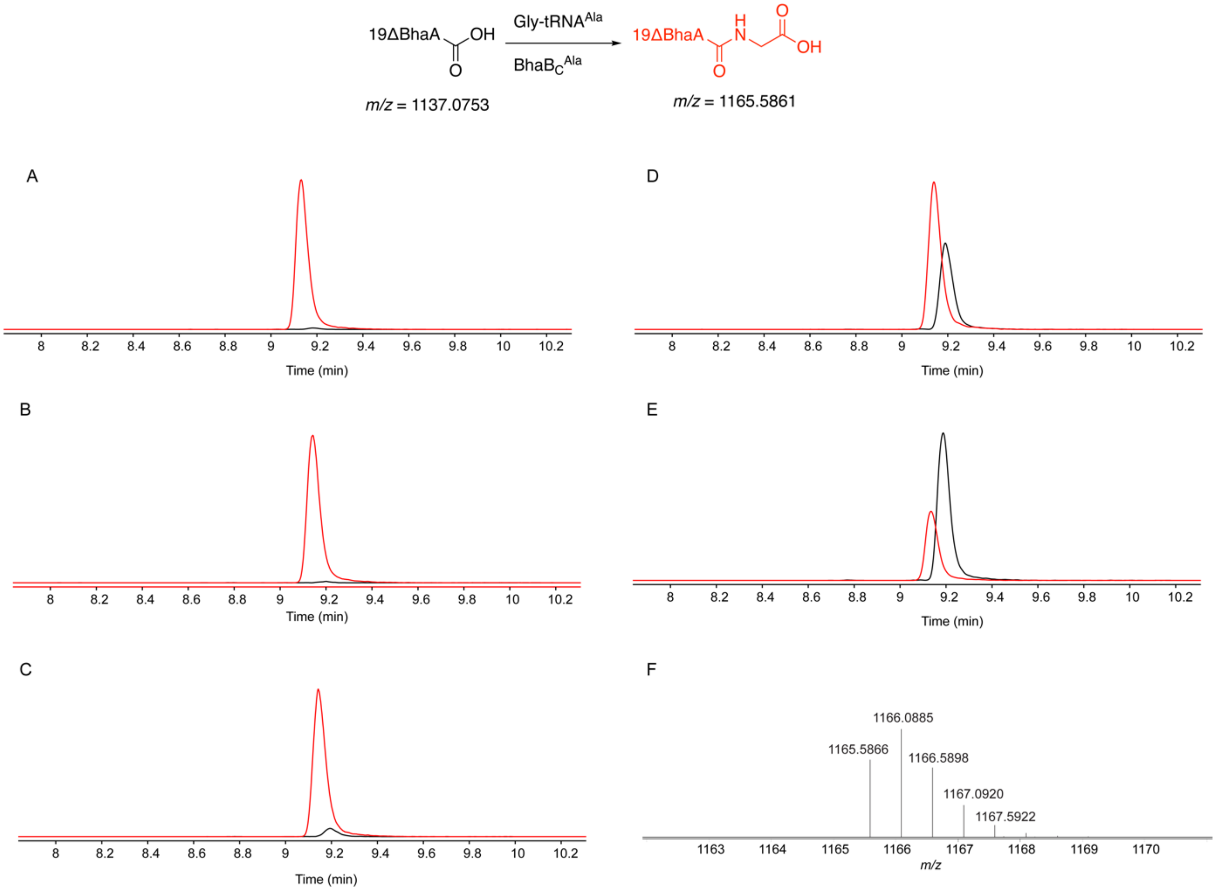

**Figure S7**. EICs of Δ19BhaA modified by $BhaB_C^{Ala}$ in the presence of *E. coli* $tRNA^{Ala}$ (UGC) charged with Gln. Each panel represents a $BhaB_C^{Ala}$ reaction utilizing different amounts of Gln-$tRNA^{Ala}$. The concentration of the substrate peptide (Δ19BhaA) was 1.8 µM. The estimated concentration of Gln-$tRNA^{Ala}$ in each panel was as follows: A) 15 µM, B) 7.7 µM, C) 3.8 µM, D) 1.9 µM, E) 0.96 µM. The estimated reaction efficiency is as follows: A) 14%, B) 9.1% , C) 6.4%, D) 5.0%, E) 2.9%. The black trace is EIC of the substrate Δ19BhaA. The red trace is EIC of the product Δ19BhaA-Gln. F) The mass spectrum of the product. The calculated error comparing calculated and observed masses is shown in Table S6.

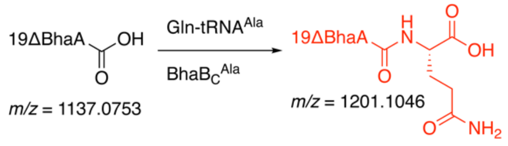

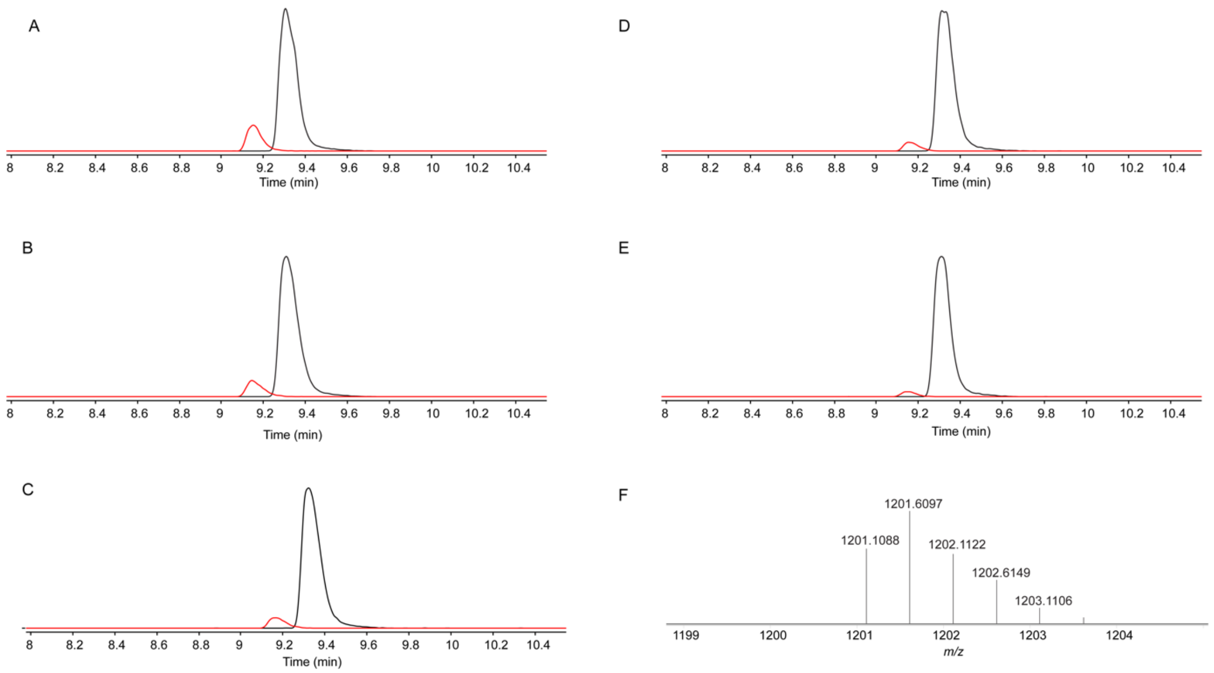

**Figure S8.** EICs of Δ19BhaA modified by $BhaB_C^{Ala}$ in the presence of *E. coli* $tRNA^{Ala}$ (UGC) charged with Glu. Each panel represents a $BhaB_C^{Ala}$ reaction when utilizing different amounts of Glu-$tRNA^{Ala}$. The concentration of the substrate peptide **(**Δ19BhaA**)** was 1.8 µM. The estimated concentration of Glu-$tRNA^{Ala}$ in each panel was as follows: A) 22 µM, B) 11 µM, C) 5.5 µM, D) 2.8 µM, E) 1.4 µM. The estimated reaction efficiency is as follows: A) 3.6%, B) 2.8% ,C) 2.4%, D) 1.9%, E) 1.3%. The black trace is EIC of the substrate Δ19BhaA. The red trace is EIC of the product Δ19BhaA-Glu. F) The mass spectrum of the product. The calculated error comparing calculated and observed masses is shown in Table S6.

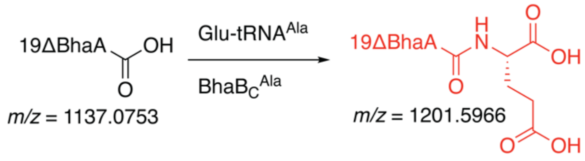

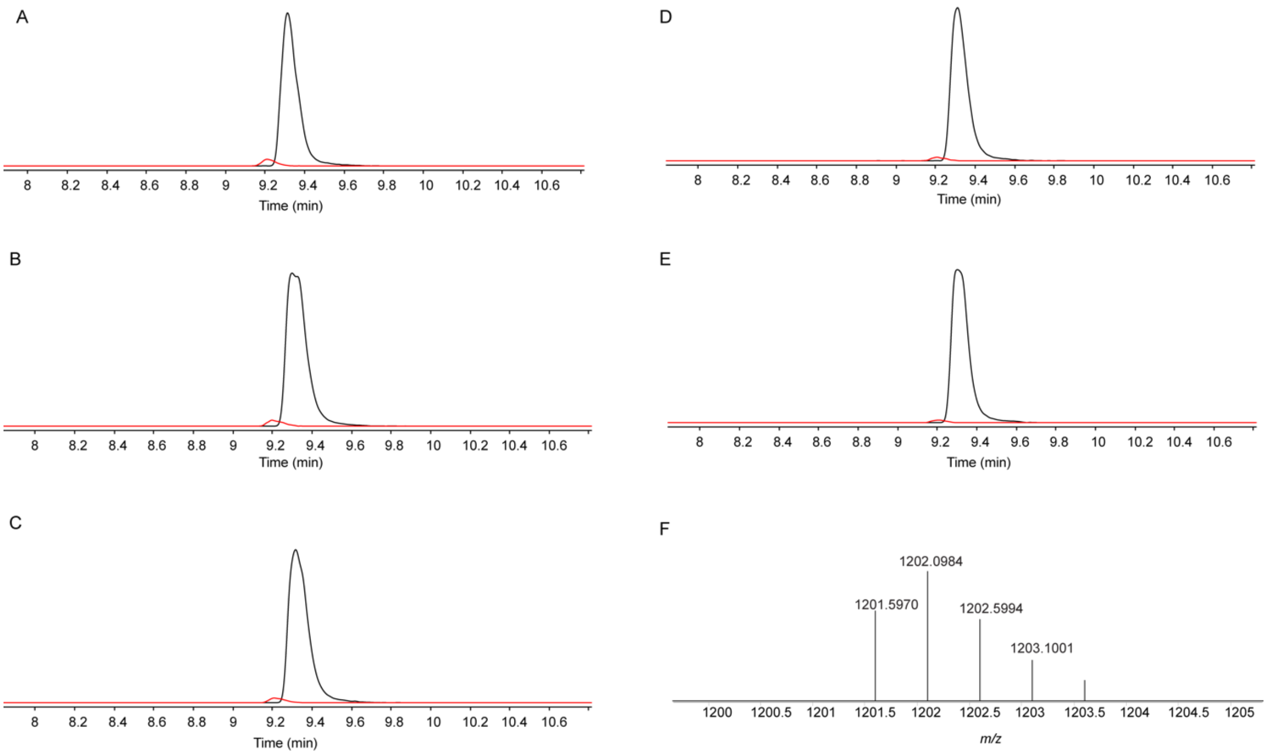

**Figure S9.** EICs of Δ19BhaA modified by $BhaB_C^{Ala}$ in the presence of *E. coli* $tRNA^{Ala}$ (UGC) charged with Phe. Each panel represents a $BhaB_C^{Ala}$ reaction when utilizing different amounts of Phe-$tRNA^{Ala}$. The concentration of the substrate peptide **(**Δ19BhaA) was 1.8 µM. The estimated concentration of Phe-$tRNA^{Ala}$ in each panel was as follows: A) 38 µM , B) 19 µM, C), 9.4 µM D) 4.7 µM , E) 2.4 µM. The estimated reaction efficiency is as follows: A) 33%, B) 38% , C) 33%, D) 24%, E) 16%. The black trace is EIC of the substrate Δ19BhaA. The red trace is EIC of the product Δ19BhaA-Phe. F) The mass spectrum of the product. The calculated error comparing calculated and observed masses is shown in Table S6.

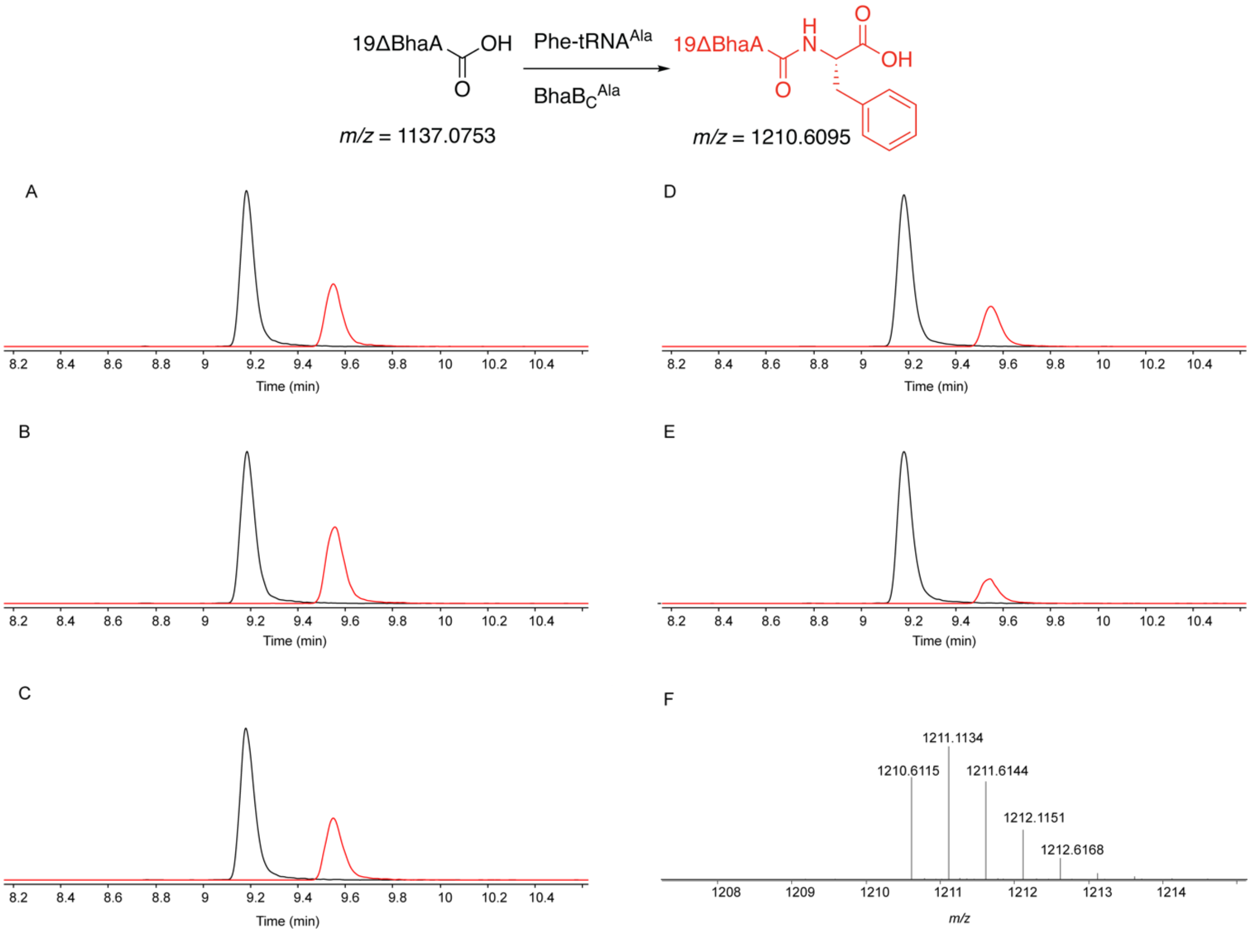

**Figure S10.** EICs of Δ19BhaA modified by $BhaB_C^{Ala}$ in the presence of *E. coli* $tRNA^{Ala}$ (UGC) charged with Trp. Each panel represents a $BhaB_C^{Ala}$ reaction when utilizing different amounts of Trp-$tRNA^{Ala}$. The concentration of the substrate peptide (Δ19BhaA) was 1.8 µM. The estimated concentration of Trp-$tRNA^{Ala}$ in each panel was as follows: A) 31 µM, B) 16 µM, C) 7.9 µM, D) 3.9 µM, E) 2.0 µM. The estimated reaction efficiency is as follows: A) 3%, B) 2%, C) 2.3%, D) 1.8%, E) 0.7%. The black trace is EIC of the substrate Δ19BhaA. The red trace is EIC of the product Δ19BhaA-Trp. F) The mass spectrum of the product. The calculated error comparing calculated and observed masses is shown in Table S6.

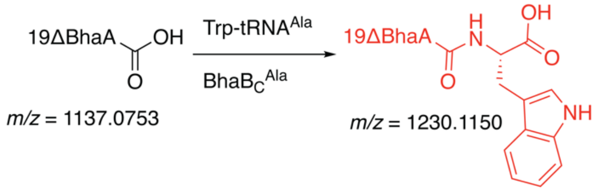

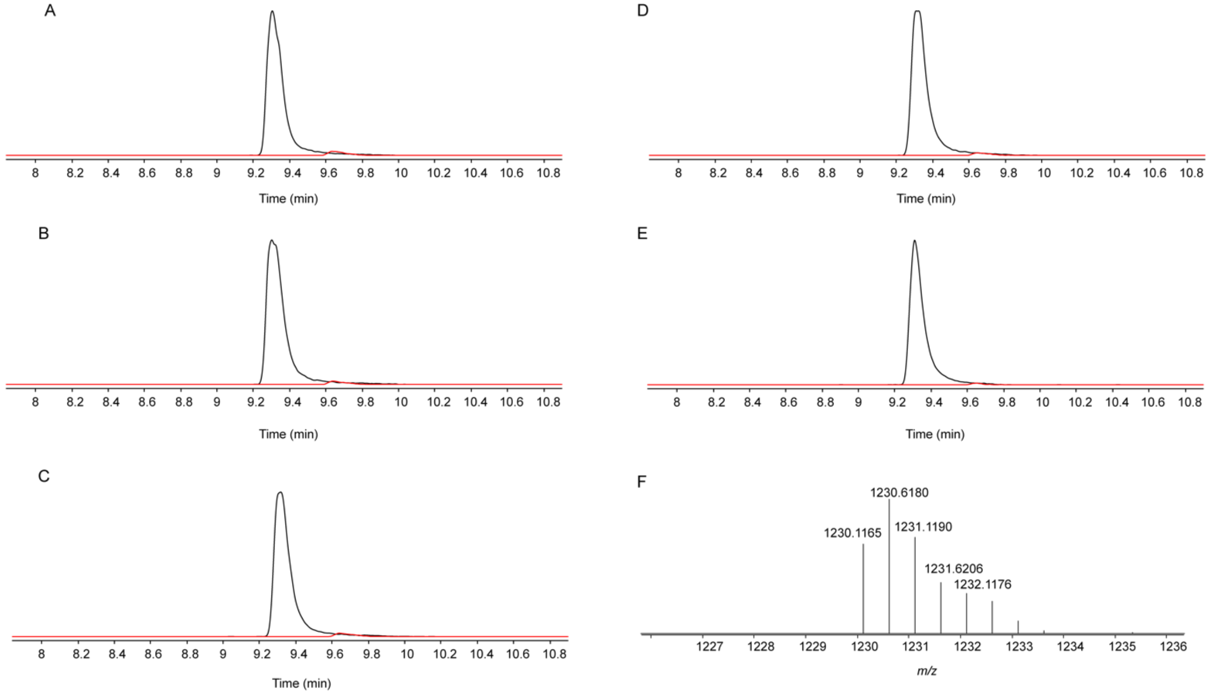

**Figure S11.** EICs of Δ19BhaA modified by $BhaB_C^{Ala}$ in the presence of *E. coli* $tRNA^{Ala}$ (UGC) charged with β-Ala. Each panel represents a $BhaB_C^{Ala}$ reaction when utilizing different amounts of (β-Ala)-$tRNA^{Ala}$. The concentration of the substrate peptide (Δ19BhaA) was 1.8 µM. The estimated concentration of (β-Ala)-$tRNA^{Ala}$ in each panel was as follows: A) 12 µM, B) 6 µM, C) 3 µM, D) 1.5 µM, E) 0.74 µM. The estimated reaction efficiency is as follows: A) 86%, B) 61%, C) 39%, D) 20%, E) 10%. The black trace is EIC of the substrate Δ19BhaA. The red trace is EIC of the product Δ19BhaA-(β-Ala). F) The mass spectrum of the product. The calculated error comparing calculated and observed masses is shown in Table S6.

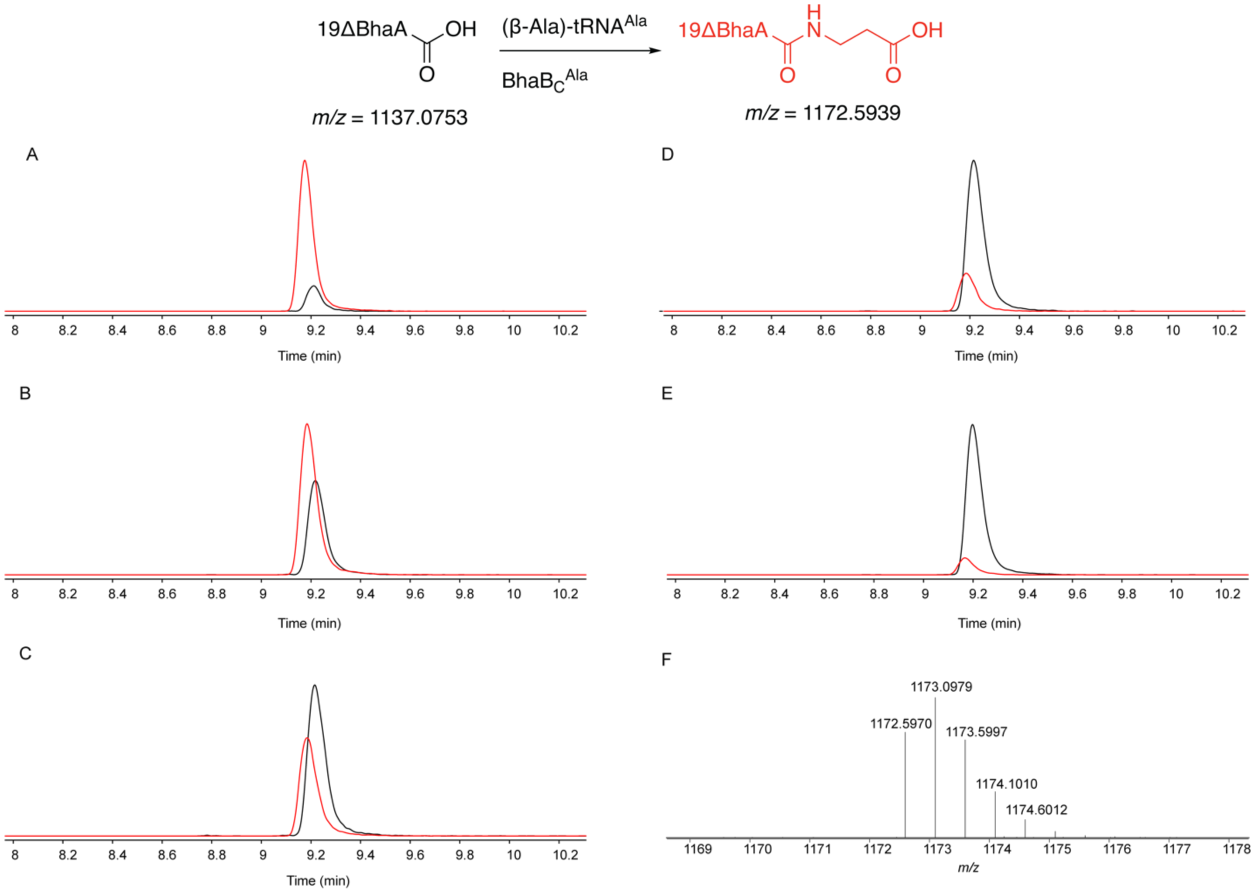

**Figure S12.** EICs of Δ19BhaA modified by BhaB$_C^{Ala}$ in the presence of *E. coli* tRNA$^{Ala}$ (UGC) charged with D-Ala. Each panel represents a BhaB$_C^{Ala}$ reaction when utilizing different amounts of (D-Ala)-tRNA$^{Ala}$. The concentration of the substrate peptide **(**Δ19BhaA) was 1.8 µM. The estimated concentration of (D-Ala)-tRNA$^{Ala}$ in each panel was as follows: A) 41 µM, B) 21 µM, C) 10 µM, D) 5.1 µM, E) 2.6 µM. The estimated reaction efficiency is as follows: A) 8.0%, B) 5.7%, C) 3.9%, D) 2.8%, E) 1.8%. The black trace is EIC of the substrate Δ19BhaA. The red trace is EIC of the product Δ19BhaA-(D-Ala). F) The mass spectrum of the product. The calculated error comparing calculated and observed masses is shown in Table S6.

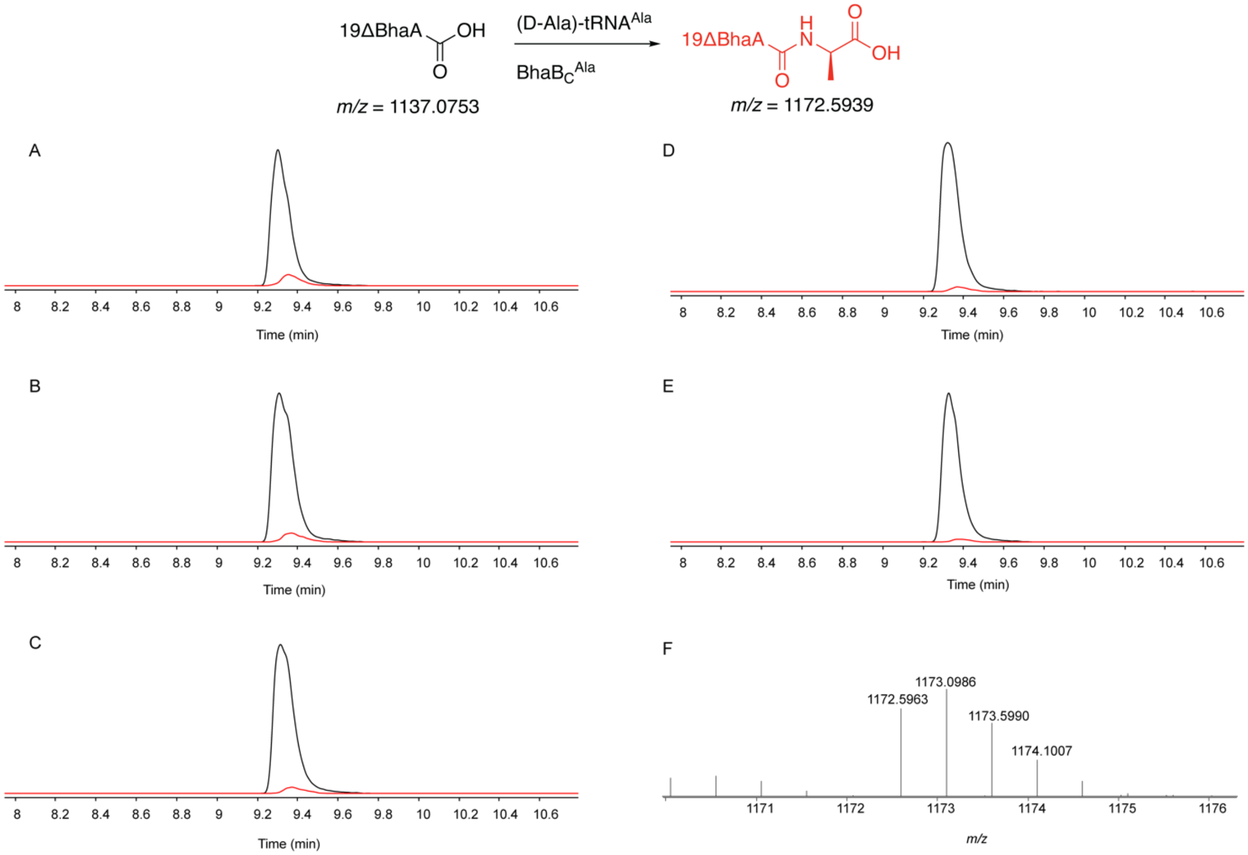

**Figure S13.** EICs of Δ19BhaA modified by $BhaB_C^{Ala}$ in the presence of *E. coli* $tRNA^{Ala}$ (UGC) charged with 2-aminoisobutyric acid (Aib). Each panel represents a $BhaB_C^{Ala}$ reaction when utilizing different amounts of Aib-$tRNA^{Ala}$. The concentration of the substrate peptide **(**Δ19BhaA) was 1.8 µM. The estimated concentration of Aib-$tRNA^{Ala}$ in each panel was as follows: A) 17 µM, B), 8.7 µM C) 4.3 µM, D) 2.2 µM, E) 1.1 µM. The product was not observed in any conditions. The black trace is the EIC of the substrate Δ19BhaA. The red trace is the EIC of the product Δ19BhaA-Aib.

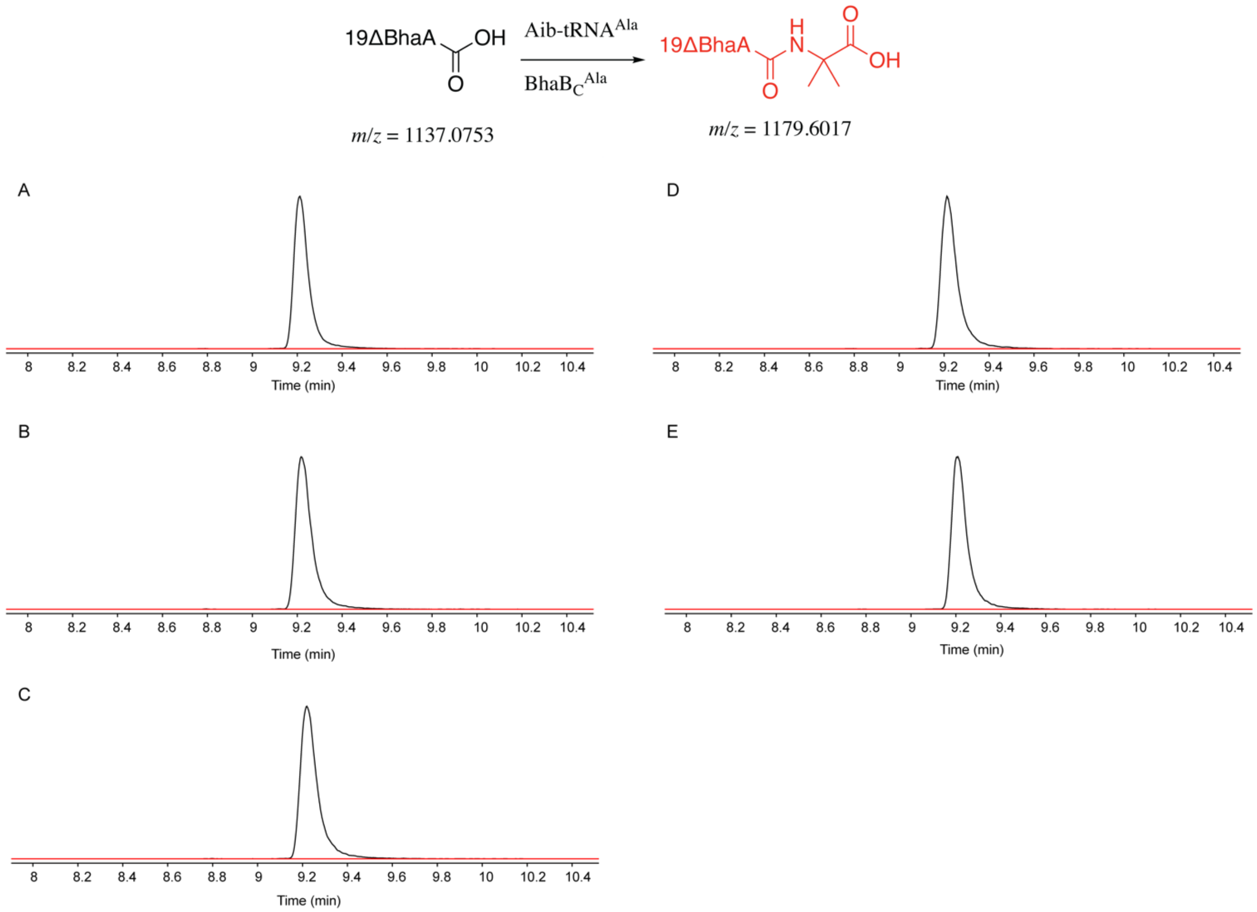

**Figure S14.** EICs of Δ19BhaA modified by $BhaB_C^{Ala}$ in the presence of *E. coli* $tRNA^{Ala}$ (UGC) charged with N-Me-Ala. Each panel represents a $BhaB_C^{Ala}$ reaction when utilizing different amounts of (N-Me-Ala)-$tRNA^{Ala}$. The concentration of the substrate peptide **(**Δ19BhaA) was 1.8 µM. The estimated concentration of (N-Me-Ala)-$tRNA^{Ala}$ in each panel was as follows: A) 40 µM, B) 20 µM, C) 10 µM, D) 5 µM, E) 2.5 µM. The product was not observed in any conditions. The black trace is the EIC of the substrate Δ19BhaA. The red trace is the EIC of the product Δ19BhaA-(N-Me-Ala).

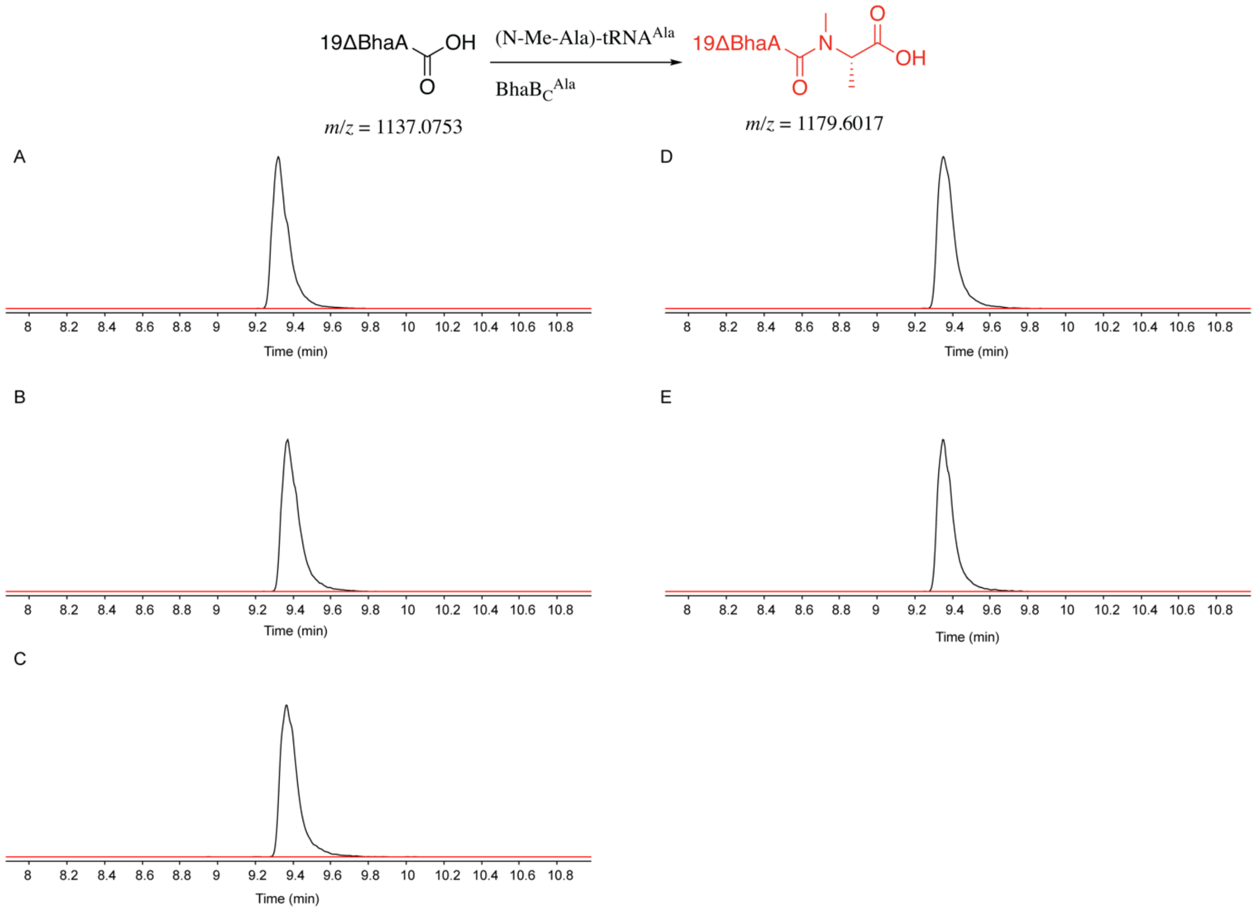

**Figure S15.** EICs of Δ19BhaA modified by $BhaB_C^{Ala}$ in the presence of *E. coli* $tRNA^{Ala}$ (UGC) charged with Lac. The concentration of the substrate peptide **(**Δ19BhaA) was 1.8 µM. Panels A–E show reactions with decreasing concentrations of Lac-$tRNA^{Ala}$, beginning with the highest concentration in Panel A and followed by two-fold serial dilutions through Panel E. The estimated reaction efficiency is as follows: A) 21%, B) 15%, C) 8.8%, D) 5.1%, E) 2%. The black trace is EIC of the substrate Δ19BhaA. The red trace is EIC of the product Δ19BhaA-Lac. F) The mass spectrum of the product. The calculated error comparing calculated and observed masses is shown in Table S6.

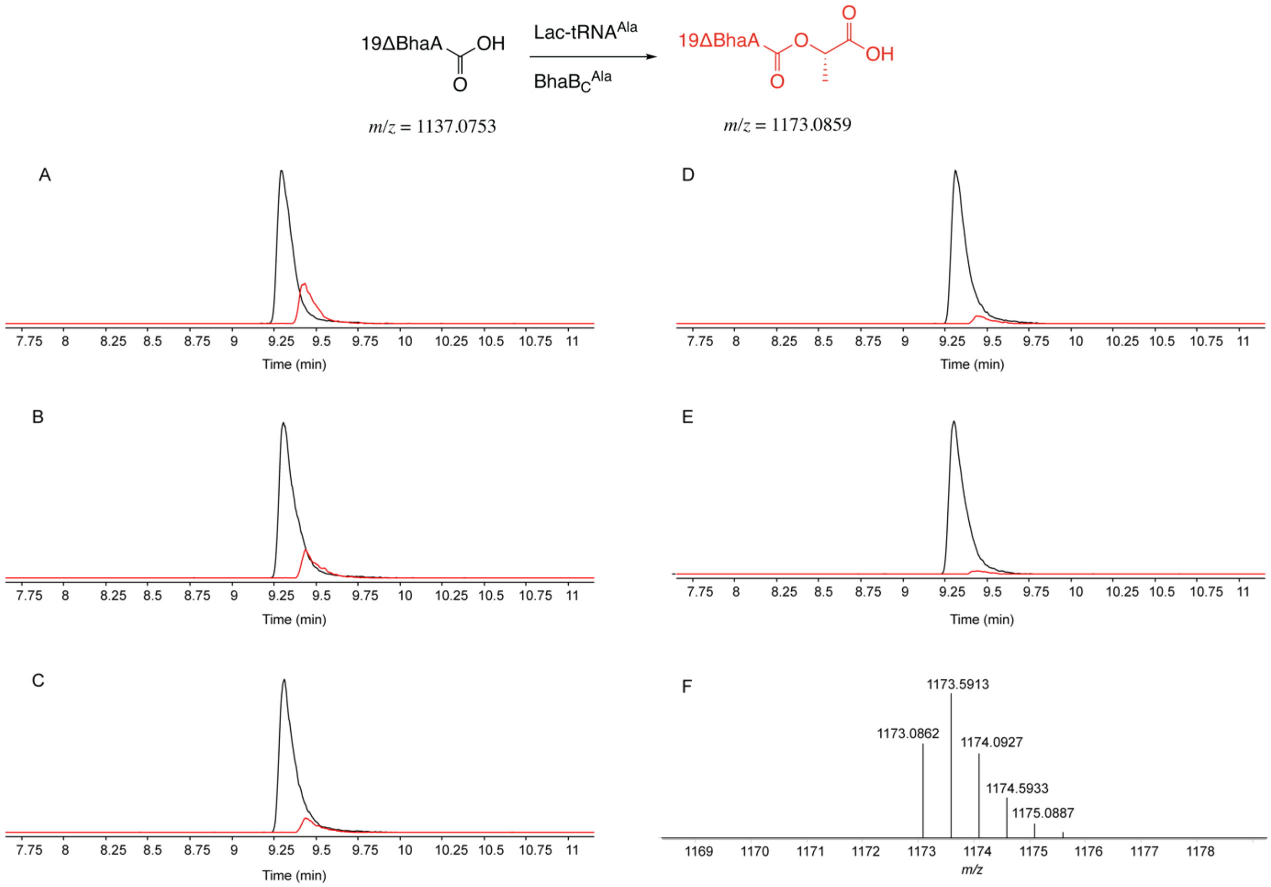

**Figure S16.** EICs of Δ19BhaA modified by $BhaB_C^{Ala}$ in the presence of *E. coli* $tRNA^{Ala}$ (UGC) charged with thioglycolic acid (ThioGly). The concentration of the substrate peptide **(**Δ19BhaA) was 1.8 µM. Panels A–E show reactions with decreasing concentrations of ThioGly-$tRNA^{Ala}$, beginning with the highest concentration in Panel A and followed by two-fold serial dilutions through Panel E. The estimated reaction efficiency is as follows: A) 3.9%, B) 1.9%, C) 0.80%, D) 0.42%, E) 0.25%. The black trace is EIC of the substrate Δ19BhaA. The blue trace is EIC of the product Δ19BhaA-ThioGly. F) The mass spectrum of Δ19BhaA-ThioGly. The calculated error comparing calculated and observed masses is shown in Table S6.

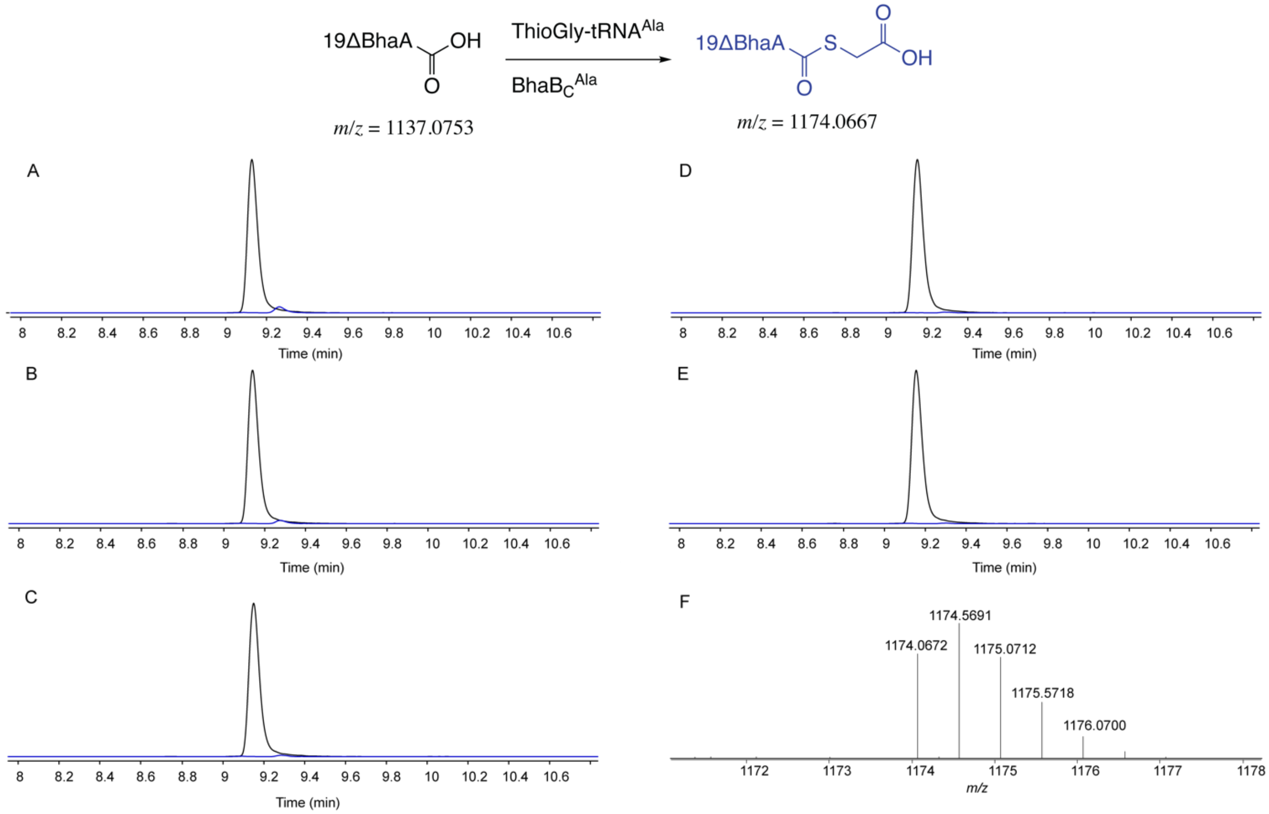

**Figure S17.** Native chemical ligation (NCL) of the peptide-thioester generated by BhaB$_C$$^{Ala}$ and ThioGly (Figure S16). EICs and mass spectra of products from BhaB$_C$$^{Ala}$-catalyzed reactions using ThioGly-tRNA$^{Ala}$ in the presence or absence of cysteamine. Cysteamine reacts with the thioester product of the BhaB$_C$$^{Ala}$ reaction via NCL.[20] The black trace corresponds to the EIC of the substrate (Δ19BhaA), the blue trace corresponds to the thioester product (Δ19BhaA–ThioGly), and the red trace corresponds to the NCL product (Δ19BhaA–cysteamine). The calculated error comparing calculated and observed masses is shown in Table S6.

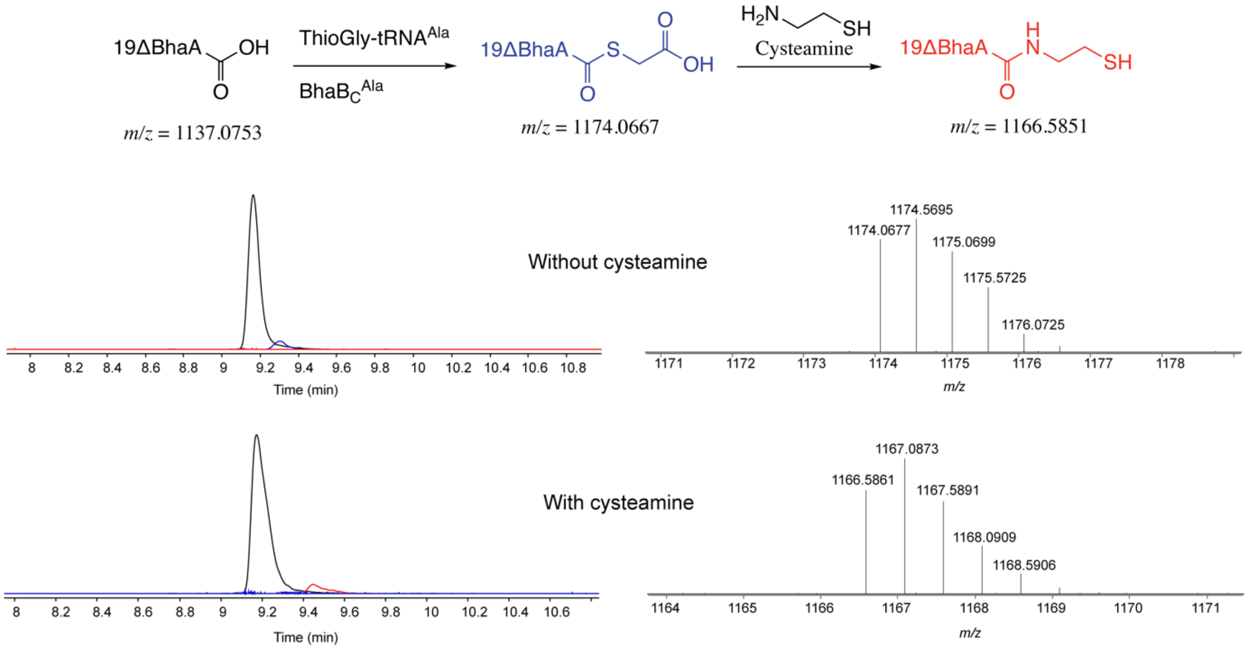

**Figure S18.** Δ19BhaA modified by $BhaB_C^{Ala}$ in the presence of *E. coli* $tRNA^{Gly}$(GCC) charged with Gly by *E. coli* GlyRS. Gly-$tRNA^{Gly}$ here was partially purified after GlyRS assay using the Monarch Spin RNA Cleanup Kit before being subjected to the $NaIO_4$-RNA extension assay and the $BhaB_C^{Ala}$ assay. A) PAGE result of the $NaIO_4$-RNA extension assay (refer to Figure S2). The estimated amount of aminoacylated tRNA (species b) compared to the total amount of RNA (species a and b) is 36%. B) EIC collected after the $BhaB_C^{Ala}$ reaction using 1.6 μM Δ19BhaA and 8.8 μM Gly-$tRNA^{Gly}$. C) The predicted clover leaf structures of *E. coli* $tRNA^{Gly}$ (GCC) using tRNAScanSE[21] and Forna.[22] D) Sequence alignments of *E. coli* $tRNA^{Ala}$ (UGC) and $tRNA^{Gly}$ (GCC).[23-25] The tRNA is colored by structural region: orange (acceptor stem), yellow (D-arm), green (anticodon arm), black (variable arm), and blue (T-stem).

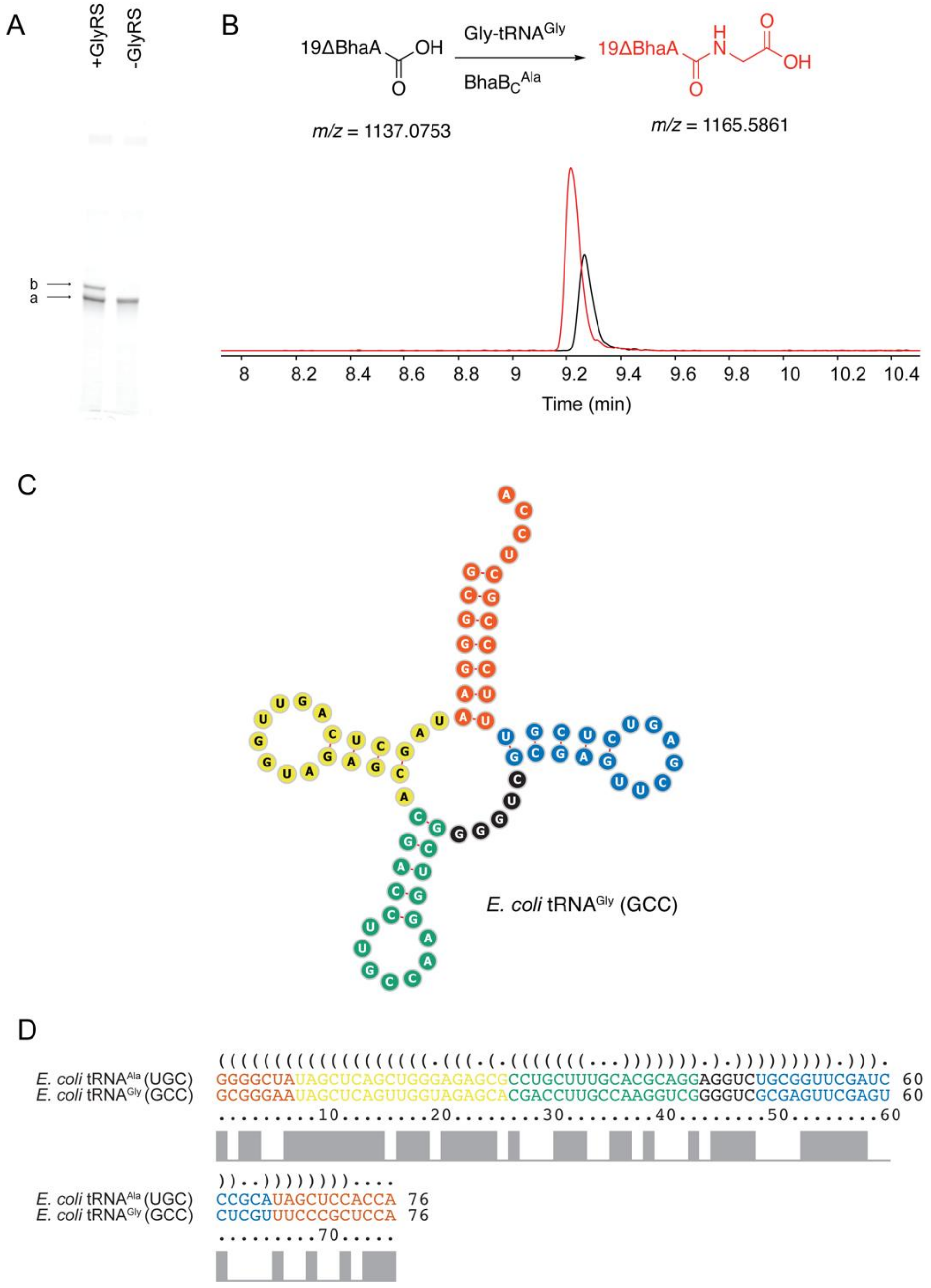

28

**Figure S19.** Predicted structures of examined tRNAs charged to Ala. Their sequences are provided in Table S3. The clover leaf structures of *E. coli* tRNA$^{Ala}$(UGC), *E. coli* tRNA$^{Trp}$, *T. bispora* tRNA$^{Glu}$ (CUC) and *P. syringae* tRNA$^{Cys}$ (GCA). The tRNA is colored by structural region: orange (acceptor stem), yellow (D-arm), green (anticodon arm), black (variable arm), and blue (T-stem). Sequence alignments comparing these tRNAs can be found in Figure S20. The structures were generated using tRNAScanSE[21] and Forna.[22]

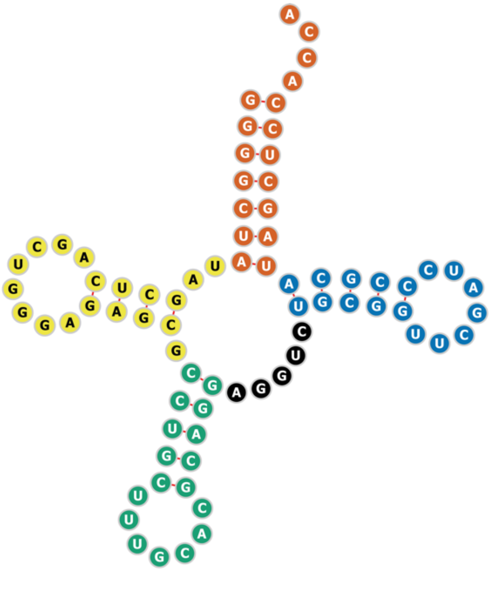

*E. coli* tRNA $^{Ala}$ (UGC)

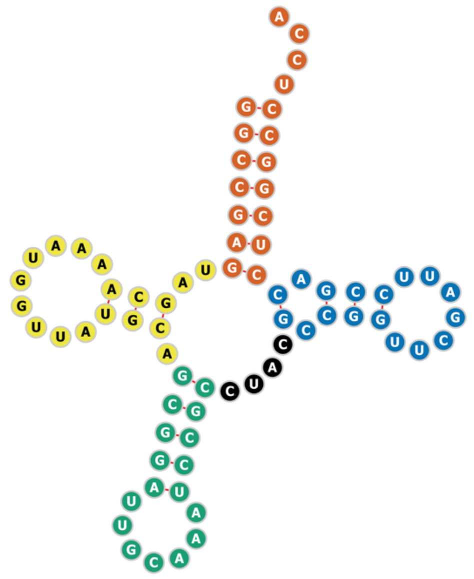

*P. syringae* tRNA$^{Cys}$ (GCA)

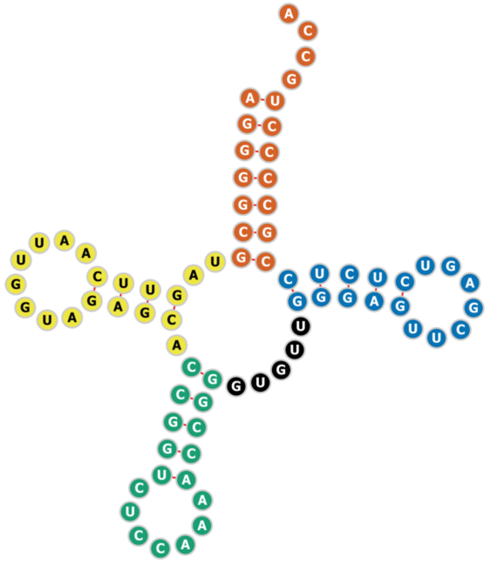

*E. coli* tRNA$^{Trp}$

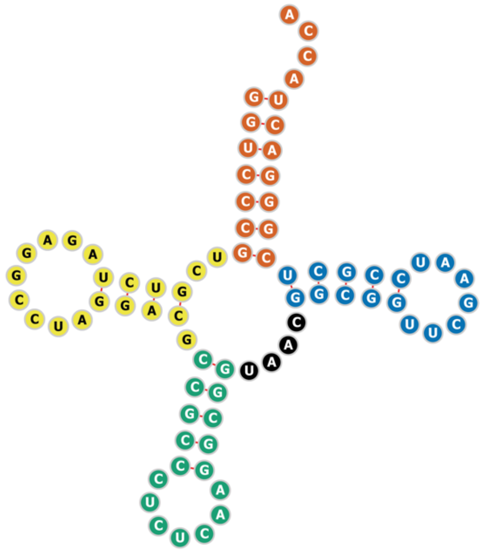

*T. bispora* tRNA$^{Glu}$ (CUC)

**Figure S20.** Sequence alignments of the tRNAs used to examine the Ala-tRNA specificity of $BhaB_C^{Ala}$. The boxed residues are the bases of the anticodon (green) and the discriminator bases (orange). Ec = *E. coli*, Ps = *P. syringae*, Tbi = *T. bispora*. The isoforms shown are $tRNA^{Cys}$(GCA), $tRNA^{Glu}$(CUC), and $tRNA^{Ala}$(UGC). The tRNA is colored by structural region: orange (acceptor stem), yellow (D-arm), green (anticodon arm), black (variable arm), and blue (T-stem) (see Figure S19). The alignments were performed using LocARNA.[23-25]

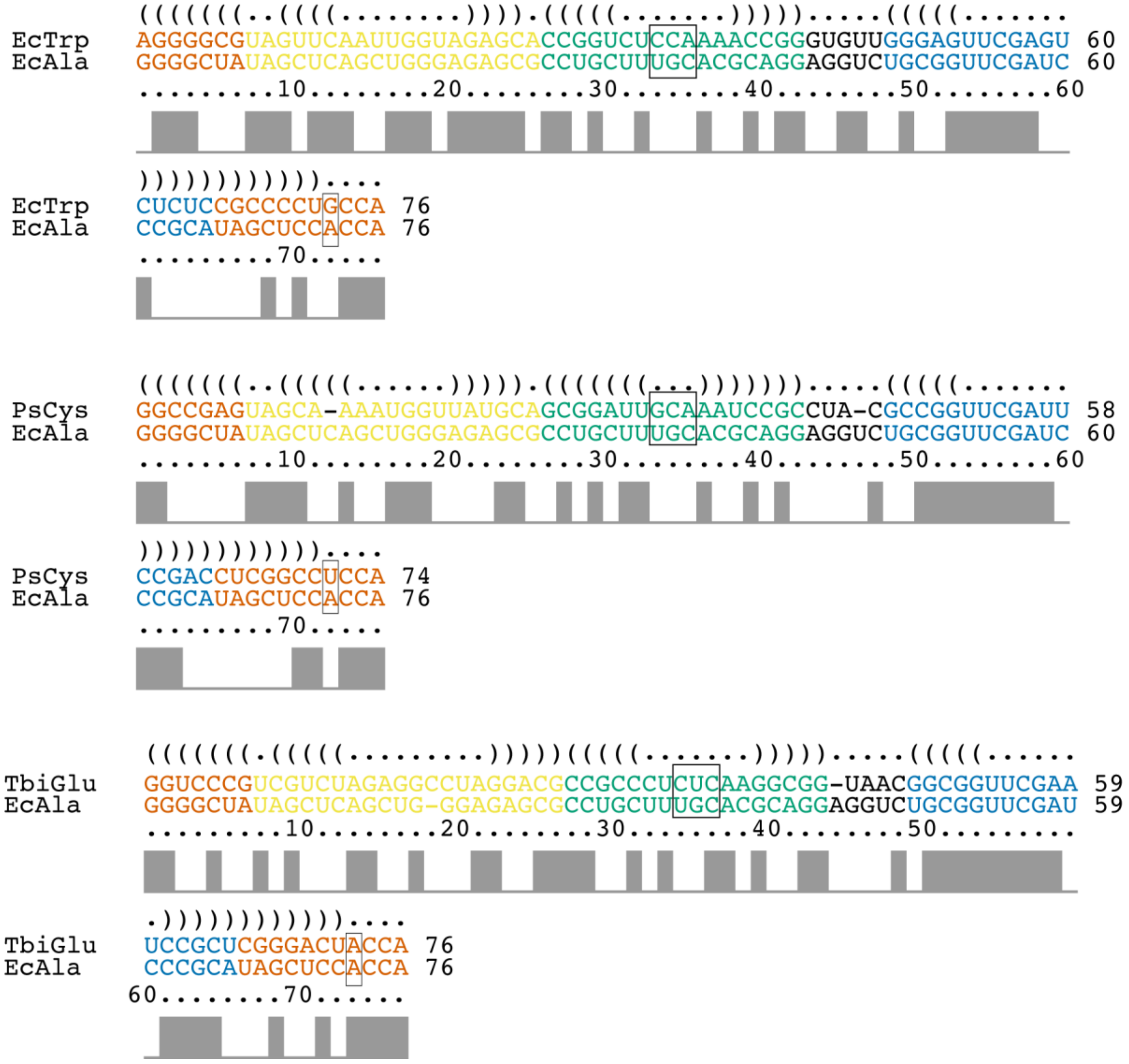

**Figure S21.** (A) PAGE analysis of the $NaIO_4$-RNA extension assay (Figure S2) with different tRNAs charged with Ala by flexizyme. The utilized primer sequence is provided in Table S4. R1, R2, and R3 represent the reactions depicted in Figure S2B. The upper bands represent species b (Figure S2) corresponding to aminoacylated tRNAs after being processed by the $NaIO_4$-RNA extension assay, and the lower bands represent species a (Figure S2) corresponding to free tRNAs after being processed by the $NaIO_4$-RNA extension assay. (B) The correspondence between gel lane numbers in panel A and the respective tRNA species, along with their estimated aminoacylation efficiencies. The sequences of each tRNA are provided in Table S3. Ec = *E. coli*, Ps = *P. syringae*, Tbi = *T. bispora*. The isoforms used were $tRNA^{Ala}$ (UGC), $tRNA^{Cys}$ (GCA) and $tRNA^{Glu}$ (CUC).

**A**

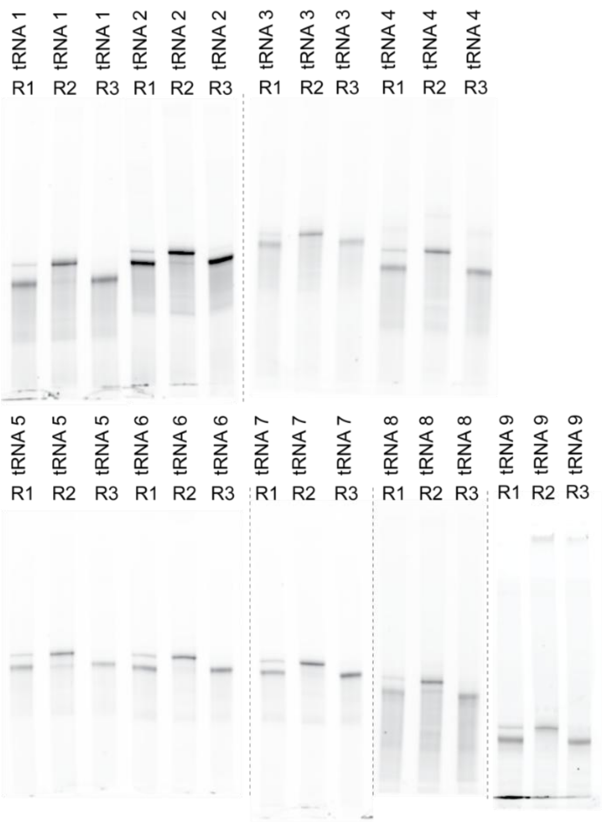

**B**

| RNA | tRNA | Estimated amino-acylation efficiency |
|---|---|---|
| tRNA 1 | Ec $tRNA^{Trp}$ | 23% |
| tRNA 2 | Ps $tRNA^{Cys}$ | 16% |
| tRNA 3 | Ec $tRNA^{Ala}$ grafted with Ps $tRNA^{Cys}$ anticodon arm | 26% |
| tRNA 4 | Ec $tRNA^{Ala}$ grafted with Ps $tRNA^{Cys}$ D-Arm | 30% |
| tRNA 5 | Tbi $tRNA^{Glu}$ | 29% |
| tRNA 6 | Ps $tRNA^{Cys}$ (C34G/A35C/U71A) | 31% |
| tRNA 7 | Ps $tRNA^{Cys}$ (U71A) | 30% |
| tRNA 8 | Ec $tRNA^{Ala}$ grafted with Ec $tRNA^{Trp}$ anticodon arm | 29% |
| tRNA 9 | Ec $tRNA^{Ala}$ grafted with Ps tRNA Cys T-Stem | 28% |

**Figure S22.** EICs of Δ19BhaA modified by $BhaB_C^{Ala}$ in the presence of *P. syringae* $tRNA^{Cys}$ (GCA) charged with Ala. Each panel represents the $BhaB_C^{Ala}$ reaction when utilizing different amounts of Ala-$tRNA^{Cys}$. The concentration of the substrate peptide (Δ19BhaA) was 1.8 µM. The estimated concentration of Ala-$tRNA^{Cys}$ in each panel was as follows: A) 24 µM, B) 12 µM, C) 5.9 µM, D) 2.9 µM, E) 1.5 µM. The estimated reaction efficiency is as follows: A) 2.6%, B) 1.8%, C) 1.2%, D) 0.87%, E) 0.62%. The black trace is the EIC of the substrate Δ19BhaA. The red trace is the EIC of the product Δ19BhaA-Ala. The predicted structure of the examined tRNA is shown in Figure 3 or Figure S19.

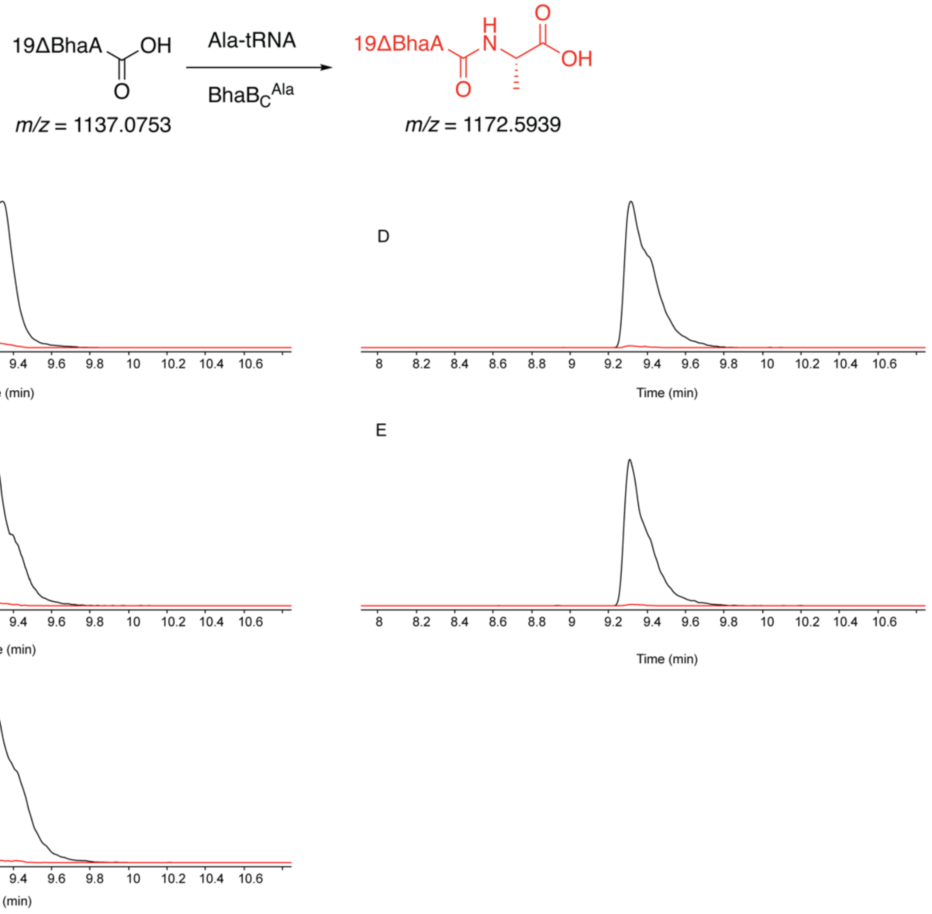

**Figure S23.** EICs of Δ19BhaA modified by $BhaB_C^{Ala}$ in the presence of *E. coli* $tRNA^{Trp}$ charged with Ala. Each panel represents a $BhaB_C^{Ala}$ reaction when utilizing different amounts of Ala-$tRNA^{Trp}$. The concentration of the substrate peptide (Δ19BhaA) was 1.8 µM. The estimated concentration of Ala-$tRNA^{Trp}$ in each panel was as follows: A) 33 µM, B) 16 µM, C) 8.2 µM, D) 4.1 µM, E) 2.1 µM. The estimated reaction efficiency is as follows: A) 5.9%, B) 4.0%, C) 2.7%, D) 1.9%, E) 1.4%. The black trace is EIC of the substrate Δ19BhaA. The red trace is EIC of the product Δ19BhaA-Ala. The predicted structure of the examined tRNA can be found in Figure 3 or Figure S19.

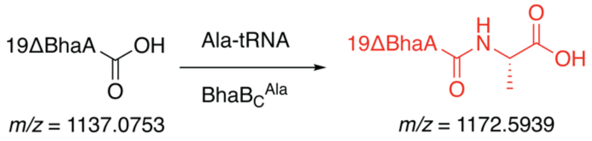

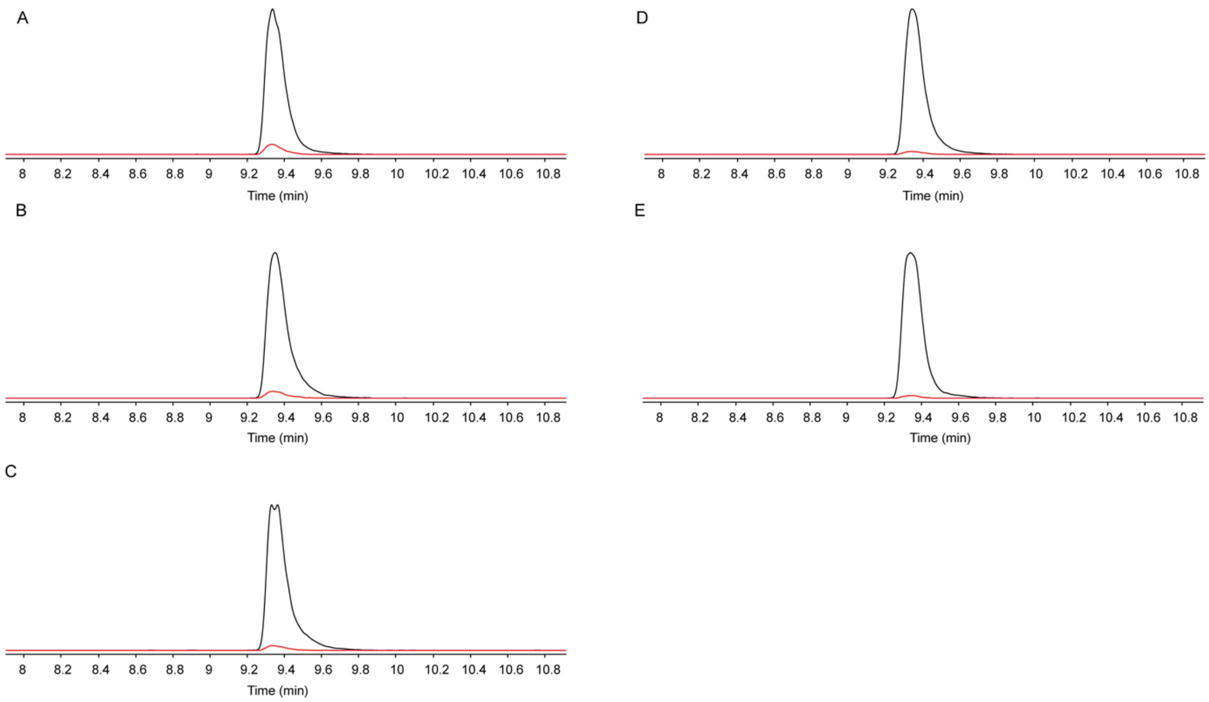

**Figure S24.** EICs of Δ19BhaA modified by $BhaB_C^{Ala}$ when reacting with Ala charged with *T. bispora* $tRNA^{Glu}$ (CUC). Each panel represents a $BhaB_C^{Ala}$ reaction when utilizing different amounts of Ala-$tRNA^{Glu}$. The concentration of the substrate peptide (Δ19BhaA) was 1.8 µM. The estimated concentration of Ala-$tRNA^{Glu}$ in each panel was as follows: A) 30 µM, B) 15 µM, C) 7.4 µM, D) 3.7 µM, E) 1.9 µM. The estimated reaction efficiency is as follows: A) 93%, B) 72%, C) 46%, D) 29%, E) 18%. The black trace is EIC of the substrate Δ19BhaA. The red trace is EIC of the product Δ19BhaA-Ala. The predicted structure of the examined tRNA can be found in Figure 3 or Figure S19.

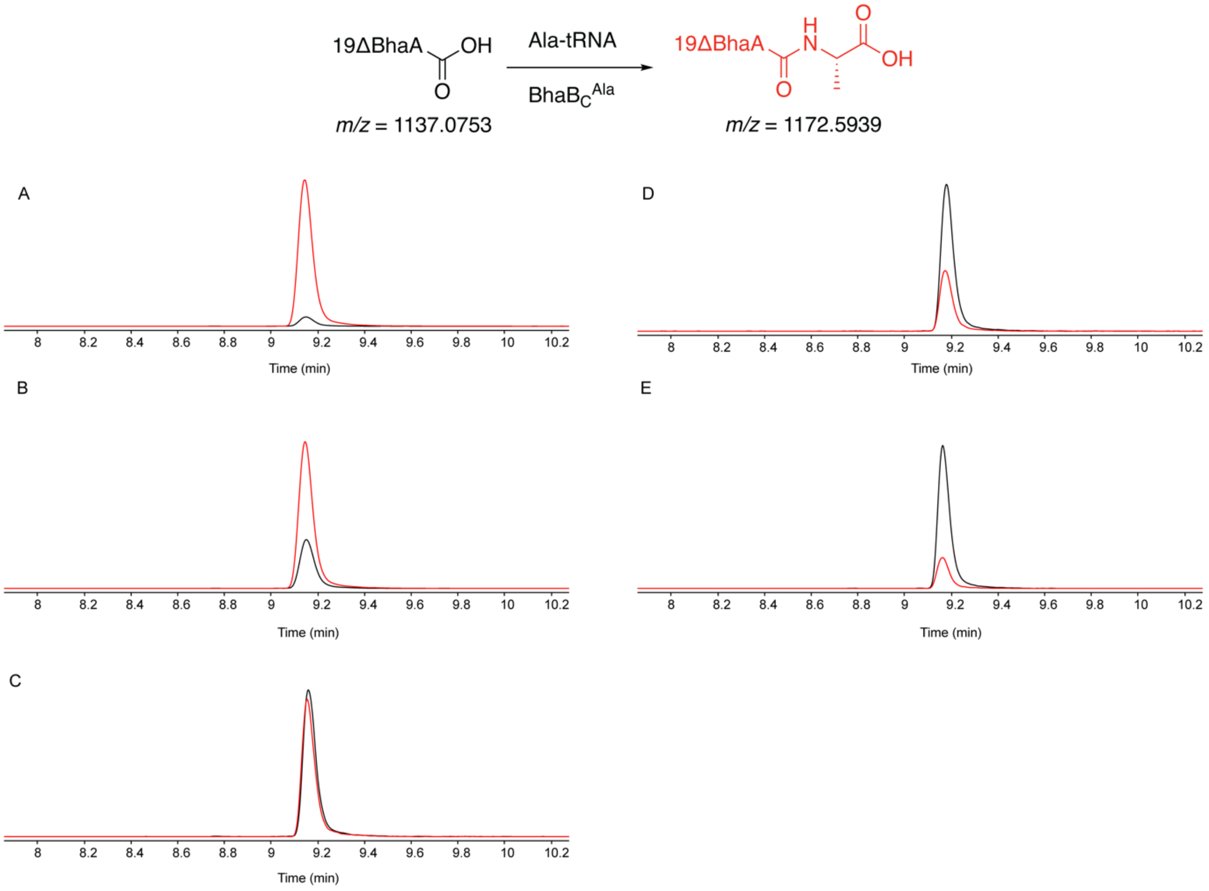

**Figure S25.** EICs of Δ19BhaA modified by $BhaB_C^{Ala}$ when reacting with Ala charged with *E. coli* $tRNA^{Ala}$ (UGC) grafted with the D-arm of *P. syringae* $tRNA^{Cys}$ ($tRNA^{Ala\text{-}Cys(D)}$). The sequence is provided in Table S3. Each panel represents a $BhaB_C^{Ala}$ reaction when utilizing different amounts of Ala-$tRNA^{Ala\text{-}Cys(D)}$. The concentration of the substrate peptide **(**Δ19BhaA) was 1.8 µM. The estimated concentration of Ala-$tRNA^{Ala\text{-}Cys(D)}$ in each panel was as follows: A) 30 µM, B) 15 µM, C) 7.6 µM, D) 3.8 µM, E) 1.9 µM. The estimated reaction efficiency is as follows: A) 99%, B) 98%, C) 73%, D) 35%, E) 17%. The black trace is EIC of the substrate Δ19BhaA. The red trace is EIC of the product Δ19BhaA-Ala. The predicted structure of the examined tRNA can be found in Figure 3.

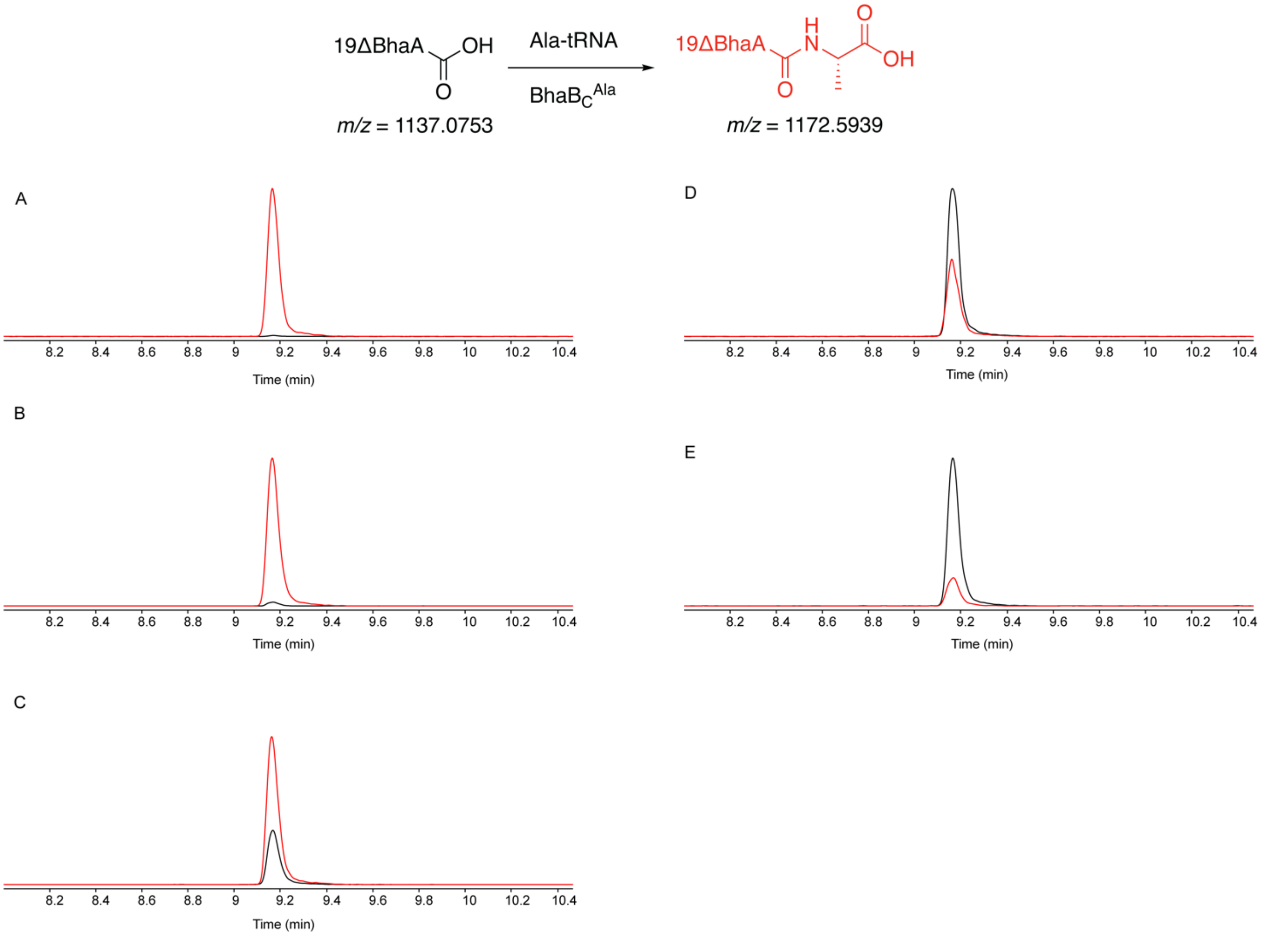

**Figure S26.** EICs of Δ19BhaA modified by $BhaB_C^{Ala}$ when reacting with Ala charged with *E. coli* $tRNA^{Ala}$ (UGC) grafted with the T-stem of *P. syringae* $tRNA^{Cys}$ ($tRNA^{Ala\text{-}Cys(T)}$). The sequence can is provided in Table S3. Each panel represents a $BhaB_C^{Ala}$ reaction when utilizing different amounts of Ala-$tRNA^{Ala\text{-}Cys(T)}$. The concentration of the substrate peptide (Δ19BhaA) was 1.8 µM. The estimated concentration of Ala-$tRNA^{Ala\text{-}Cys(T)}$ in each panel was as follows: A) 34 µM, B) 17 µM, C) 8.5 µM, D) 4.3 µM, E) 2.1 µM. The estimated reaction efficiency is as follows: A) 98%, B) 99%, C) 99%, D) 99%, E) 79%. The black trace is EIC of the substrate Δ19BhaA. The red trace is EIC of the product Δ19BhaA-Ala. The predicted structure of the examined tRNA can be found in Figure 3.

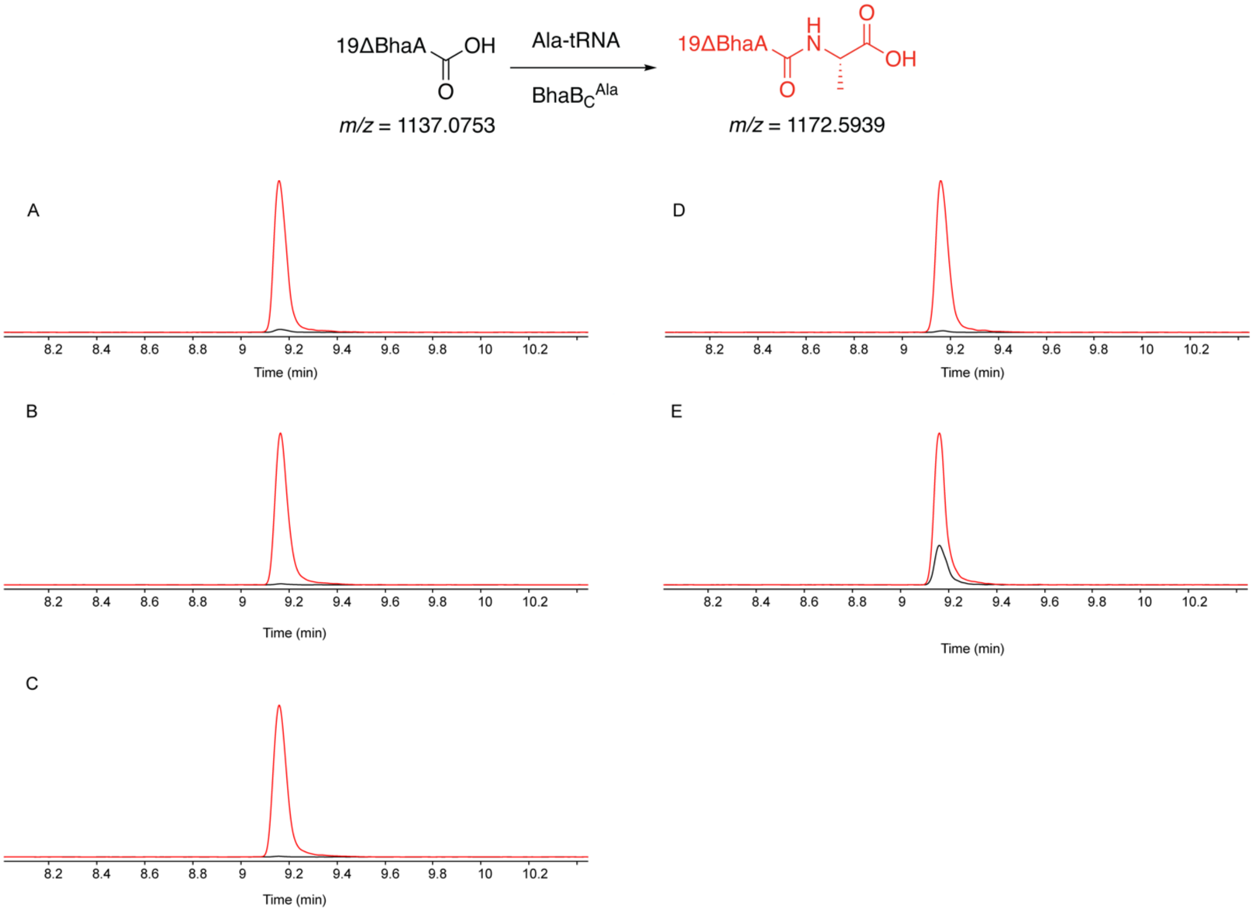

**Figure S27.** EICs of Δ19BhaA modified by $BhaB_C^{Ala}$ when reacting with Ala charged to *E. coli* $tRNA^{Ala}$ (UGC) grafted with the anticodon Arm of *P. syringae* $tRNA^{Cys}$ ($tRNA^{Ala\text{-}Cys(Anti)}$). The sequence is provided in Table S3. Each panel represents a $BhaB_C^{Ala}$ reaction when utilizing different amounts of Ala-$tRNA^{Ala\text{-}Cys(Anti)}$. The concentration of the substrate peptide (Δ19BhaA) was 1.8 μM. The estimated concentration of Ala-$tRNA^{Ala\text{-}Cys(Anti)}$ in each panel was as follows: A) 30 μM, B) 15 μM, C) 7.6 μM, D) 3.8 μM, E) 1.9 μM. The estimated reaction efficiency is as follows: A)33%, B) 13%, C) 4.5%, D) 2.3%, E) 1.8%. The black trace is EIC of the substrate Δ19BhaA. The red trace is EIC of the product Δ19BhaA-Ala. The predicted structure of the examined tRNA can be found in Figure 3.

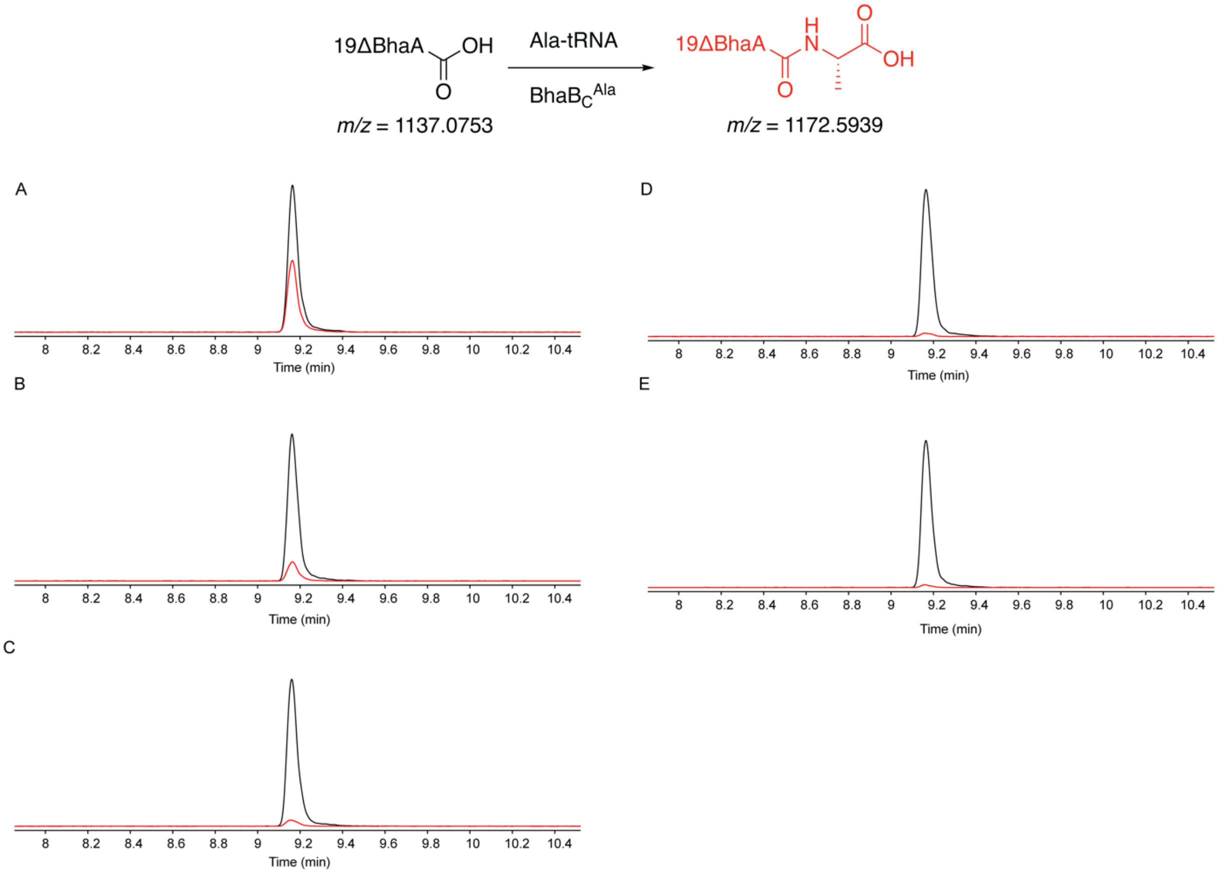

**Figure S28.** EICs of Δ19BhaA modified by $BhaB_C^{Ala}$ when reacting with Ala charged to *E. coli* $tRNA^{Ala}$ (UGC) grafted with the anticodon arm of *E. coli* $tRNA^{Trp}$ ($tRNA^{Ala\text{-}Trp(Anti)}$). The sequence is provided in Table S3. Each panel represents a $BhaB_C^{Ala}$ reaction when utilizing different amounts of Ala-$tRNA^{Ala\text{-}Trp(Anti)}$. The concentration of the substrate peptide (Δ19BhaA) was 1.8 µM. The estimated concentration of Ala-$tRNA^{Ala\text{-}Trp(Anti)}$ in each panel was as follows: A) 31 µM, B) 15 µM, C) 7.7 µM, D) 3.9 µM, E) 1.9 µM. The estimated reaction efficiency is as follows: A) 21%, B) 6.2%, C) 2.1%, D) 1.2%, E) 0.78%. The black trace is EIC of the substrate Δ19BhaA. The red trace is EIC of the product Δ19BhaA-Ala. F) The predicted structure of the examined tRNA.

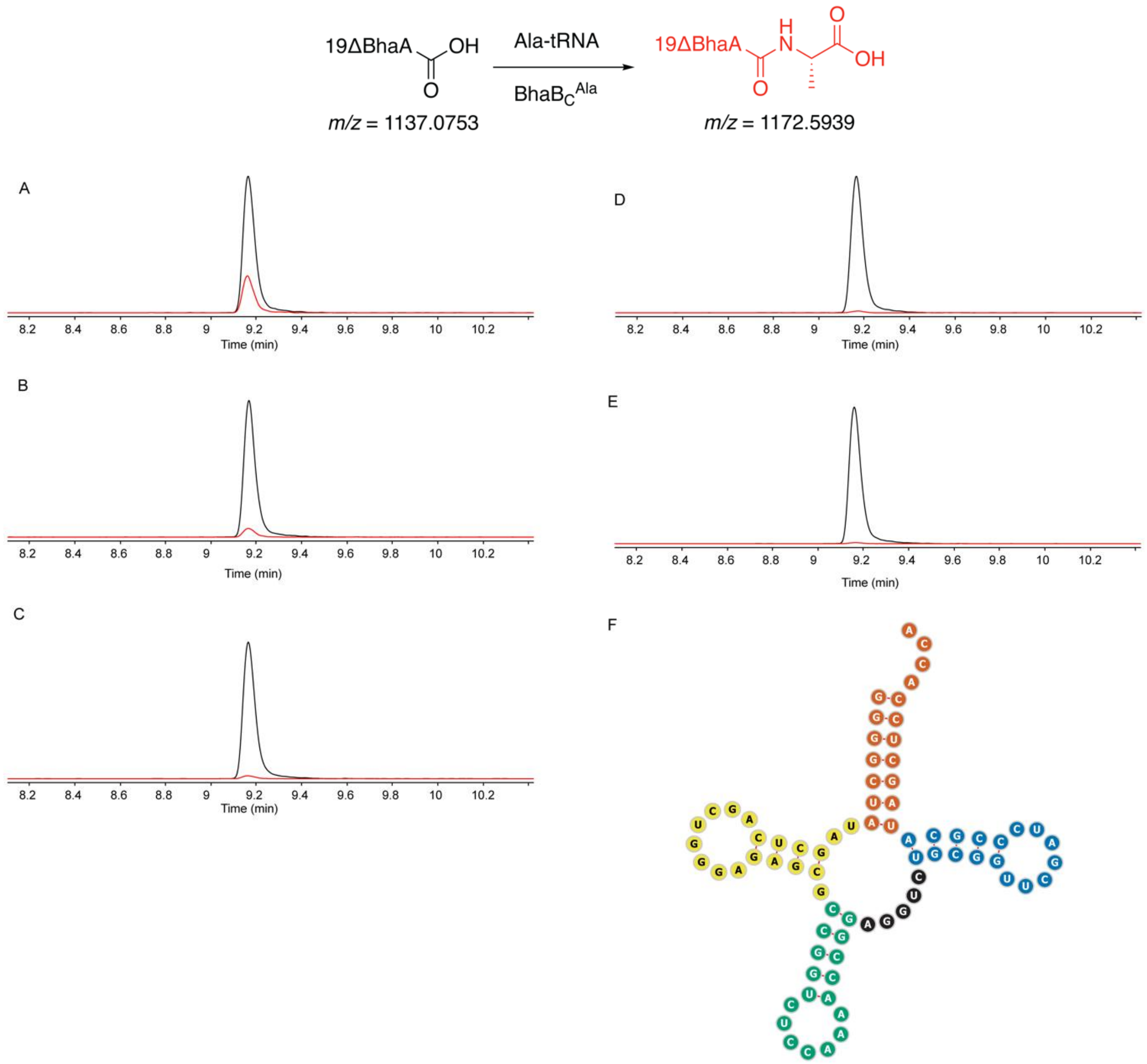

**Figure S29.** EICs of Δ19BhaA modified by $BhaB_C^{Ala}$ when reacting with Ala charged onto *P. syringae* $tRNA^{Ala}$ (GCA) (U71A). Each panel represents a $BhaB_C^{Ala}$ reaction when utilizing different amounts of Ala-tRNA. The concentration of the substrate peptide (Δ19BhaA) was 1.8 µM. The estimated concentration of Ala-tRNA in each panel was as follows: A) 30 µM, B) 15 µM, C) 7.6 µM, D) 3.8 µM, E) 1.9 µM. The estimated reaction efficiency is as follows: A) 47%, B) 27%, C) 14%, D) 9.4%, E) 6.4%. The black trace is EIC of the substrate Δ19BhaA. The red trace is EIC of the product Δ19BhaA-Ala. The predicted structure of the examined tRNA can be found in Figure 3.

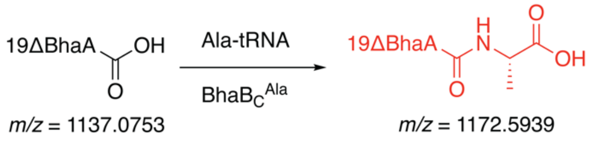

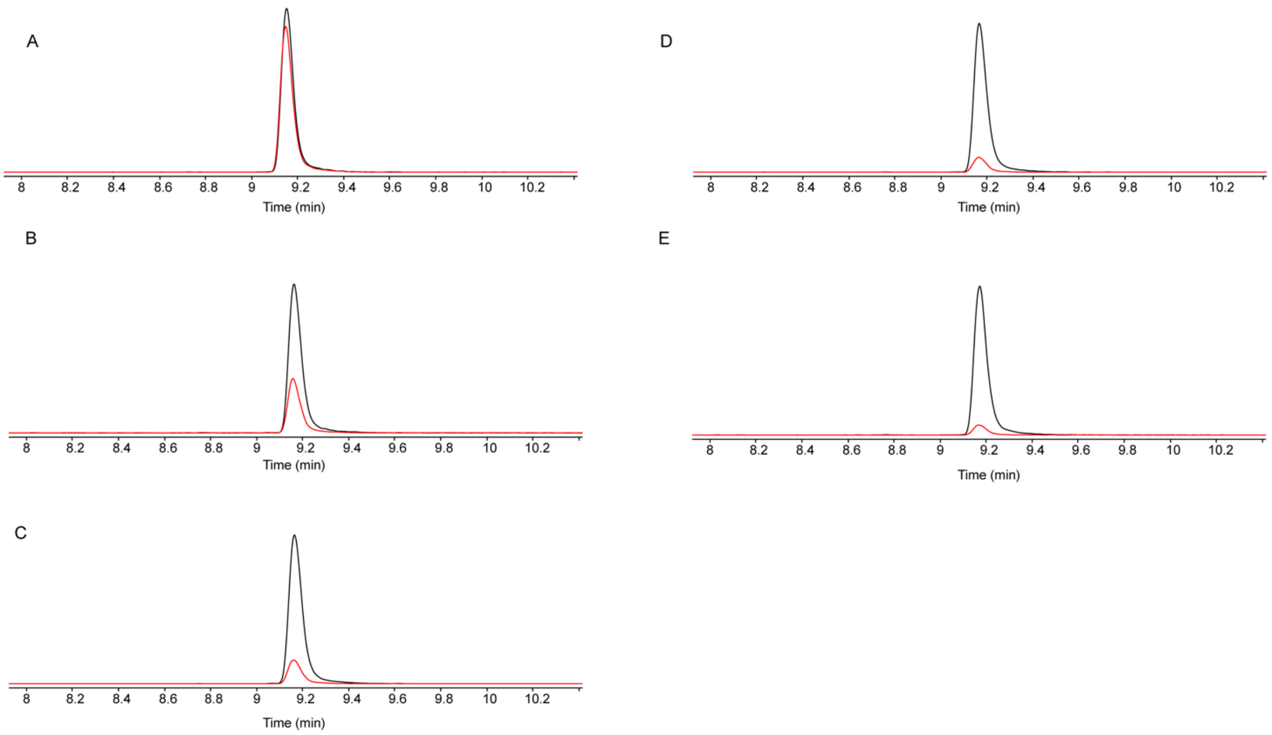

**Figure S30.** EICs of Δ19BhaA modified by $BhaB_C^{Ala}$ when reacting with Ala charged with *P. syringae* $tRNA^{Ala}$ (GCA) (C34G/A35C/U71A). The discriminator base and the anticodon of this non-cognate tRNA match those of *E. coli* $tRNA^{Ala}$ (GGC). Each panel represents a $BhaB_C^{Ala}$ reaction when utilizing different amounts of Ala-tRNA. The concentration of the substrate peptide (Δ19BhaA) was 1.8 μM. The estimated concentration of Ala-tRNA in each panel was as follows: A) 30 μM, B) 15 μM, C) 7.5 μM, D) 3.8 μM, E) 1.9 μM. The estimated reaction efficiency is as follows: A) 85%, B) 57%, C) 34%, D) 20%, E) 13%. The black trace is EIC of the substrate Δ19BhaA. The red trace is EIC of the product Δ19BhaA-Ala. The predicted structure of the examined tRNA can be found in Figure 3.

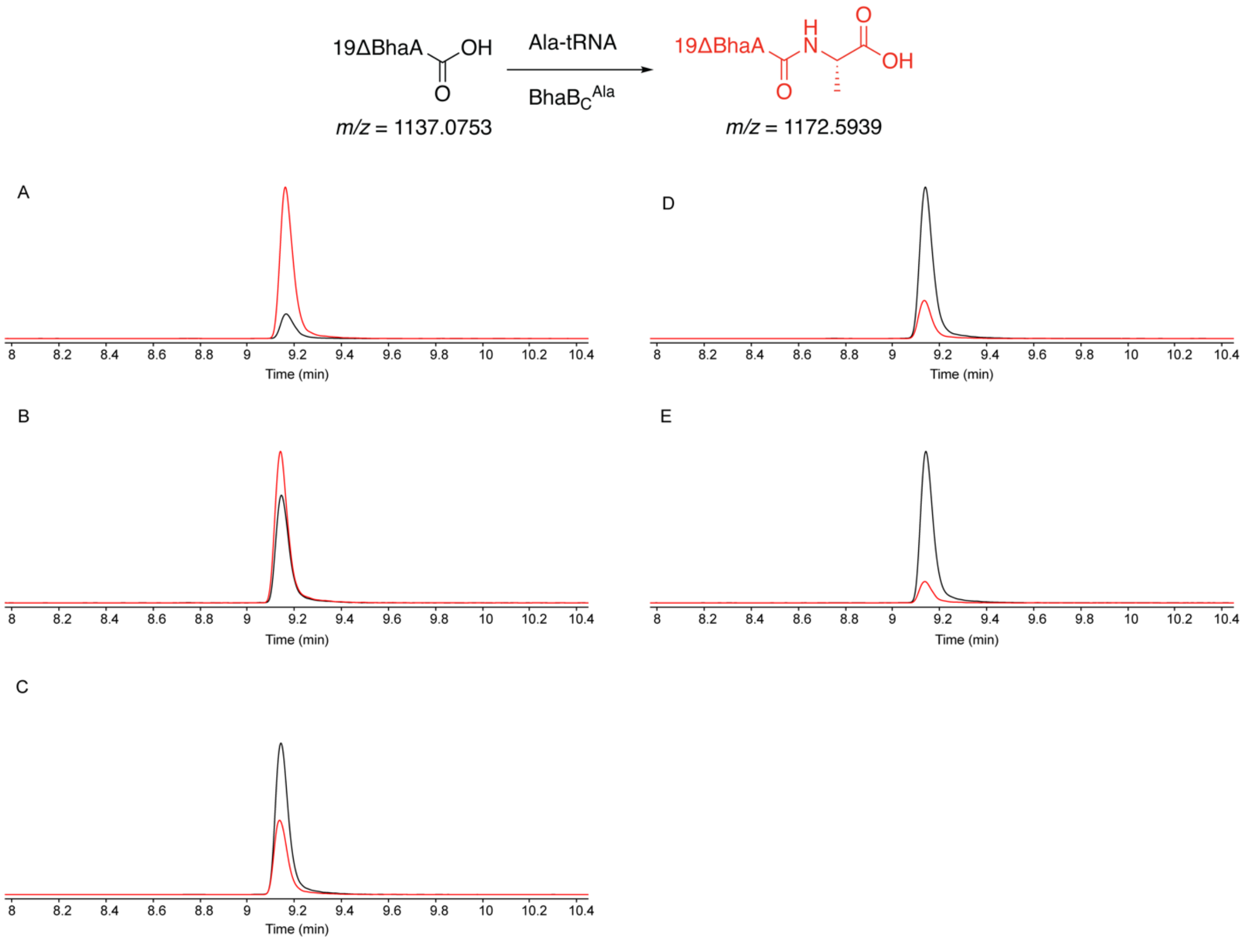

**Figure S31.** Alphafold3 model of the $BhaB_C^{Ala}$-BhaA-tRNA complex. A) The structure includes $BhaB_C^{Ala}$, BhaA, *E. coli* $tRNA^{Ala}$ (UGC), $Mg^{2+}$, and ATP. A) The tRNA is colored by structural region: orange (acceptor stem), yellow (D-arm), green (anticodon arm), black (variable arm), and blue (T-arm). The figure was made using Chimera.[26] The tRNA cloverleaf structure was generated using tRNAScanSE[21] and Forna.[22] B) Alphafold3 model colored by predicted local distance difference test (pLDDT) scores (x); C) Predicted aligned error (pAE) plot for the model. The model file can be found in Supplementary Data 1.

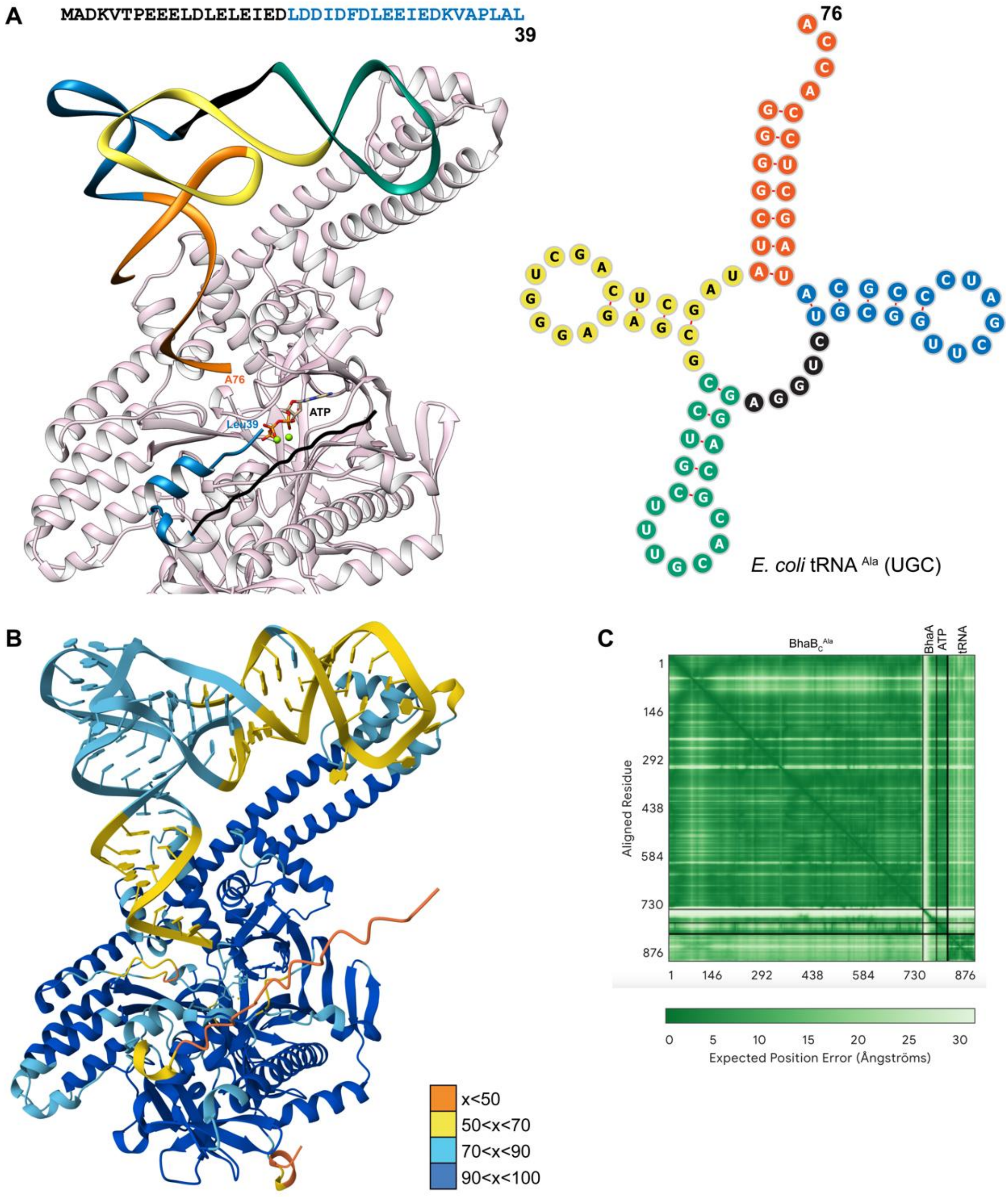

**Figure S32.** An active site view of the Alphafold3 model of BhaA, $BhaB_C^{Ala}$, ATP, $Mg^{2+}$, and *E. coli* $tRNA^{Ala}$(UGC) depicting that the first 19 residues of BhaA do not interact with $BhaB_C^{Ala}$. The peptide sequence is displayed above the model, with colors corresponding to those used in the AlphaFold3 model. The depicted Leu39 is the C-terminal residue of BhaA. The figure was made using Chimera.[26]

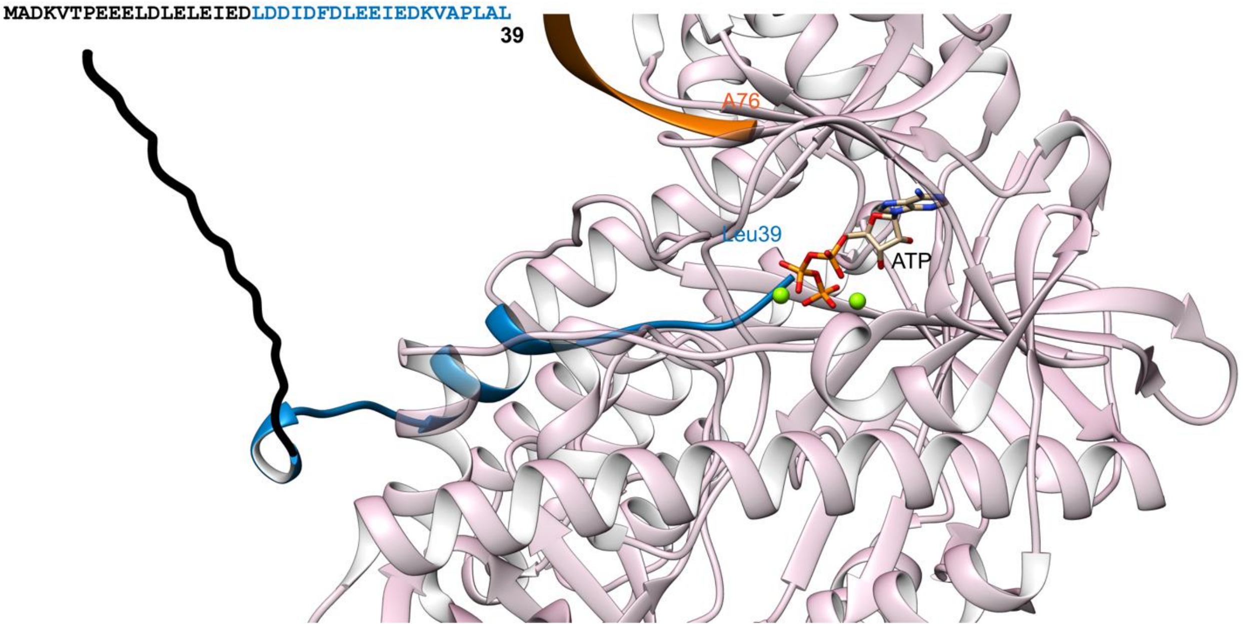

**Figure S33.** (A) PAGE analysis of RNAs used in this study. The ladder is low range ssRNA ladder (NEB). (B) The correspondence between indicated gel lane numbers and the respective tRNA species. The sequences of each tRNA are provided in Table S3. Ec = *E. coli*, Ps = *P. syringae*, Tbi = *T. bispora*. $tRNA^{Ala}$ here is $tRNA^{Ala}$ (UGC), tRNACys here is $tRNA^{Cys}$ (GCA), and $tRNA^{Glu}$ here is $tRNA^{Glu}$ (CUC).

**A**

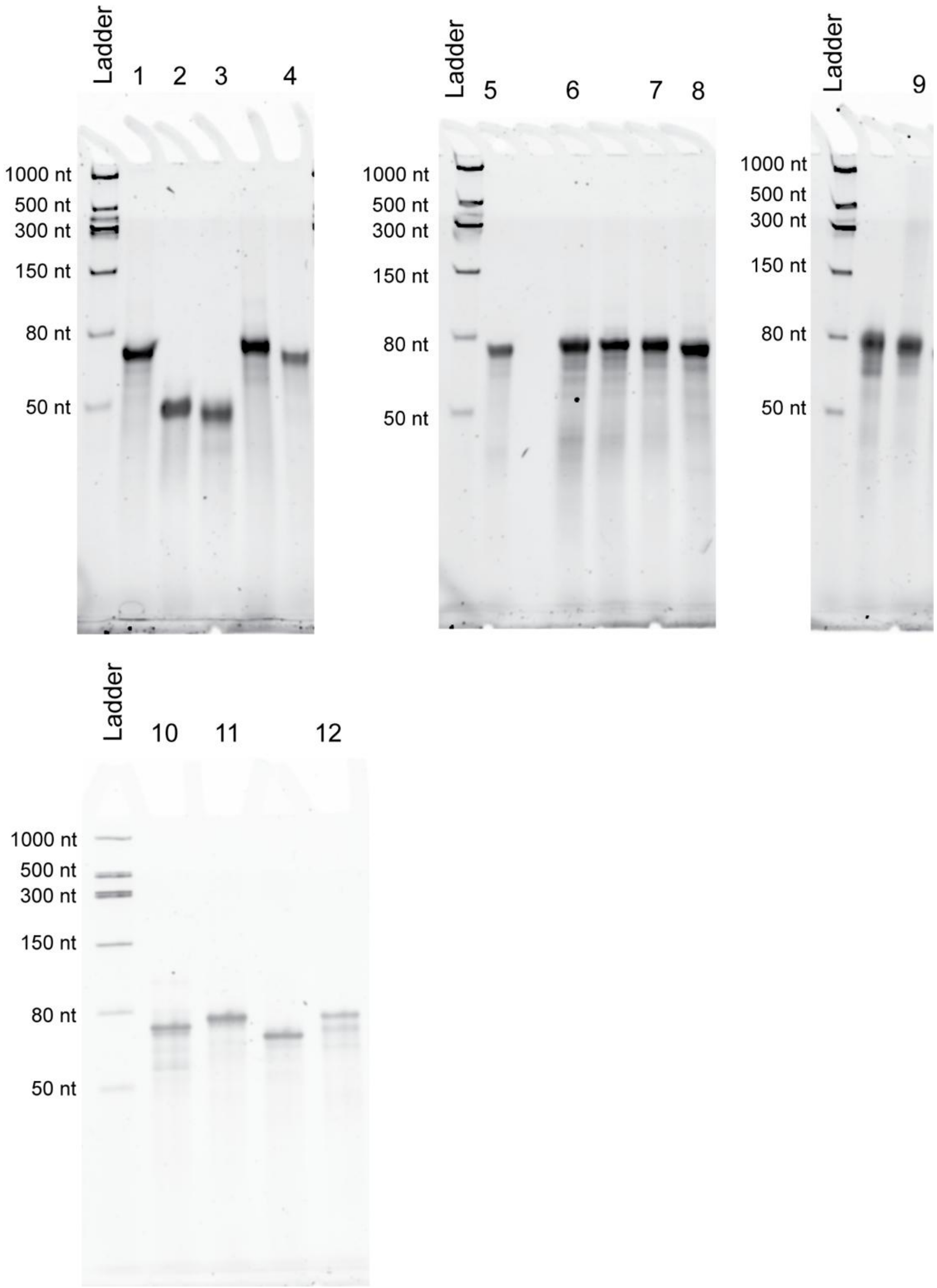

**B**

| Number | RNA |
|---|---|
| 1 | Ec $tRNA^{Ala}$ |
| 2 | dFx |
| 3 | eFx |
| 4 | Ec $tRNA^{Ala}$ grafted with Ec $tRNA^{Trp}$ Anticodon Arm |
| 5 | Ps $tRNA^{Cys}$ |
| 6 | Ps $tRNA^{Cys}$ (U71A) |
| 7 | Ps $tRNA^{Cys}$ (C34G/A35C/U71A) |
| 8 | Tbi $tRNA^{Glu}$ |
| 9 | Ec $tRNA^{Trp}$ |
| 10 | Ec $tRNA^{Ala}$ grafted with Ps $tRNA^{Cys}$ D-Arm |
| 11 | Ec $tRNA^{Ala}$ grafted with Ps $tRNA^{Cys}$ T-Stem |
| 12 | Ec $tRNA^{Ala}$ grafted with Ps $tRNA^{Cys}$ Anticodon Arm |

**Table S1.** Summary of BhaB$_C$$^{Ala}$ activity using non-cognate tRNAs charged with L-Ala. The EICs are shown in Figures S22-S30. Ec = *E. coli*, Ps = *P. syringae*, Tbi = *T. bispora*. tRNA$^{Ala}$ here is tRNA$^{Ala}$ (UGC), tRNA$^{Cys}$ is tRNA$^{Cys}$ (GCA), and tRNA$^{Glu}$ is tRNA$^{Glu}$ (CUC).

| **Ala-tRNA examined** | **Estimated [Ala-tRNA], reaction efficiency** | | | | |
|---|---|---|---|---|---|
| Ala-Ps tRNA$^{Cys}$ | 24 µM, 2.6% | 12 µM, 1.8% | 5.9 µM, 1.2% | 2.9 µM, 0.87% | 1.5 µM, 0.62% |
| Ala-Ps tRNA$^{Cys}$ (U71A) | 30 µM, 47% | 15 µM, 27% | 7.6 µM, 14% | 3.8 µM, 9.4% | 1.9 µM, 6.4% |
| Ala-Ps tRNA$^{Cys}$ (C34G/A35C/U71A) | 30 µM, 85% | 15 µM, 57% | 7.5 µM, 34% | 3.8 µM, 20% | 1.9 µM, 13% |
| Ala-Ec tRNA$^{Ala}$ grafted with Ps tRNA$^{Cys}$ D-Arm | 30 µM, 99% | 15 µM, 98% | 7.6 µM, 73% | 3.8 µM, 35% | 1.9 µM, 17% |
| Ala-Ec tRNA$^{Ala}$ grafted with Ps tRNA$^{Cys}$ T-Stem | 34 µM, 98% | 17 µM, 99% | 8.5 µM, 99% | 4.3 µM, 99% | 2.1 µM, 79% |
| Ala-Ec tRNA$^{Ala}$ grafted with Ps tRNA$^{Cys}$ Anticodon Arm | 30 µM, 33% | 15 µM, 13% | 7.6 µM, 4.5% | 3.8 µM, 2.3% | 1.9 µM, 1.8% |
| Ala-Ec tRNA$^{Trp}$ | 33 µM, 5.9% | 16 µM, 4.0% | 8.2 µM, 2.7% | 4.1 µM, 1.9% | 2.1 µM, 1.4% |
| Ala-Ec tRNA$^{Ala}$ grafted with Ec tRNA$^{Trp}$ Anticodon Arm | 31 µM, 21% | 15 µM, 6.2% | 7.7 µM, 2.1% | 3.9 µM, 1.2% | 1.9 µM, 0.78% |
| Ala-Tbi tRNA$^{Glu}$ | 30 µM, 93% | 15 µM, 72% | 7.4 µM, 46% | 3.7 µM, 29% | 1.9 µM, 18% |

**Table S2.** Sequence of the plasmid used to express $BhaB_C^{Ala}$ (NCBI accession: BAB05753.1). *E. coli* codon-optimized DNA sequence encoding $BhaB_C^{Ala}$ is highlighted in red.

taatacgactcactataggggaattgtgagcggataacaattcccctctagaaataattttgtttaactttaag
aaggagatataccatgggcagcagccatcatcatcatcatcacagcagcggcctggtgccgcgcggcagccata
tgtggttcaatacttccacatggcgttggttaaagccatttgtgctccgcagtacctctatccccatcgagatg
gttttgaacttaaggttaaaagaatcaataaagcaactaaatgaatggagcaaaagccaacaatcatttgatca
atggtgcaaccattggattaaagaaattaggtctgtaccagtagagcgaataagtcttccgcaccaacgaaaag
tcatacgaagaataaaaaaaagaaacccactaaccgaacaagataaaatcattgttaccaagtggggaatggaa
aagttgattgatgagtactgctcacagcagcaatcattaataagcttaaagaaaaaggcgcaacaatcttttca
tgaagagtttgaacgactacaacaacagttaattgagaggtttcaatctagccaatatgaattagctctgtata
cattgaaccacaagctttggcatttctggattaaggagaatcgtcttgaaaatctaactgatgcacaaaaaaaa
caaacatgtagaacccttttcgcctatcttcaacgggtctcaactaaaaatgacactatcggtgaatatggccc
tatcagttatggtgcattcaatcaaggcaagctgattcaaactaggcaaattacgaaaagaaaagcctatttttg
cctatcaagggctacaaaaattgctaagtgttataaaaaaggatttaatgccgcaggctcatgcctggaatctc
ccatcgtccacggtagatgggtggaagcttgttagagaacgggttgaagattgtaatgatcctcgttcagaaat
atggatacgaatcctagatcaacttgaggagactagacatttgtttgagacgtccactcaaccaaacgaaaaag
aacaagccttagctcaagctgagaaactatttacaaacataacaggggaggcttttcaaggagagaatcagcaa
tacttcgcagataaaacattgctgtatgaagaatgccatgatcagcaatatttacccgtcaccctatctgtcga
tttatggacagagttagatcatgctttgttgattcatgccaaactacgcctagacatttggagatcttggcaaa
atattgcggtccagaaattcgagcaattggcgaatggaaatgtttcgattccgttaatgaaatgggtttcctat
tggttgcgttatcctccaagccttgacaaccaatcagcaaccgatctgctttcaccttggctagttgatcgtaa
tggctacctttcaattaaactcccggaccgagttatccgaagctggcggaaggagattaactcattcagatggt
tactctcacctgatttaatggtaaaaggaaaatttgagactaatcgtttcaatgtagcatccatcgtgataggg
gaattacatcacggttttaccgccgatggttggatgtttgaatttcatcctgacaaaaaagaaatcaatcgaat
agtccggcaagctatgccagaaaagaatccaagctttacttgggctaattggatctttcaaaggaaaatgaaaa
gtactccgcaagagtacccaggtgtatcagtaaagcttacaggtcattctgaatacccagatgagaagtcattc
tctctatatgacttggaggttagaaggttaggcgataaagtggctgtttttttacaaggaactgataatgctct
aatgttttatcccccctgcctatggatttaacgaaacaagcttttttcccttttgcacttttctgttcacctatga
tggaaccttcatccaattactcaaaagggagacacaaagcaataaagttaggaaatgtcacattaattagggaa
cattgggttttcgagccgacagattggggagaagtccatgcatcacttgatggatttcaacaaatggcgtgttt
acaagagattaaggttaaatatcaattaccgcaagtaggatacttaagattttctcaagagctaaaacctgttt
ggtttgactttaataatccatttgtgtcgatttgtttttcaatatggctaaaaaagcagaatggtttacgttt
tccagaatggacccaggggtggaggatttatttctgaaggatgatcgaggccattattgttgtgaattgcgaac
ctttgcttatcggcaagagataacttccaaagatatgctcatgggttaactcgagcaccaccaccaccaccact
gagatccggctgctaacaaagcccgaaaggaagctgagttggctgctgccaccgctgagcaataactagcataa
ccccttggggcctctaaacgggtcttgaggggttttttgctgaaaggaggaactatatccggattggcgaatgg
gacgcgccctgtagcggcgcattaagcgcggcgggtgtggtggttacgcgcagcgtgaccgctacacttgccag
cgccctagcgcccgctcctttcgctttcttcccttcctttctcgccacgttcgccggctttccccgtcaagctc
taaatcgggggctccctttagggttccgatttagtgctttacggcacctcgaccccaaaaaacttgattagggt
gatggttcacgtagtgggccatcgccctgatagacggtttttcgccctttgacgttggagtccacgttctttaa
tagtggactcttgttccaaactggaacaacactcaaccctatctcggtctattcttttgatttataagggattt
tgccgatttcggcctattggttaaaaaatgagctgatttaacaaaaatttaacgcgaattttaacaaaatatta
acgcttacaatttaggtggcacttttcggggaaatgtgcgcggaacccctatttgtttatttttctaaatacat
tcaaatatgtatccgctcatgaattaattcttagaaaaactcatcgagcatcaaatgaaactgcaatttattca
tatcaggattatcaataccatatttttgaaaaagccgtttctgtaatgaaggagaaaactcaccgaggcagttc
cataggatggcaagatcctggtatcggtctgcgattccgactcgtccaacatcaatacaacctattaatttccc
ctcgtcaaaaataaggttatcaagtgagaaatcaccatgagtgacgactgaatccggtgagaatggcaaaagtt
tatgcatttctttccagacttgttcaacaggccagccattacgctcgtcatcaaaatcactcgcatcaaccaaa
ccgttattcattcgtgattgcgcctgagcgagacgaaatacgcgatcgctgttaaaaggacaattacaaacagg
aatcgaatgcaaccggcgcaggaacactgccagcgcatcaacaatattttcacctgaatcaggatattcttcta
atacctggaatgctgttttcccggggatcgcagtggtgagtaaccatgcatcatcaggagtacggataaaatgc
ttgatggtcggaagaggcataaattccgtcagccagtttagtctgaccatctcatctgtaacatcattggcaac
gctacctttgccatgtttcagaaacaactctggcgcatcgggcttcccatacaatcgatagattgtcgcacctg

attgcccgacattatcgcgagcccatttatacccatataaatcagcatccatgttggaatttaatcgcggccta
gagcaagacgtttcccgttgaatatggctcataacaccccttgtattactgtttatgtaagcagacagttttat
tgttcatgaccaaaatcccttaacgtgagttttcgttccactgagcgtcagaccccgtagaaaagatcaaagga
tcttcttgagatcctttttttctgcgcgtaatctgctgcttgcaaacaaaaaaaccaccgctaccagcggtggt
ttgtttgccggatcaagagctaccaactctttttccgaaggtaactggcttcagcagagcgcagataccaaata
ctgtccttctagtgtagccgtagttaggccaccacttcaagaactctgtagcaccgcctacatacctcgctctg
ctaatcctgttaccagtggctgctgccagtggcgataagtcgtgtcttaccgggttggactcaagacgatagtt
accggataaggcgcagcggtcgggctgaacggggggttcgtgcacacagcccagcttggagcgaacgacctaca
ccgaactgagatacctacagcgtgagctatgagaaagcgccacgcttcccgaagggagaaaggcggacaggtat
ccggtaagcggcagggtcggaacaggagagcgcacgagggagcttccagggggaaacgcctggtatctttatag
tcctgtcgggtttcgccacctctgacttgagcgtcgatttttgtgatgctcgtcaggggggcggagcctatgga
aaaacgccagcaacgcggcctttttacggttcctggccttttgctggccttttgctcacatgttctttcctgcg
ttatcccctgattctgtggataaccgtattaccgcctttgagtgagctgataccgctcgccgcagccgaacgac
cgagcgcagcgagtcagtgagcgaggaagcggaagagcgcctgatgcggtattttctccttacgcatctgtgcg
gtatttcacaccgcaatggtgcactctcagtacaatctgctctgatgccgcatagttaagccagtatacactcc
gctatcgctacgtgactgggtcatggctgcgccccgacacccgccaacacccgctgacgcgccctgacgggctt
gtctgctcccggcatccgcttacagacaagctgtgaccgtctccgggagctgcatgtgtcagaggttttcaccg
tcatcaccgaaacgcgcgaggcagctgcggtaaagctcatcagcgtggtcgtgaagcgattcacagatgtctgc
ctgttcatccgcgtccagctcgttgagtttctccagaagcgttaatgtctggcttctgataaagcgggccatgt
taagggcggttttttcctgtttggtcactgatgcctccgtgtaagggggatttctgttcatgggggtaatgata
ccgatgaaacgagagaggatgctcacgatacgggttactgatgatgaacatgcccggttactggaacgttgtga
gggtaaacaactggcggtatggatgcggcgggaccagagaaaaatcactcagggtcaatgccagcgcttcgtta
atacagatgtaggtgttccacagggtagccagcagcatcctgcgatgcagatccggaacataatggtgcagggc
gctgacttccgcgtttccagactttacgaaacacggaaaccgaagaccattcatgttgttgctcaggtcgcaga
cgttttgcagcagcagtcgcttcacgttcgctcgcgtatcggtgattcattctgctaaccagtaaggcaacccc
gccagcctagccgggtcctcaacgacaggagcacgatcatgcgcacccgtggggccgccatgccggcgataatg
gcctgcttctcgccgaaacgtttggtggcgggaccagtgacgaaggcttgagcgagggcgtgcaagattccgaa
taccgcaagcgacaggccgatcatcgtcgcgctccagcgaaagcggtcctcgccgaaaatgacccagagcgctg
ccggcacctgtcctacgagttgcatgataaagaagacagtcataagtgcggcgacgatagtcatgccccgcgcc
caccggaaggagctgactgggttgaaggctctcaagggcatcggtcgagatcccggtgcctaatgagtgagcta
acttacattaattgcgttgcgctcactgcccgctttccagtcgggaaacctgtcgtgccagctgcattaatgaa
tcggccaacgcgcggggagaggcggtttgcgtattgggcgccagggtggtttttcttttcaccagtgagacggg
caacagctgattgcccttcaccgcctggccctgagagagttgcagcaagcggtccacgctggtttgccccagca
ggcgaaaatcctgtttgatggtggttaacggcgggatataacatgagctgtcttcggtatcgtcgtatcccact
accgagatatccgcaccaacgcgcagcccggactcggtaatggcgcgcattgcgcccagcgccatctgatcgtt
ggcaaccagcatcgcagtgggaacgatgccctcattcagcatttgcatggtttgttgaaaaccggacatggcac
tccagtcgccttcccgttccgctatcggctgaatttgattgcgagtgagatatttatgccagccagccagacgc
agacgcgccgagacagaacttaatgggcccgctaacagcgcgatttgctggtgacccaatgcgaccagatgctc
cacgcccagtcgcgtaccgtcttcatgggagaaaataatactgttgatgggtgtctggtcagagacatcaagaa
ataacgccggaacattagtgcaggcagcttccacagcaatggcatcctggtcatccagcggatagttaatgatc
agcccactgacgcgttgcgcgagaagattgtgcaccgccgctttacaggcttcgacgccgcttcgttctaccat
cgacaccaccacgctggcacccagttgatcggcgcgagatttaatcgccgcgacaatttgcgacggcgcgtgca
gggccagactggaggtggcaacgccaatcagcaacgactgtttgcccgccagttgttgtgccacgcggttggga
atgtaattcagctccgccatcgccgcttccactttttcccgcgttttcgcagaaacgtggctggcctggttcac
cacgcgggaaacggtctgataagagacaccggcatactctgcgacatcgtataacgttactggtttcacattca
ccaccctgaattgactctcttccgggcgctatcatgccataccgcgaaaggttttgcgccattcgatggtgtcc
gggatctcgacgctctcccttatgcgactcctgcattaggaagcagcccagtagtaggttgaggccgttgagca
ccgccgccgcaaggaatggtgcatgcaaggagatggcgcccaacagtcccccggccacggggcctgccaccata
cccacgccgaaacaagcgctcatgagcccgaagtggcgagcccgatcttccccatcggtgatgtcggcgatata
ggcgccagcaaccgcacctgtggcgccggtgatgccggccacgatgcgtccggcgtagaggatcgagatctcga
tcccgcgaaat

**Table S3.** Sequences of tRNA molecules and the corresponding primers used for assembling templates for *in vitro* transcription. All sequences are provided 5' to 3' (left to right). Lowercase m indicates 2' O-methylation of the following residue.

| RNAs | Sequence |
|---|---|
| Dinitro flexizyme (dFx) | GGAUCGAAAGAUUUCCGCAUCCCCGAAAGGGUACAUGGCGUUAGGU |
| Enhanced flexizyme (eFx) | GGAUCGAAAGAUUUCCGCGGCCCCGAAAGGGGAUUAGCGUUAGGU |
| *E. coli* $tRNA^{Ala}$ (UGC) | GGGGCUAUAGCUCAGCUGGGAGAGCGCCUGCUUUGCACGCAGGAGGUCUGCGGUUCGAUCCCGCAUAGCUCCACCA |
| *P. syringae* $tRNA^{Cys}$ (GCA) | GGCCGAGUAGCAAAAUGGUUAUGCAGCGGAUUGCAAAUCCGCCUACGCCGGUUCGAUUCCGACCUCGGCCUCCA |
| E. coli $tRNA^{Trp}$ | AGGGGCGUAGUUCAAUUGGUAGAGCACCGGUCUCCAAAACCGGGUGUUGGGAGUUCGAGUCUCUCCGCCCCUGCCA |
| *T. bispora* $tRNA^{Glu}$ (CUC) | GGUCCCGUCGUCUAGAGGCCUAGGACGCCGCCCUCUCAAGGCGGUAACGGCGGUUCGAAUCCGCUCGGGACUACCA |
| *P. syringae* $tRNA^{Cys}$ (GCA) (U71A) | GGCCGAGUAGCAAAAUGGUUAUGCAGCGGAUUGCAAAUCCGCCUACGCCGGUUCGAUUCCGACCUCGGCCACCA |
| *P. syringae* $tRNA^{Cys}$ (GCA) (C34G/A35C/U71A) | GGCCGAGUAGCAAAAUGGUUAUGCAGCGGAUUGGCAAUCCGCCUACGCCGGUUCGAUUCCGACCUCGGCCACCA |
| *E. coli* $tRNA^{Ala}$ (UGC) grafted with *E. coli* $tRNA^{Trp}$ Anticodon Arm | GGGGCUAUAGCUCAGCUGGGAGAGCGCCGGUCUCCAAAACCGGAGGUCUGCGGUUCGAUCCCGCAUAGCUCCACCA |
| *E. coli* $tRNA^{Ala}$ (UGC) grafted with *P. syringae* $tRNA^{Cys}$ Anticodon Arm | GGGGCUAUAGCUCAGCUGGGAGAGCGGCGGAUUGCAAAUCCGCAGGUCUGCGGUUCGAUCCCGCAUAGCUCCACCA |
| *E. coli* $tRNA^{Ala}$ (UGC) grafted with *P. syringae* $tRNA^{Cys}$ D-Arm | GGGGCUAUAGCAAAAUGGUUAUGCACCUGCUUUGCACGCAGGAGGUCUGCGGUUCGAUCCCGCAUAGCUCCACCA |
| *E. coli* $tRNA^{Ala}$ (UGC) grafted with *P. syringae* $tRNA^{Cys}$ T-Stem | GGGGCUAUAGCUCAGCUGGGAGAGCGCCUGCUUUGCACGCAGGAGGUCGCCGGUUCGAUUCCGACUAGCUCCACCA |
| *E. coli* $tRNA^{Gly}$ (GCC) | GCGGGAAUAGCUCAGUUGGUAGAGCACGACCUUGCCAAGGUCGGGGUCGCGAGUUCGAGUCUCGUUUCCCGCUCCA |

| RNAs | Forward Primer Sequence | Reverse Primer Sequence |
|---|---|---|
| Dinitro flexizyme (dFx) | GGCGTAATACGACTCACTATAGGATCGAAAGATTTCCG | mAmCCTAACGCCATGTACCCTTTCGGGGATGCGGAAATCTTTCGATC |
| Enhanced flexizyme (eFx) | GGCGTAATACGACTCACTATAGGATCGAAAGATTTCCGCG | mAmCCTAACGCTAATCCCCTTTCGGGGCCGCGGAAATCTTTCG |

| | | |
|---|---|---|
| *E. coli* tRNA$^{Ala}$ (UGC) | AATTCCTGCAGTAATACGACTCACTATAGGGGCTATAGCTCAGCTGGGAGAGCGCCTGC | mUmGGTGGAGCTATGCGGGATCGAACCGCAGACCTCCTGCGTGCAAAGCAGGCGCTCTCCCAG |
| *P. syringae* tRNA$^{Cys}$ (GCA) | AATTCCTGCAGTAATACGACTCACTATAGGCCGAGTAGCAAAATGGTTATGCAGC | mUmGGAGGCCGAGGTCGGAATCGAACCGGCGTAGGCGGATTTGCAATCCGCTGCATAACC |
| E. coli tRNA$^{Trp}$ | AATTCCTGCAGTAATACGACTCACTATAAGGGGCGTAGTTCAATTGGTAGAGCACCGGTC | mUmGGCAGGGGCGGAGAGACTCGAACTCCCAACACCCGGTTTTGGAGACCGGTGCTCTACC |
| *T. bispora* tRNA$^{Glu}$ (CUC) | AATTCCTGCAGTAATACGACTCACTATAGGTCCCGTCGTCTAGAGGCCTAGGACGC | mUmGGTAGTCCCGAGCGGATTCGAACCGCCGTTACCGCCTTGAGAGGGCGGCGTCCTAGG |
| *P. syringae* tRNA$^{Cys}$ (GCA) (U71A) | AATTCCTGCAGTAATACGACTCACTATAGGCCGAGTAGCAAAATGGTTATGCAGC | mUmGGTGGCCGAGGTCGGAATCGAACCGGCGTAGGCGGATTTGCAATCCGCTGCATAACC |
| *P. syringae* tRNA$^{Cys}$ (GCA) (C34G/A35C/U71A) | AATTCCTGCAGTAATACGACTCACTATAGGCCGAGTAGCAAAATGGTTATGCAGC | mUmGGTGGCCGAGGTCGGAATCGAACCGGCGTAGGCGGATTGCCAATCCGCTGCATAACC |
| *E. coli* tRNA$^{Ala}$ (UGC) grafted with *E. coli* tRNA$^{Trp}$ Anticodon Arm | AATTCCTGCAGTAATACGACTCACTATAGGGGCTATAGCTCAGCTGGGAGAGCGCCGGT | mUmGGTGGAGCTATGCGGGATCGAACCGCAGACCTCCGGTTTTGGAGACCGGCGCTCTCCCAG |
| *E. coli* tRNA$^{Ala}$ (UGC) grafted with *P. syringae* tRNA$^{Cys}$ Anticodon Arm | AATTCCTGCAGTAATACGACTCACTATAGGGGCTATAGCTCAGCTGGGAGAGCGGCGGA | mUmGGTGGAGCTATGCGGGATCGAACCGCAGACCTGCGGATTTGCAATCCGCCGCTCTCCCAG |
| *E. coli* tRNA$^{Ala}$ (UGC) grafted with *P. syringae* tRNA$^{Cys}$ D-Arm | AATTCCTGCAGTAATACGACTCACTATAGGGGCTATAGCAAAATGGTTATGCACCTGCT | mUmGGTGGAGCTATGCGGGATCGAACCGCAGACCTCCTGCGTGCAAAGCAGGTGCATAACCAT |
| *E. coli* tRNA$^{Ala}$ (UGC) grafted with *P. syringae* tRNA$^{Cys}$ T-Stem | AATTCCTGCAGTAATACGACTCACTATAGGGGCTATAGCTCAGCTGGGAGAGCGCCTGC | mUmGGTGGAGCTAGTCGGAATCGAACCGGCGACCTCCTGCGTGCAAAGCAGGCGCTCTCCCAG |
| *E. coli* tRNA$^{Gly}$ (GCC) | AATTCCTGCAGTAATACGACTCACTATAGCGGGAATAGCTCAGTTGGTAGAGCACGACC | mUmGGAGCGGGAAACGAGACTCGAACTCGCGACCCCGACCTTGGCAAGGTCGTGCTCTACCAA |

**Table S4.** Sequences of the Cy3-labeled primers used for the $NaIO_4$-RNA extension assays. All sequences are provided 5' to 3' (left to right). Cy3 was incorporated at the 3'-OH of these primers.

| RNAs | Cy3 Primer Sequence |
|---|---|
| *E. coli* $tRNA^{Ala}$ (UGC) | TTAAAAAAAAAAAAAAAAAAAAAAAAAAAAATGGTGGAGCTATGC GGGATCGAACCGCAGACC |
| *P. syringae* $tRNA^{Cys}$ (GCA) | TTAAAAAAAAAAAAAAAAAAAAAAAAAAAAATGGAGGCCGAGGTC GGAATCGAACCGGCGTAG |
| E. coli $tRNA^{Trp}$ | TTAAAAAAAAAAAAAAAAAAAAAAAAAAAAATGGCAGGGGCGGAG AGACTCGAACTCCCAACA |
| *T. bispora* $tRNA^{Glu}$ (CUC) | TTAAAAAAAAAAAAAAAAAAAAAAAAAAAAATGGTAGTCCCGAGC GGATTCGAACCGCCGTTA |
| *P. syringae* $tRNA^{Cys}$ (GCA) (U71A) | TTAAAAAAAAAAAAAAAAAAAAAAAAAAAAATGGAGGCCGAGGTC GGAATCGAACCGGCGTAG |
| *P. syringae* $tRNA^{Cys}$ (GCA) (C34G/A35C/U71A) | TTAAAAAAAAAAAAAAAAAAAAAAAAAAAAATGGTGGCCGAGGTC GGAATCGAACCGGCGTAG |
| *E. coli* $tRNA^{Ala}$ (UGC) grafted with *E. coli* $tRNA^{Trp}$ Anticodon Arm | TTAAAAAAAAAAAAAAAAAAAAAAAAAAAAATGGTGGAGCTATGC GGGATCGAACCGCAGACC |
| *E. coli* $tRNA^{Ala}$ (UGC) grafted with *P. syringae* $tRNA^{Cys}$ Anticodon Arm | TTAAAAAAAAAAAAAAAAAAAAAAAAAAAAATGGTGGAGCTATGC GGGATCGAACCGCAGACC |
| *E. coli* $tRNA^{Ala}$ (UGC) grafted with *P. syringae* $tRNA^{Cys}$ D-Arm | TTAAAAAAAAAAAAAAAAAAAAAAAAAAAAATGGTGGAGCTATGC GGGATCGAACCGCAGACC |
| *E. coli* $tRNA^{Ala}$ (UGC) grafted with *P. syringae* $tRNA^{Cys}$ T-Stem | TTAAAAAAAAAAAAAAAAATGGTGGAGCTAGTCGGAATCGAACCG GCGACCTCCTGCGTGC |
| *E. coli* $tRNA^{Gly}$ (GCC) | TTAAAAAAAAAAAAAAAAAAAAAAAAAAAAATGGAGCGGGAAACG AGACTCGAACTCGCGACC |

**Table S5.** Activated amino acids used in this study as substrates for flexizymes along with summary of tested flexizyme conditions. CBT = 4-chlorobenzyl thioester, DBE = 3,5-dinitrobenzyl ester, CME = cyanomethyl ester.

| **Amino Acid, Concentration** | **Flexizyme** | **pH** | **[$Mg^{2+}$]** | **Reaction time** | **% DMSO (*v/v*)** |
|---|---|---|---|---|---|
| Ala-DBE, 5 mM | dFx | 7.5 | 100 mM | 6 h | 20 |
| Glu-DBE, 5 mM | dFx | 7.5 | 100 mM | 6 h | 20 |
| Gln-DBE, 5 mM | dFx | 7.5 | 100 mM | 6 h | 20 |
| (N-Me-Ala)-DBE, 5 mM | dFx | 7.5 | 100 mM | 6 h | 20 |
| (D-Ala)-DBE, 5 mM | dFx | 7.5 | 100 mM | 6 h | 20 |
| Aib-DBE, 5 mM | dFx | 7.5 | 600 mM | 2.5 h | 20 |
| ThioGly-DBE, 5 mM | dFx | 9.0 | 600 mM | 6 h | 40 |
| Phe-CME, 5 mM | eFx | 7.5 | 600 mM | 2.5 h | 20 |
| Trp-CME, 5 mM | eFx | 7.5 | 600 mM | 2.5 h | 20 |
| Ala-CBT, 5 mM | eFx | 7.5 | 600 mM | 2.5 h | 20 |
| Gly-CBT, 5 mM | eFx | 7.5 | 600 mM | 2.5 h | 20 |
| (β-Ala)-CBT, 5 mM | eFx | 7.5 | 600 mM | 2.5 h | 20 |
| Lac-CBT, 5 mM | eFx | 7.5 | 482 mM | 2.5 h | 32 |

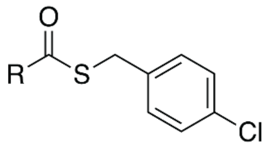

Acid charged with CBT

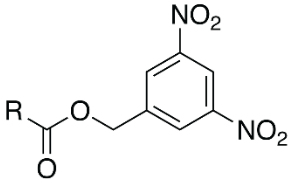

Acid charged with DBE

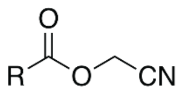

Acid charged with CME

**Table S6.** Calculated and observed *m/z* values for products and starting materials in this study.

| Peptide | Calculated *m/z* (z =2) | Observed *m/z* | Error (ppm) |
|---|---|---|---|
| Δ19BhaA | 1137.0753 | 1137.0764 | 0.97 |
| Δ19BhaA-Ala | 1172.5939 | 1172.5972 | 2.81 |
| Δ19BhaA-Lac | 1173.0859 | 1173.0862 | 0.28 |
| Δ19BhaA-Gln | 1201.1046 | 1201.1088 | 3.49 |
| Δ19BhaA-Glu | 1201.5966 | 1201.5970 | 0.32 |
| Δ19BhaA-(N-Me-Ala) | 1179.6017 | No product observed | |
| Δ19BhaA-Trp | 1230.1150 | 1230.1165 | 1.24 |
| Δ19BhaA-ThioGly | 1174.0667 | 1174.0672 | 0.43 |
| Δ19BhaA-Cysteamine | 1166.5851 | 1166.5861 | 0.88 |
| Δ19BhaA-Gly | 1165.5861 | 1165.5866 | 0.47 |
| Δ19BhaA-Phe | 1210.6095 | 1210.6115 | 1.63 |
| Δ19BhaA-(β-Ala) | 1172.5939 | 1172.5970 | 2.64 |
| Δ19BhaA-(D-Ala) | 1172.5939 | 1172.5963 | 2.05 |
| Δ19BhaA-Aib | 1179.6017 | No product observed | |
